# Reduced reproductive success is associated with selective constraint on human genes

**DOI:** 10.1101/2020.05.26.116111

**Authors:** Eugene J. Gardner, Matthew D. C. Neville, Kaitlin E. Samocha, Kieron Barclay, Martin Kolk, Mari E. K. Niemi, George Kirov, Hilary C. Martin, Matthew E. Hurles

**Affiliations:** Wellcome Sanger Institute, Wellcome Genome Campus, Cambridge, Hinxton, United Kingdom; Max Planck Institute for Demographic Research, Rostock, Germany; Demography Unit, Department of Sociology, Stockholm University, Stockholm, Sweden; Swedish Collegium for Advanced Study, Uppsala, Sweden; Division of Psychological Medicine and Clinical Neurosciences, School of Medicine, Cardiff University, Cardiff, UK

## Abstract

Genome-wide sequencing of human populations has revealed substantial variation among genes in the intensity of purifying selection acting on damaging genetic variants^1^. While genes under the strongest selective constraint are highly enriched for associations with Mendelian disorders, most of these genes are not associated with disease and therefore the nature of the selection acting on them is not known^2^. Here we show that genetic variants that damage these genes are associated with markedly reduced reproductive success, primarily due to increased childlessness, with a stronger effect in males than in females. We present evidence that increased childlessness is likely mediated by genetically associated cognitive and behavioural traits, which may mean male carriers are less likely to find reproductive partners. This reduction in reproductive success may account for 20% of purifying selection against heterozygous variants that ablate protein-coding genes. While this genetic association could only account for a very minor fraction of the overall likelihood of being childless (less than 1%), especially when compared to more influential sociodemographic factors, it may influence how genes evolve over time.

## Background

The most damaging genetic variants, gene deletions and protein-truncating variants (PTVs), are removed from the population by selection with strength that varies substantially from gene to gene^1,3^. The strength of selection against heterozygous PTVs has been estimated by the selection coefficient, s_het_, which exhibits a continuous spectrum across human genes^2,4^, although most attention has been focused on a subset of ~3,000 genes with the highest probability of Loss-of-function Intolerance (pLI)^1^.

The selection pressures acting on these most selectively constrained genes have not been fully characterised, but, *a priori*, could include natural selection against variants increasing pre-reproductive mortality or decreasing fertility, and sexual selection acting on mate choice or intra-sexual competition^5^. Gene deletions and PTVs in these genes have been shown to be associated with lower educational attainment^6,7^ and general intelligence^8^, as well as increased risk of intellectual disability, and some psychiatric disorders^9^. Moreover, these constrained genes are strongly enriched for associations with dominant early-onset Mendelian diseases (including neurodevelopmental disorders), many of which are associated with increased pre-reproductive mortality, indicating that natural selection likely makes a substantive contribution to this selective constraint. However, the majority (65%) of highly constrained genes (pLI ≥ 0.9) have not yet been associated with a Mendelian disease. This raises the fundamental question of whether natural selection represents the sole mechanism imposing this form of selective constraint on human genes.

### Genetic association testing

To explore the nature of selection acting on these genes we identified PTVs and genic deletions in the UK Biobank^10^ comprising largely post-reproductive individuals (median age at recruitment: 58 years, range: 39-73 years, birth years: 1934-1970; Supplementary Figure 1), and investigated the association with reproductive success. We called large copy number variants (deletions and duplications) from SNP genotyping array data on 340,925 unrelated participants of European genetic ancestry (Supplementary Figure 2), and identified PTVs from exome sequencing among a subset of 139,477 participants (Supplementary Figure 3)^11^. For each participant, we then calculated the cumulative burden of private (only observed in one individual), heterozygous genic deletions and PTVs by combining s_het_ selection coefficients of each autosomal gene impacted by these variants (under the assumption that fitness is multiplicative, see Methods), which we term their s_het_ burden. The distribution of s_het_ burden was statistically indistinguishable between males and females: for participants with only genic deletion data available, 0.56% and 0.55% respectively had an s_het_ burden ≥ 0.15 (Kolmogorov-Smirnov p=1.00; Figure 1C), and for participants with both genic deletion and PTV data available the analogous proportions were 4.60% and 4.59%, respectively (Kolmogorov-Smirnov p=0.52; Figure 1D).

**Figure 1.**
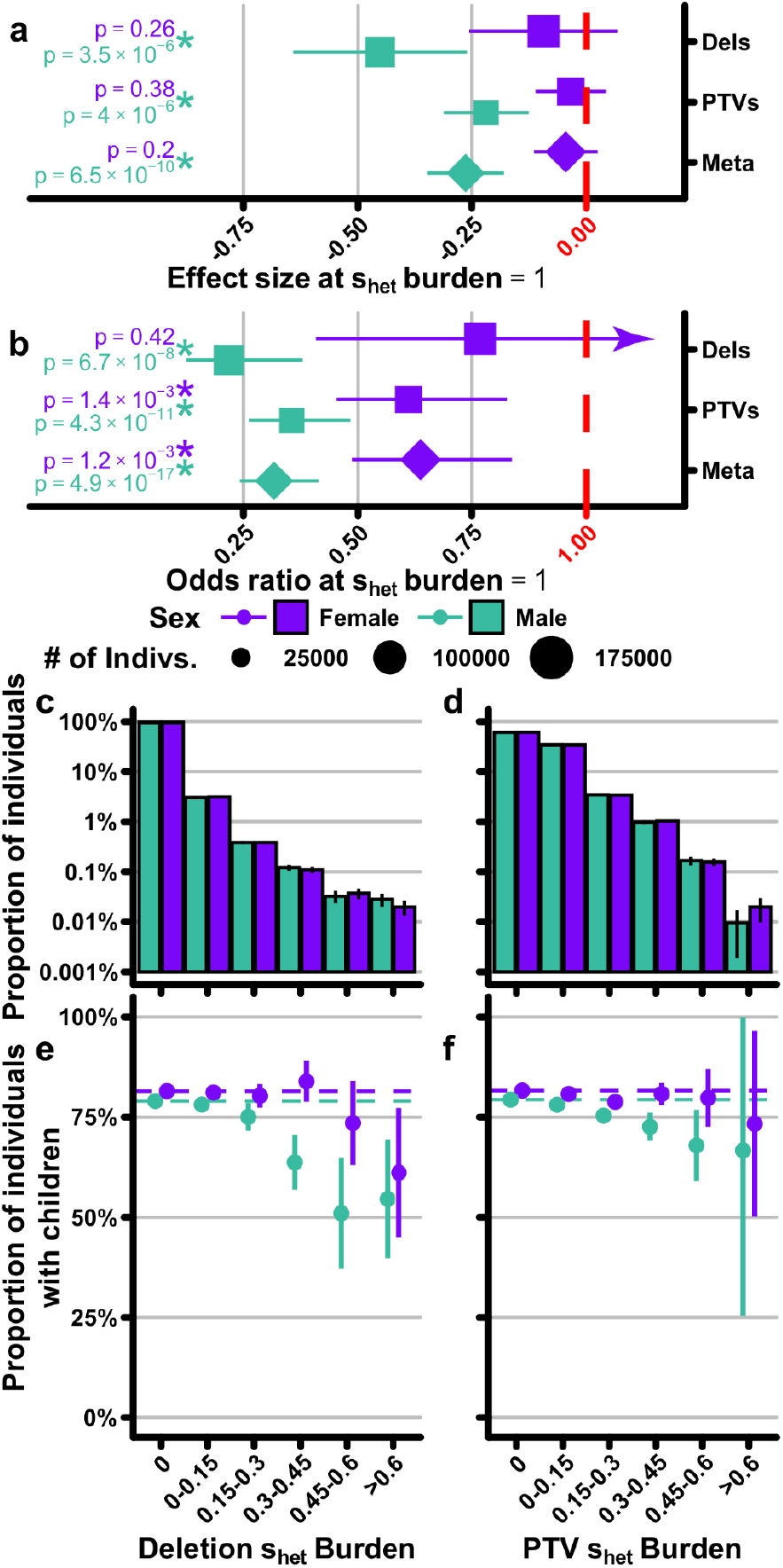
Differences in male and female reproductive success as a function of cumulative rare deleterious genetic variation. (A, B) Effect size/odds ratio estimates for the association of cumulative deleterious variation for deletions (dels), SNV and InDel Protein Truncating Variants (PTVs), and a combined meta-analysis with number of children (Meta) (A) and childlessness (B) separated for males (jade) and females (violet). Number of individuals included in each analysis is indicated by the size of the point. Asterisks indicate significant associations after Bonferroni correction for 20 tests (p < 2.5×10^-3^; Methods). The arrow in panel (B) indicates the confidence interval stretches beyond the limits of the y-axis. (C; D) Proportion of individuals in 0.15 s_het_ bins for deletions (C) and PTVs (D). (E; F) Percentage of individuals with children in bins based on s_het_ burden for deletions (E) and PTVs (F). Error bars for panels C-F are 95% confidence intervals calculated on the population proportion.

We assessed the relationship between s_het_ burden and number of children, using a linear regression correcting for age, genetic ancestry, and birth cohort (Methods; Figure 1A; Supplementary Table 1). We observed that an s_het_ burden of 1 (the highest possible burden) is associated with a decrease in the overall total number of children for males (0.26 fewer children [95% CI 0.18-0.34], p=6.5×10^-10^) but not females (0.05 fewer children [95% CI −0.02-0.11], p=0.20) when combining results from deletion and PTV-based analyses.

To determine if the observed association of s_het_ burden with reproductive success was primarily linked to an increased likelihood of remaining childless, we performed two analyses. Firstly, we evaluated childlessness using logistic regression and again observed a striking sex difference in participants’ probability of having any children given their s_het_ burden, for both PTVs and genic deletions (Figure 1B). Combining the analyses of genic deletions and PTVs, we found that an s_het_ burden of 1 is associated with a lower probability of males having any children (OR=0.32 [95% CI 0.24-0.41], p=4.9×10^-17^) much more than females (OR=0.64 [95% CI 0.49-0.84], p=1.2×10^-3^). We also observed that private duplications and likely damaging private missense variants exhibit similar but weaker associations with childlessness (Extended Data Figure 1). Secondly, if we remove childless individuals from the analysis, s_het_ burden ceases to have a significant correlation with the number of offspring (Extended Data Figure 2). This is supported by the observation that, when we stratify individuals with children by the number of children, the s_het_ distribution does not vary and confirms that the observed association with reproductive success relates primarily to increased childlessness (Supplementary Figure 4). This observation is consistent with studies that have associated demographic factors with reproductive success, which have also observed that associations are weakened or disappear when childless individuals are removed from the analysis (e.g. Barthold et al.^12^).

We also considered whether ascertainment biases or differences in fertility between the UK Biobank sample and the UK population as a whole could affect these results. As UK Biobank participants included in these analyses are enriched for females (54%), the observed sex difference is not due to having greater statistical power to detect an effect on reproductive success in males. Likewise, fertility rates between the UK Biobank population and the UK population as a whole are highly similar; the average total fertility rate in the UK from 1983-2000 (where data are available for both males and females), when UK Biobank participants would have been reproductively active, was 1.78±0.07 s.e.m. for males^13^ and 1.76±0.05 s.e.m. for females^14^, which is very similar to that observed among UK Biobank participants (average number of children for males = 1.77; females = 1.80).

We observed a consistent sex difference in the association of s_het_ burden with childlessness when performing this analysis in different ways, including: (i) limiting analyses to carriers of private genic deletions or PTVs in the genes under most selective constraint (following thresholds set by their authors: pLI ≥ 0.9^1^ or s_het_ ≥ 0.15^2^; Supplementary Figure 5), (ii) extending analysis to more frequent, but still rare genic deletions and PTVs (Supplementary Figure 6), (iii) excluding genes known to be associated with a disease (male OR=0.33 [95% CI 0.24-0.46], p=4.1×10^-11^; female OR=0.68 [95% CI 0.49-0.94], nominal p=0.02; Methods), and (iv) restricting analysis to individuals in specific birth cohorts (Extended Data Figure 3).

### Evaluating different hypotheses

We evaluated three hypotheses that could account for the association with increased childlessness: (i) impaired fertility (e.g. inability to produce viable gametes), (ii) adverse health conditions, and (iii) cognitive and behavioural factors (which may be associated with decreased chances of finding a reproductive partner or increased voluntary childlessness). We observed that s_het_ burden is not significantly associated (after correcting for multiple testing) with an increased risk of male (OR=6.37 [95% CI 1.07-37.87], nominal p=0.04) or female infertility (OR=0.83 [95% CI 0.33-2.09], p=0.70) as defined based on combined health outcomes data for all UK Biobank participants (combined hospital episode statistics, primary care records, self-reported conditions, and death records). Three additional lines of evidence suggest impaired fertility is not the predominant cause of the sex-differential association between s_het_ burden and childlessness. First, when introducing male infertility status as a covariate in the association testing, we observed minimal reduction in the strength of the association between s_het_ burden and male childlessness (OR=0.32 [95% CI 0.24-0.42], p=6.2×10^-17^; Supplementary Table 2). Second, we observed minimal change in the association between s_het_ burden and male reproductive success after removing all 150 autosomal genes for which at least limited evidence exists of an association to male infertility (OR=0.32 [95% CI 0.24-0.42], p=1.1×10^-16^)^15^ or all 742 genes associated with male infertility in mice (OR=0.33 [95% CI 0.25-0.43], p=3.1×10^-15^)^16^. Finally, genes under the highest selective constraint (s_het_ ≥ 0.15) do not appear to be more highly expressed in testis, unlike the genes currently known to be associated with male infertility (Supplementary Figure 7). Together, these findings are consistent with a previous study that sought but did not find a widespread role for highly-penetrant dominant deletions in spermatogenic failure^17^.

Previous studies have shown that physical birth defects are associated with reduced reproductive success^18,19^. We assessed whether any adverse health conditions contributed to the association between s_het_ burden and childlessness. We independently tested 19,154 ICD-10 codes (from both hospital episode statistics and combined health outcomes data across four levels of the ICD-10 hierarchy; Methods) as a covariate in the association test of s_het_ burden with childlessness (Extended Data Figures 4, 5; Supplementary Figures 8, 9; Supplementary Table 2; Methods). We found that while many ICD-10 codes are associated with having children, in particular positive associations with male-specific codes for elective sterilisation and female-specific codes associated with pregnancy and childbirth (Extended Data Figure 5), correcting for any ICD-10 code had minimal impact on the strength of association between s_het_ burden and childlessness (Extended Data Figure 4; Supplementary Figures 8, 9). The ICD-10 code with the largest (but still modest) impact on the association of s_het_ burden and male childlessness was observed for ‘developmental disorders of scholastic skills’, although this only reduced the Odds Ratio of 0.317 to 0.325 (Extended Data Figure 4; Supplementary Table 2). In addition to diseases, clinically annotated ICD-10 codes are also available for a range of factors denoting health status and contact with health services. We noted that one of these also had a modest impact on the association of s_het_ burden and male childlessness when included as covariates in association testing (Extended Data Figure 4; Supplementary Table 2). This code relates to social environment problems (code Z60), with the association driven primarily by the status of living alone (subcode Z60.2). This code is also positively associated with s_het_ burden.

There is existing evidence, primarily based on examinations of western populations, that behavioural and cognitive traits are associated with reproductive success in a sex-differential manner. First, the reduced reproductive success associated with a range of psychiatric disorders is much more pronounced in males^20^. Second, personality traits associated with increased reproductive success differ between males and females, with increased extraversion in males but greater neuroticism in females being linked to increased reproductive success^21^. Third, although the most highly ranked characteristics in a reproductive partner are highly concordant between the sexes^22^, some mate preferences differ between the sexes, with males placing greater value on physical attractiveness and females valuing cues relating to earning potential^21–24^. Finally, low socioeconomic status and low educational attainment have been more strongly linked to increased childlessness in males than females across populations^25–28^. This has been hypothesized to be due to males of lower socioeconomic status finding it harder to find a partner^12,29^.

Highly constrained genes exhibit higher expression levels and broader expression across tissues than less constrained genes^1^. Ubiquitous expression potentially impedes insight into the underlying biology of selection. We generated s_het_ burden scores excluding genes expressed in 53 tissues characterised by GTEx^30^ and then used these scores to test the association with childlessness (Supplementary Figure 10). As expected due to the large number of genes excluded for each tissue, we observed a substantial decrease in statistical significance for all tissues in males and all but one tissue in females. However, overlapping confidence intervals on effect size estimates make identification of tissues most relevant for selection challenging. Nevertheless, we did observe that s_het_ was more strongly positively correlated with gene expression levels in all 13 brain regions than all 40 non-brain regions (including testis)^30^. When s_het_ values are weighted by the number of PTVs observed in UK Biobank (thus incorporating consideration of each gene’s contribution to s_het_ burden) expression in brain tissues remains more strongly correlated than any other tissue (Supplementary Figure 11). These analyses highlight the potential greater relevance of the brain in the biological processes underlying selection against PTVs in highly constrained genes.

Some of the observations about behavioural and cognitive associations with sex-differential reproductive success have been related to sexual selection by mate choice, in which one sex (typically female) tends to be more discriminating in their choice of reproductive partners. One theory posits that this is due to differential parental investment^31^; the sex that invests more in offspring (typically female) is more discriminating in its choice of reproductive partners, especially with regard to those partners’ potential to invest in offspring. Alternative theories have been proposed regarding the causes of sexual selection, including those that focus on disparities in gamete size (Darwin-Bateman paradigm^32,33^).

One prediction of the hypothesis that differential mate choice underpins the observation of a stronger male association of s_het_ burden with increased childlessness is that males with a high s_het_ burden should find it harder to find reproductive partners than females. We observed that UK Biobank participants with high s_het_ burden were significantly less likely to have reported currently living with a partner – consistent with the findings from ICD-10 codes – and that, like reproductive success, this association was significantly stronger in males than in females (Figure 2A). UK Biobank males currently living with a partner are also much more likely to have children (OR=5.80 [95% CI 5.64-5.97], p < 1×10^-100^; Extended Data Figure 6A). We note that the status of currently living with a partner is an imperfect proxy for partner status during peak reproductive years, but the latter information is not currently available in UK Biobank. We also found that s_het_ burden was significantly associated with a lower probability of ever having had sex for both male (OR=0.06 [95% CI 0.03-0.14], p=2.5×10^-11^) and female (OR=0.11 [95% CI 0.05-0.27], p=1.2×10^-6^) UK Biobank participants, without significant sex-difference (Figure 2B). Additionally, while same sex sexual behaviour is strongly associated with increased childlessness in UK Biobank (male OR=0.14 [95% CI 0.13-0.15], p<1×10^-100^; female OR=0.27 [95% CI 0.25-0.29], p<1×10^-100^; Extended Data Figure 6H), we observed no significant association after multiple-testing correction of s_het_ burden with the likelihood of having engaged in same sex sexual behaviour among males (OR=1.98 [95% CI 1.10-3.55], nominal p=0.02; Supplementary Figure 12) nor did we observe any change in the association between s_het_ burden and childlessness when excluding individuals who engaged in same-sex sexual behaviour from our primary model (OR=0.33 [95% CI 0.25-0.43], p=4.5×10^-15^).

**Figure 2.**
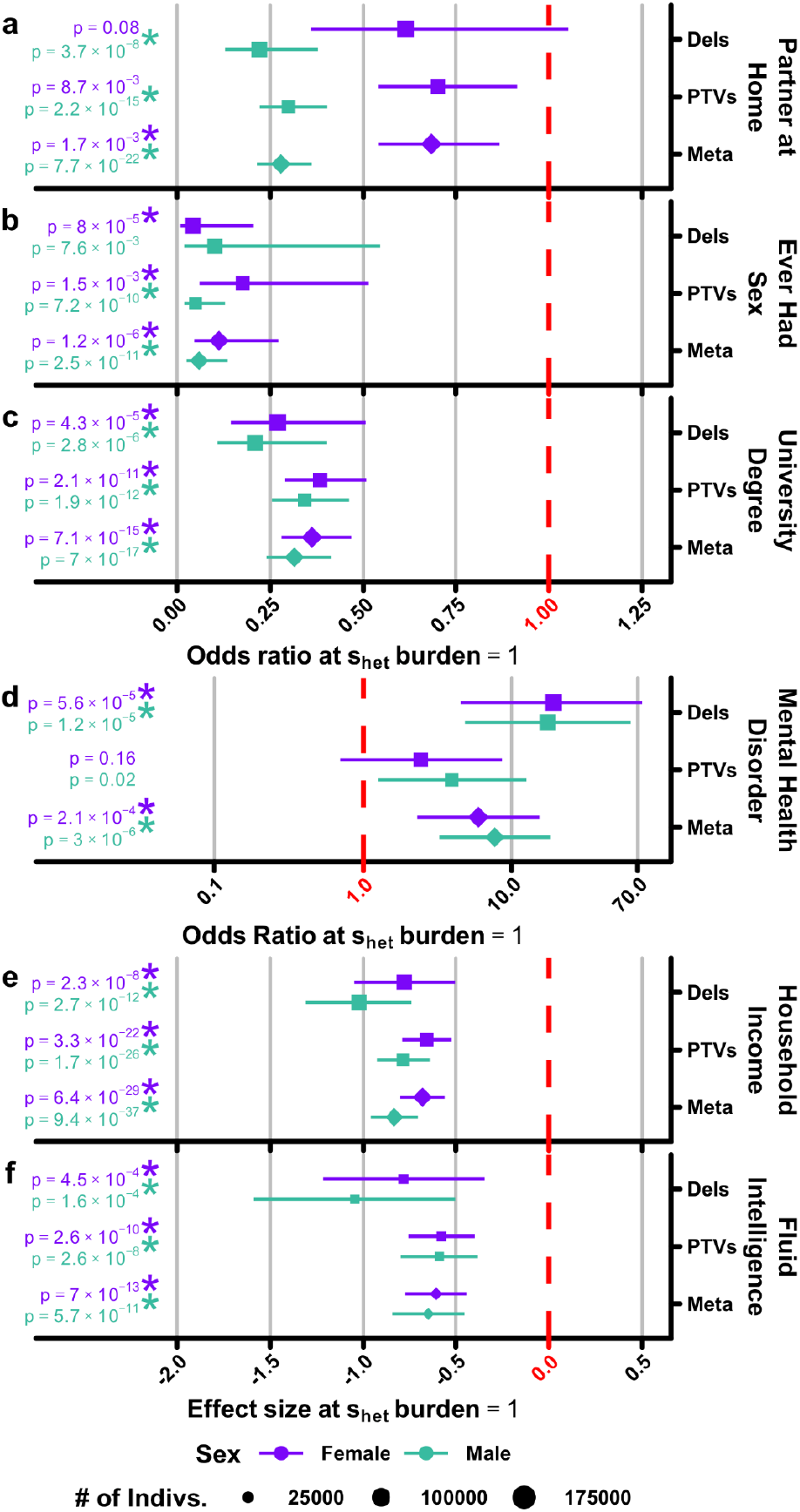
Association of s_het_ burden with traits known to be associated with reproductive success. Shown are similar plots to Figure 1A,B for six additional phenotypes: (A) having a partner at home, (B) ever having engaged in sexual intercourse, (C) educational attainment as measured by having a university degree, (D) household income (as measured by income bracket and corrected for having a partner at home; see methods), (E) fluid intelligence (in standard deviations), and (F) having a mental health disorder. For each trait, we tested using a logistic (A,B,C,D) or linear (E,F) model for the association of s_het_ burden with each phenotype shown above (methods). Note that the x-axis for plot (D) is in log rather than linear scale. Asterisks indicate significant associations after Bonferroni correction for 20 tests (p < 2.5×10^-3^; Methods).

While these findings are consistent with a role for the ability to find a partner in the association between s_het_ burden and reproductive success, this relationship need not be caused by sex differences in mate preferences. It could also result from sex differences in the severity of cognitive and behavioural traits that are associated with s_het_ burden, coupled to mate choice preferences that are not sex-differential. These mechanisms are not mutually exclusive; both could be contributing to an overall sex-differential reduction in reproductive success, albeit on different traits. We explored in UK Biobank whether the association of s_het_ burden on reproductive success might plausibly be mediated through some of the specific factors highlighted by the previous psychiatric, demographic and psychosocial research summarised above. Firstly we investigated the association of s_het_ burden with cognition as measured by fluid intelligence in 106,299 (49,631 males, 56,668 females) UK Biobank participants. We found that s_het_ burden was associated with significantly reduced fluid intelligence scores of males and females with similar effect sizes (Figure 2F). Increasing s_het_ burden is also associated with lower educational attainment (Figure 2C), lower household income (Figure 2E), and greater socioeconomic deprivation (Extended Data Figure 7), again with similar effect sizes in males and females.

To evaluate the potential link between lower cognitive performance and male reproductive success in a more quantitative manner in a less biased population sample, we extended previously published work relating the results of IQ tests taken by 95% of Swedish males during military conscription to their completed family size^34^. We estimated that lower scores in IQ tests could account for 11% [95% CI 8%-12%] of the reduced male reproductive success associated with high s_het_ burden (Supplementary Figure 13, Methods). We also note that, in the Swedish data, the association between reduced reproductive success and lower IQ scores was most pronounced in males with IQ<70 (Supplementary Figure 14)^34^; such individuals are likely proportionately fewer in UK Biobank relative to the general population.

Analysis of psychiatric disorders in UK Biobank is complicated by both recruitment bias away from more severe psychiatric disorders and incomplete data on participants^10,35–37^. The most comprehensive data are available on a subset of UK Biobank individuals from a mental health questionnaire for which participants were invited by email (n = 157,366)^36^. We observed that a high s_het_ burden was very strongly associated with not having an email address (male OR=0.30 [95% CI 0.24-0.39], p=4.1×10^-21^; female OR=0.45 [95% CI 0.36-0.57], p=1.4×10^-11^; Supplementary Figure 15), which likely explains why individuals with a high s_het_ burden were much less likely to complete the questionnaire (male OR=0.43 [95% CI 0.33-0.56], p=5.5×10^-10^; female OR=0.46 [95% CI 0.36-0.58], p=2.1×10^-10^; Supplementary Figure 15). Therefore we focused analyses of mental health disorders on the complete health outcomes data available for all participants. These data corroborate a previous finding^9^ that high s_het_ burden is associated with increased risk of psychiatric disorders previously linked with reduced reproductive success (intellectual disability, schizophrenia, autism, attention deficit hyperactivity disorder, and bipolar disorder; Figure 2D, Supplementary Figure 16)^20^. In accordance with prior literature on the association of psychiatric diagnoses with reproductive success^20^ and among UK Biobank participants, psychiatric disorders are associated with decreased probability of having any children in both males (OR=0.36 [95% CI 0.32-0.40], p=8.9×10^-78^) and females (OR=0.58 [95% CI 0.52-0.66], p=2.1×10^-18^), albeit with substantially stronger association in males (Extended Data Figure 6D). This finding accords with a previous study showing that copy number variants associated with increased risk of schizophrenia are also associated with disproportionately reduced reproductive success in males^38^. Removal of carriers of well-characterised neurodevelopmental disorder-associated copy number variants (n = 12,608 individuals; Methods), which includes schizophrenia-associated variants, does not significantly alter the association of s_het_ burden with reduced male reproductive success (OR=0.31 [95% CI 0.23-0.40], p=6.3×10^-17^).

We subsequently tested the association of s_het_ burden with childlessness excluding individuals with any evidence of a mental health disorder linked with reduced reproductive success (from hospital episode statistics, combined health outcomes data, or the mental health questionnaire). We observed very similar effect sizes to when analysing all individuals (male OR=0.33 [95% CI 0.25-0.43], p=1.6×10^-15^; female OR=0.63 [95% CI 0.48-0.83], p=1.1×10^-3^), suggesting that the association with childlessness is not predominantly driven by this subset of mental health disorders. We explored this further, using data external to UK Biobank that are less affected by the limitations described above. Using previous estimates of the increased risk of mental health disorders associated with PTVs in highly constrained genes^9^, and the reduced reproductive success associated with those disorders^20^, we estimated that these mental health disorders could account for 16% [8 - 35%] of the reduced male reproductive success associated with high s_het_ burden (Methods). Thus, in UK Biobank, both lower performance in tests of fluid intelligence and increased risk of psychiatric disorders likely account for only modest proportions of increased male childlessness associated with s_het_ burden.

We next used multiple regression to explore how much of the association between s_het_ burden and childlessness can be accounted for by the multiple correlated factors described above (where available for the entire cohort), namely: living with a partner, having had sex, having a mental health disorder associated with reduced reproductive success, having a university degree and having an infertility code in health records. Collectively, these factors can account for most of the association of s_het_ burden with childlessness in males (68%) but a smaller proportion in females (16%), as assessed by the difference in incremental Nagelkerke’s r^2^ of PTV s_het_ burden between models (Methods; Extended Data Figure 8A; Supplementary Figure 17). By far the biggest contribution comes from living with a partner and having had sex (Extended Data Figure 8A). Importantly, this analysis also showed that, despite the statistical significance of the association between s_het_ burden and childlessness, s_het_ burden accounts for less than 1% of the likelihood of being childless for both sexes.

### Contribution to selective constraint

Overall, we estimate that reduced reproductive success due to s_het_ burden potentially accounts for 20% [13-28%] (Extended Data Figure 8B; Extended Data Figure 9) of the total reduction in fitness expected due to purifying selection against PTVs as predicted by s_het_ (Methods)^4^, with this reduction in fitness being much stronger in males. We note that the total reduction in fitness predicted by s_het_ will include a substantive contribution from pre-reproductive mortality, which is not quantified here. We also note that current estimates of s_het_ are based on data from aggregated research cohorts, and may thus be biased upwards due to individuals with high s_het_ burden likely being under-represented; participation in research has previously been shown to be biased with respect to gender, socioeconomic status and genetic variation^39^. Therefore, PTVs within highly constrained genes may be present at higher frequencies in the general population than in research cohorts. This bias could result in the true value of s_het_ being lower than currently estimated, and, consequently, the contribution of reduced reproductive success to the overall reduction in fitness due to purifying selection being greater than estimated here.

These estimates of reproductive success and selection coefficients are reflective of a population at a particular point in time. The proportionate contribution of reduced reproductive success to the overall reduction in fitness associated with genic purifying selection is likely to change over time. Medical advances have altered the landscape of infertility and pre-reproductive mortality substantially. Moreover, childlessness is highly dynamic over time. Demographic data demonstrate that population-wide childlessness can double in just two decades, a trend that is readily apparent in UK Biobank (Supplementary Figure 1). We cannot discount that sex-differential sociodemographic factors, in addition to sex differences in trait severity and mate choices, could also be contributing to dampening the apparent association between s_het_ burden and childlessness in women. Higher educational attainment has been shown to be one of the factors most strongly positively associated with childlessness in females^27^. Indeed, when we correct the association between s_het_ burden and childlessness for having a university degree, we see an increased association of s_het_ burden with childlessness in females (OR=0.56 [95% CI 0.42-0.73], p=2.8×10^-5^; Extended Data Figure 8A; Supplementary Figure 17); but the association with male childlessness remains considerably stronger than in females.

## Discussion

In summary, we find that, in individuals of European ancestry within the UK Biobank, reduced reproductive success, especially in males, is associated with purifying selection acting on human genes, and that this may be mediated primarily by mate choice based on preferences relating to cognitive and behavioural traits. The mechanisms underlying this association have not been fully resolved here. Mate preferences are multi-dimensional and vary across cultures and time^24^, and studies of mate preferences are often biased to western contexts. It is likely that reduced reproductive success associated with increased s_het_ burden involves multiple cognitive and behavioural traits, not all of which will have been studied here. The negative association of s_het_ burden with measures of fluid intelligence, household income and educational attainment, together with the previously documented stronger female preference for reproductive partners with good financial prospects^22^ suggest that sex differences in mate preferences might contribute in part to the sex-difference in reproductive success with increased s_het_ burden. However, as we are not able to assess the association of s_het_ burden with all characteristics that are valued in a reproductive partner, especially those that are ranked most highly by both sexes (e.g. emotional stability and maturity)^22^, we cannot exclude the possibility that sex differences in the association of s_het_ burden with these traits also contribute to the sex difference in reproductive success. Nonetheless, this study is consistent with the relevance of Darwin’s theory of sexual selection^5^ to contemporary human populations. For further discussion of the limitations and implications of these findings, please see Supplementary Note 1 - Frequently Asked Questions.

We note that while this study demonstrates that rare heterozygous genetic variation has a much stronger association with reproductive success in males than females, Genome-wide association studies (predominantly in individuals with European ancestry) have suggested a greater contribution of common genetic variation to variance in reproductive success in females than males^40^, although the genetic correlation in total fertility between men and women is highly correlated^41^. As narrow-sense heritability of childlessness does not differ between sexes^42^, these two observations are potentially complementary: the larger contribution of heterozygous rare genetic variation to male reproductive success could be lowering the proportionate contribution of common genetic variation. Previous work demonstrated that the proportion of the genome that is homozygous is also associated with decreased reproductive success as measured through both increased childlessness and number of children ever born, although without an apparent sex-difference in the latter, and proposed that these associations are largely driven by rare homozygous variation, which we did not assess here^43^. Our study is focused on understanding why some genes might be under much stronger selective constraint than others, and not on a comprehensive analysis of the genetic basis of reproductive success.

These findings may help to explain why only a minority of highly constrained genes have been associated with genetic disorders that increase pre-reproductive mortality or cause infertility. While there are likely more genetic disorders to be discovered among these genes^44,45^, we anticipate that these highly constrained genes will not neatly divide into those that cause genetic disorders and those that reduce reproductive success. Rather, we predict that damaging variants in many of these genes will perturb neurodevelopment resulting in a spectrum of cognitive and behavioural outcomes, which may increase an individual’s risk of childlessness, but only result in a clinically-ascertainable condition in some individuals.

Sex-differential reproductive success potentially contributes to some other sex-differential patterns of genetic associations for cognitive and behavioural traits. For example, preferential transmission from mothers of alleles increasing risk of neurodevelopmental disorders has been related to the greater ‘resilience’ of females to such alleles^46^. However, our findings that the association of cognitive and behavioural traits with the damaging genetic variation studied here is similar between the sexes suggests that sex-differential reproductive success may be a more plausible explanation for some of these observations, as seen for the 22q11.2 deletion^47^; although, it is unlikely to explain sex-differential genetic associations in autism spectrum disorder^48^.

Childlessness can be voluntary or involuntary. A survey of childless individuals (at age 42) in the 1970 British Birth Cohort found that 28% of men and 31% of women reported not wanting to have children^49^. Involuntary childlessness can have serious consequences for mental health, and further studies of the factors influencing involuntary childlessness are warranted (see further discussion as part of Supplementary Note 1).

These analyses have several limitations. First, longitudinal relationship data for UK Biobank participants that might clarify the potential association of s_het_ burden with the ability to initiate and/or sustain a reproductive partnership during peak reproductive years is not available. Second, we have not investigated the association of s_het_ burden with the full range of cognitive and behavioural traits relating to mate preferences and reproductive success. We anticipate that teasing out the relative contributions of correlated cognitive and behavioural traits will be challenging. Third, UK Biobank participants are biased towards higher health, educational attainment and socioeconomic status^39^, and estimates of the negative association of s_het_ burden with reproductive fitness in UK Biobank possibly underestimate the true effects in the general population. Finally, we cannot account for as-yet-undiscovered male infertility genes in these analyses; nonetheless, our results based on clinically- and self-reported infertility (Extended Data Figures 4, 5) suggest a minor contribution of male infertility to the sex-differential relationship between s_het_ burden and childlessness.

Our study focused on individuals of European ancestry and analogous studies across different populations and cultures are needed. Males have considerably greater variance in reproductive success than females across cultures^50^, including higher levels of childlessness than females^27^, highlighting the potential for sexual selection acting on male reproductive success to act across populations. We also note that some trends relating to mate preferences and male childlessness have some evidence of being cross-cultural in nature^22,26,50^. We anticipate future studies that integrate genome-wide sequencing data on large population samples from a range of ancestries to more fully characterise the nature of selection pressures acting on damaging genetic variation in our species.

## Methods

### Sample Selection and Phenotype Collation

To collate phenotypes for all individuals in UK Biobank, we downloaded bulk phenotype files from the UK Biobank data showcase (https://www.ukbiobank.ac.uk/data-showcase/; data acquired 22 Jan 2020). Due to ascertainment biases with post-recruitment data (Supplementary Figure 15), we only retained data which were ascertained at time of recruitment as opposed to those ascertained via followup (i.e. instance 0 in the UK Biobank data showcase). Please see Supplementary Table 1 for detailed descriptions of all phenotypes assessed in this manuscript, which includes how each was processed, if applicable. Individuals with missing data for a relevant phenotype were excluded from analysis when testing that phenotype.

Following phenotype collation, we next selected for final analysis individuals of broadly European ancestry as determined by Bycroft et al.^51^, which left a total of 409,617 individuals. To identify and remove related individuals, we first downloaded the relatedness file from the UK Biobank data showcase using the ukbgene tool, which contains 107,124 relatedness pairs among UK Biobank participants^51^. Next, we sorted individuals by the total number of related pairs within this file, and removed the individual with the most related pairs and recalculated the total number of relationships for all other individuals. We repeated this process until no related pairs remained, leaving a total of 342,717 individuals for downstream analysis.

### Calling, Quality Control, and Annotation of Copy Number Variants from SNP Microarrays

To ascertain copy number variants from 488,377 UK Biobank participants with available genetic data^51^, we utilised the PennCNV CNV-ascertainment pipeline^52^. Raw CEL files were downloaded in 107 independent batches, of which 95 batches were genotyped with the standard UK Biobank array platform and 12 batches were genotyped with the UKBiLEVE array platform. Each batch was then processed independently through the following calling pipeline: first, raw CEL files were genotyped with Affymetrix power tools (http://media.affymetrix.com/support/developer/powertools/changelog/index.html) ‘genotype’ with default settings. Next, using the ‘generate_affy-geno_cluster.pl’ and ‘normalize_affy_geno_cluster.pl’ scripts provided as part of PennCNV, genotyped samples within each batch were clustered and normalised, respectively. Normalised clustering output was then split into one file per individual and provided as input to ‘detect_cnv.pl’ to generate an initial call set of CNVs. Finally, initial CNVs were then passed to the ‘clean_cnv.pl’ script with “-fraction” set to 0.25 in order to merge nearby CNV calls in each individual. Following CNV calling, we excluded all individuals with ≥ 20 CNVs and absolute waviness factor > 0.3, and all variants on either the X or Y chromosome, which left 485,593 individuals and 3,101,974 raw, redundant CNVs.

To perform quality control of ascertained CNVs, we developed a novel approach which uses individuals for which CNVs have been ascertained with both array and exome-based approaches. In short, we started with the basic logistic regression concept outlined in Mace et al.^53^ but instead used the intersect of array- and whole exome sequencing (WES)-ascertained CNVs as the dependent variable in a random forest model^54^, with various per-individual and per-CNV metrics as predictors. To train this model, we utilised an additional set of 46,856 individuals collected as part of the INTERVAL study^55^ genotyped on the same array as participants in UK Biobank, of which 4,465 also had matched WES data. For INTERVAL individuals, we performed array-based CNV calling identically to the method as described above and ascertained exome-based CNVs using three different algorithms with default settings: XHMM^56^, CANOES^57^, and CLAMMS^58^. For each INTERVAL participant for which we had both array and exome-based CNVs, we then determined a “WES overlap score” as a product of the overlap of each array-based CNV with the three WES-based callers, corrected for whether or not any overlap was possible due to probe/exon bias. Scoring results in a roughly continuous metric for each array-ascertained CNV of between zero and three, where zero represents a lack of overlap with any WES CNV call and three represents a perfect overlap with all three algorithms. For predictor covariates, we used several metrics already shown to be of high quality for CNV quality control^53,59^, per-CNV metrics based on these (e.g. mean log R ratio for each probe within a CNV rather than for all probes across an entire individual), and a novel metric which uses specific probes on the array known to be biased for CNV calls on bad arrays (Supplementary Table 3; see code availability). To determine estimated sensitivity/specificity of our model we performed 10-fold cross-validation, where all array CNVs which overlapped at least two exons were split into equal test and training sets and provided, separately for deletions and duplications, as input into the randomForest implementation in R as a linear predictor with nTrees set to 500. To generate a call set of final quality controlled CNVs for downstream analyses, we then trained a final random forest using all INTERVAL individuals with matched array and WES data and generated predicted WES overlap scores for all 3,101,974 raw UK Biobank CNVs identified with PennCNV as described above. CNVs were then filtered based on a predicted sensitivity of 95% based on cross-validation, leaving a remaining 1,612,831 CNVs (1,043,717 deletions, 569,114 duplications).

CNVs passing quality control were then provided as input to a custom java pipeline which merged all CNVs, regardless of whether they were deletions or duplications, based on 75% reciprocal overlap to generate a set of 173,871 nonredundant loci. Following filtering to 328,899 unrelated individuals of broadly European ancestry for which CNV data was available, each locus was quantified for allele frequency. Loci were then assessed for overlap with a set of known pathogenic CNVs identically to Crawford, et al.^59^ and annotated using Variant Effect Predictor (VEP) v97^60^. Only loci with an annotation of ‘transcript_ablation’ or ‘feature_trunctation’ and ‘coding_sequence_variant’ for deletions, and ‘transcript_amplification’ or ‘feature_elongation’ and ‘coding_sequence_variant’ for duplications were considered to be affecting a target gene. A total of 1,118,859 redundant CNVs remained for downstream analysis following all filtering and annotation (721,536 deletions, 397,323 duplications; Supplementary Figure 2).

### Processing SNV/InDel Data from WES

To collate protein truncating, missense, and synonymous variants for all 200,629 individuals whole exome sequenced by UK Biobank^11^, we downloaded the GRCh38-aligned population-level variant call format files from the UK Biobank (UK Biobank field 23156). All autosomal variants were then annotated with VEP v102^60^, CADDv1.6^61^, allele frequency from gnomAD^62^, and, where relevant, MPC^63^ and LOFTEE^62^. MPC scores were converted from build37 to build38 using the CrossMap tool^64^. Variants were assigned to a gene based on the primary ENSEMBL transcript with the most severe consequence. Variants were considered to be PTVs if they were annotated by VEP as having a splice acceptor/donor, stop gained, or frameshift consequence. We then retained only variants with a gnomAD-specific allele frequency < 1×10^-3^. Missense variants were only retained if they had MPC > 2 and CADD > 25. PTVs were only retained if they were annotated by LOFTEE as high confidence, had CADD > 25, and were not located in the last exon or intron of the canonical transcript as annotated by ENSEMBL^65^. We next applied genotype-level filtering where a given genotype was set to null (i.e../.) if the genotype had a depth < 7, genotype quality < 20, or a binomial test p.value for alternate versus reference reads for only heterozygous genotypes ≤ 0.001. If more than 50% of genotypes were missing for a given variant, that variant was filtered. These filtering approaches left a total of 11,455,900 redundant autosomal SNVs and InDels across all 139,480 unrelated individuals of broadly European ancestry included in this study (Supplementary Figure 3).

### Calculating s_het_ Burden for UK Biobank Participants

To calculate an individual’s s_het_ burden, assuming that fitness is multiplicative and that there is no epistasis between genes which are lost, we utilised the following formula:

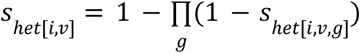

where *s*_*het*[*i,v*]_ indicates individual *i*’s s_het_ burden for variant class *v* and *s*_*het*[*i,v,g*]_ indicates the s_het_ score for gene *g* with a qualifying annotation for variant class *v*in individual *i*. As indicated by the formula above, s_het_ values were calculated independently for each variant type, with possible values for *v* of PTV, missense, synonymous, deletion, or duplication. Per-gene s_het_ values were obtained from Weghorn et al.^4^, under their demographic model which includes drift and scores for 16,189 protein coding genes which we were able to definitively map to an ENSEMBL gene ID. To ensure that our primary result of the association of s_het_ burden with childlessness is unaffected by the version of s_het_ we use to calculate our burden scores, we also utilised an earlier derivation of s_het_ from Cassa et al.^2^ which does not take into account a demographic model. Utilisation of s_het_ scores from Cassa et al.^2^ did not significantly change our primary result (Supplementary Figure 18).

To explore if genes known to be associated with male infertility were responsible for our observed association with male reproductive success, we also generated individual s_het_ scores for each variant class excluding a set of 150 autosomal genes known to be associated with male infertility (Supplementary Table 5 from Oud et al.^15^). Genes with an annotation of limited, moderate, strong, or definitive evidence were excluded from calculated s_het_ scores^16^. Similarly, and to test if a greater number of 742 genes associated with male infertility in mice were responsible for our observed association with male reproductive success, we queried all genes from Mouse Genome Informatics^16^ with a phenotype code of MP:0001925. Gene IDs were then translated to their human homologues and s_het_ burden scores excluding these genes were then generated and provided as input to logistic regression as described above.

To test if our observed relationship was robust when excluding genes with a known disease annotation, we also generated individual s_het_ scores where we removed 4,414 disease-associated genes. We considered a gene to be disease-associated based on being a confirmed or probable developmental disorder gene in the Developmental Disorders Genotype-Phenotype Database (DDG2P; https://decipher.sanger.ac.uk/info/ddg2p), in the Online Mendelian Inheritance in Man (OMIM; https://omim.org/) Morbid Map after excluding ‘non disease’ and ‘susceptibility to multifactorial disorder’ entries, or in ClinVar^66^ with a pathogenic/likely pathogenic variant linked to a phenotype.

### Logistic and Linear Modelling of Phenotypes

To test the association of each s_het_ burden (i.e. *s*_*het*[*i,v*]_) per variant class with a given phenotype (e.g. those in Supplementary Table 1), we used a general linear model via the ‘glm’ function in R of the form:

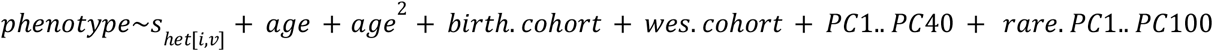

As age and year of birth are not perfectly correlated in the UK Biobank due to primary recruitment taking place over approximately 3 years, we also incorporated a categorical dummy term (*birth.cohort*) in our model for all possible 5 year birth cohort windows (e.g. born between 1940-1945, 1946-1950, etc.).*wes.cohort*represents UK Biobank sequencing batch, and is set to either 0, 1, or 2 for individuals without sequencing data (i.e. individuals with only CNV s_het_ scores), sequenced in the first batch of UK Biobank WES^67^, or as part of the newest batch of WES data^11^, respectively. Pre-computed ancestry principal components (*PC1*.. *PC*40) for each UK Biobank participant were generated by Bycroft et al.^51^ and acquired via the UK Biobank data showcase. To alleviate concerns about a potentially arbitrary selection of the number of ancestry principal components used in our models, we repeated our primary analysis of the association between having children and s_het_ burden in males with between 10 and 40 ancestry principal components and did not observe any change in our result (Supplementary Figure 19). To ensure that more recent ancestry was not biasing our results, we acquired a sparse identity-by-descent (IBD)-sharing matrix for 329,004 UK Biobank participants (thus excluding 13,713 participants) of unrelated European ancestry computed for the last 10 generations from Nait Saada et al.^68^ (data retrieved from UK Biobank return 3623). This matrix was provided as input to a custom Python script which computed the first 100 principal components for each participant using the ‘eigsh’ function from SciPy^69^. These principal components were then provided as a covariate in all models (*rare.PC*1.. *PC*100). As with PCs derived from common SNPs, we tested if the arbitrary inclusion of various IBD-derived PCs made a significant difference to our primary result and did not observe any significant deviation (Supplementary Figure 19).

All models were run separately for males and females, for all five possible variant classes, *v*, and at two allele frequency thresholds (Singletons and MAF < 1×10^-3^) for a total of 20 tests per phenotype; all tests were evaluated for significance based on a Bonferonni threshold of p < 2.5×10^-3^. For Supplementary Figure 6, we also tested two additional MAF cutoffs, < 1×10^-5^ and < 1×10^-4^. For binary phenotypes, ‘family’ was set to ‘binomial’ and for continuous phenotypes other than total number of children, ‘family’ was set to ‘gaussian’. When considering the overall number of children per participant (e.g. Figure 1A and Extended Data Figure 2) ‘family’ was set to ‘quasipoisson’ to account for overdispersion of the response variable. To combine the effect sizes or log odds ratios for CNVs and PTVs (e.g. for Figure 1A, B), we used the ‘metagen’ function from the ‘meta’ package^70^ in R to perform a fixed-effects meta analysis. For logistic regression, we set parameters ‘method.tau’ to ‘SJ’ and ‘sm’ to ‘OR’. For linear or quasi-Poisson regression, we set the parameter ‘sm’ to “SMD”. To avoid including an individual twice in our meta analysis, for samples with both CNV and PTV data available, we prioritised PTV-derived s_het_ scores.

When using raw variant counts as in Supplementary Figure 5, the s_het_ term in the above formula was changed to the total number of qualifying genes affected per individual, where qualifying genes were either those with pLI ≥ 0.9^62^ or those with s_het_ ≥ 0.15^4^. Individuals with > 3 genes lost for deletions (pLI ≥ 0.9 n = 28; s_het_ ≥ 0.15 n = 12) and PTVs (pLI ≥ 0.9 n = 1; s_het_ ≥ 0.15 n = 0) were removed prior to regression analyses. To provide a negative control for our association tests, we also performed associations for several neutral phenotypes we hypothesised to not be under negative selection: fresh fruit intake, handedness, and blonde hair colour (Supplementary Table 1). None of these associations were significant after correcting for multiple testing (Supplementary Figure 20).

To test the association of individual phenotypes with the likelihood of having children (Extended Data Figure 6), we used a logistic model with the ‘family’ parameter of the ‘glm’ function set to ‘binomial’ in R of the form:

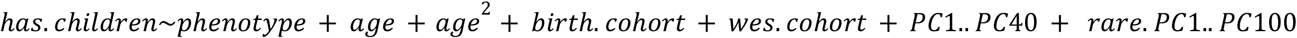

As with estimating the contribution of s_het_ burden to phenotypes, all analyses were run separately for both males and females and with a WES batch covariate as necessary. For all models involving household income, we additionally included partner at home status as a covariate, as household income was recorded per household, not per recruited individual.

All odds ratios, effect sizes, standard errors, p values, and total individuals per association test reported in this manuscript can be found in Supplementary Table 4.

### Collation and Testing of Participant Medical Data

To assess if a broad range of medical conditions play a role in mediating the association of high s_het_ burden with childlessness, we queried two relevant datasets provided by the UK Biobank^10^: hospital episode statistics (HES) and combined health outcomes data (CHOD; Supplementary Table 1). Briefly, for each UK Biobank participant, HES data incorporates electronic inpatient data provided directly from NHS hospitals and CHOD aggregates HES, general practitioner records, self-reported conditions, and death records. All data sources are coded according to the International Classification of Disorders v10 (ICD-10). For the purposes of this work, we ignored all cancer codings from HES and CHOD data (ICD chapter II and codes O00-O08 of chapter XV), and instead used independent cancer registry data; the UK Biobank acquires information on cancer diagnoses from the UK cancer registry which aggregates a wide range of data sources including general practitioners, nursing homes, and hospitals and is considered the more accurate data source for UK Biobank participant cancer diagnoses. For ease of display, tests involving cancer codes are shown with HES data (Extended Data Figure 4, 5, Supplementary Figures 8, 9).

We utilised both HES and CHOD sources to examine a broad set of medical conditions in UK Biobank participants. Complete HES are available for all UK Biobank participants but are probably depleted of conditions that are unlikely to be seen in a hospital setting (e.g. male infertility). CHOD are also likely to be more sensitive to a wide variety of conditions as they incorporate aforementioned HES data with both general practitioner records and self-reported outcomes. Importantly, while general practitioner records are only available for 46% (n = 230,090) of UK Biobank participants, when we tested for an association between s_het_ burden and whether a participant had general practitioner records or not, we did not observe a significant association for either males (OR=1.03 [95% CI 0.80-1.31], p=0.84) or females (OR=1.14 [95% CI 0.91-1.43], p=0.25; Supplementary Figure 15)^37^. This indicated that we were unlikely to see biases due to including CHOD from individuals who were missing general practitioner records. As such, we prioritised the use of CHOD in most analyses presented in the text, figures, and supplementary information of this manuscript – complete results for all codes in both HES and CHOD data are available as Supplementary Table 2. Exceptions include when testing codings from chapters XVII to XXII which are beyond the diagnostic scope of CHOD, and cancer codes better ascertained from the UK Biobank cancer registry as noted above (Extended Data Figure 4, 5; Supplementary Figures 8, 9).

To determine the role of 19,154 and 2,361 diseases, disorders, and special codes collated from HES and CHOD, respectively, in the relationship between s_het_ burden and childlessness (Extended Data Figure 4, 5), we used a modified version of our primary logistic model of the form:

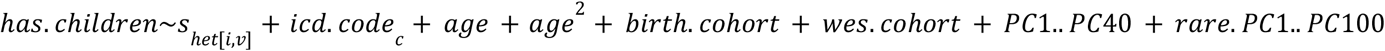

Where *icd.code*represents a binarised presence/absence of one of 19,154 different ICD-10 diseases, disease groups, and chapters, *c*. Tests were only performed when a given code was represented by at least 2 individuals in both genetic (i.e. CNVs) and WES (i.e. PTVs) data. When considering individuals who have a particular code, *c*, we utilised the hierarchical information present within the ICD-10 coding system. For example, when we tested if inclusion of a term for individuals with non-insulin-dependent diabetes mellitus (ICD-10 code E11) has an association with childlessness, we also considered individuals with any sub-code (i.e. E11.0-E11.9). This same principle was also used for disease groupings – when testing the more general diabetes mellitus group (ICD-10 block E10-E14) we included all individuals with any code between E10 and E14, including disease subtypes (e.g. E11.0). For each model, we retained both an odds ratio for the association of individual s_het_ burden and presence of a given code with childlessness (Supplementary Table 2).

### Evaluation of Gene Expression

To determine the expression in various tissues of all genes assessed in this study, we downloaded processed median transcripts per million (TPM) values for all genes provided by v7 of the GTEx study^30^ (https://storage.googleapis.com/gtex_analysis_v8/rna_seq_data/GTEx_Analysis_2017-06-05_v8_RNASeQCv1.1.9_gene_median_tpm.gct.gz). Only genes for which an s_het_ score was available were retained from this file.

To evaluate genes known to be involved in male infertility, we assessed if each gene was affected by either a private deletion or PTV in a UK Biobank individual. We then plotted *ln* (*testis expression*)in two ways: (i) as a factor of being a male infertility gene or not or (ii) having or not having a qualifying variant (Supplementary Figure 7).

To assess if genes expressed in a particular tissue modulated the relationship between s_het_ burden and childlessness, we also generated tissue-specific s_het_ scores for all individuals where we excluded all genes with a tissue-specific TPM value greater than 0.5. s_het_ scores for all 53 tissues were calculated as described above and used to test the association between s_het_ burden and childlessness (Supplementary Figure 10).

To examine the correlation between expression and s_het_, we performed two linear regressions in all 53 tissues queried by the GTEx study: 1) the association between s_het_ and *ln* (*expression_tissue_*,)corrected for gene coding sequence length and 2) the association between s_het_ times the number of singleton LoF variants and *ln* (*expression_tissue_*)corrected for gene coding sequence length. For each tissue and model, we extracted the overall variance explained for each model (r^2^) and plotted it as part of Supplementary Figure 11.

### Modelling the Contribution of Phenotypes to Observed Reduction in Fitness

#### Variant s_het_ Burden

To estimate the contribution of s_het_ to overall fitness (Extended Data Figure 8B), we extracted log odds ratio estimates for the association of s_het_ with having children from our logistic model and estimated the proportion of childless individuals at various s_het_ scores (0 to 1 at 0.1 intervals; Extended Data Figure 9). To calculate the error in our estimates (i.e. the shaded areas in Extended Data Figure 8B), we used the 95% confidence intervals for the s_het_ burden log odds ratio from our original logistic regression. We note that we assume that input parameters calculated with UK Biobank data are generalisable to the UK population as a whole, but may slightly underestimate the error in our final result. Please see Supplementary Note 2 for a more detailed description of how the contribution of s_het_ to overall fitness was calculated.

#### General Cognition

When possible, we used independent estimates from population level or external data to alleviate biases in UK Biobank phenotype ascertainment (Supplementary Figure 15). As such, data on cognitive ability and fertility are collected from Swedish population-level government administrative registers that have been linked to Swedish conscription registers^71^. To assess assignment into different branches of a universal conscription for Swedish men, the Swedish government included an extensive cognitive ability test which all men in Sweden had to take part in. Information on childbearing is based on birth records, and linkage to both men and women is nearly universal, partly due to universal government identity numbers, combined with serious paternity investigations in case of missing information of the biological father. This information was used to calculate reproductive fertility histories in 2012 for all men included in this study. We include data on all Swedish born men who participated in the military conscription test at age 18-20 who were born 1965-1967. The conscription registers are described in more detail elsewhere^34,72^.

For the current study, we did not rely on the official cognitive ability scores assigned for each man following their cognitive ability test as in Kolk and Barclay^34^, but instead made manual calculations to create a more finely grained measure from raw test scores based on a battery of cognitive ability tests that are available for 3 years in our conscription registers. The Swedish military created an official IQ-measure based on a 9-score stanine scale that has been used in a large number of scientific studies^34,73^. In the current study we developed a more detailed score using information on the actual test scores of men participating in the test. The conscription test consisted of four large subtests measuring different dimensions of IQ with logical, spatial, verbal, and technical subtest^72,74,75^. To get a more finely tuned IQ measure than the official stanine measure we used the raw test scores of each of these four tests and summed the total number of correct questions for these four sub-tests. Within each stanine IQ score, we then examined the distribution of test scores and after standardising the test scores using only variation within each stanine score, calculated a new detailed IQ score. This procedure is done to anchor our new IQ measure in the official stanine IQ score. As our test scores have some missing values for men with very high and very low stanine scores, this procedure results in a slightly underdispersed distribution and our new calibrated IQ score has μ = 100 & σ = 12, as compared to the official stanine measure with μ = 100 & σ = 15.

This score allows us to calculate cognitive ability by single digit IQ scores (Supplementary Table 5); however, as we had to rely on only observations with complete test scores for all test batteries, our data has a higher share of excluded men than the official cognitive ability scores (used by Kolk and Barclay^34^ and others). In addition to the ~5% of men that did not take the test (e.g. they were ineligible for military service due to handicap such as visual impairments, that they were abroad, or were conscripted at an atypical age), we additionally excluded a number of men for which scores of all test batteries were not available. Our manually computed fine-grained measure was later standardised against the official cognitive ability test score to maintain comparability and to assure our slightly smaller population is still representative of the complete cohort. Compared to most other measures of cognitive ability in the scientific literature, we argue that our population is unusually representative as little (indirect) pre-selection due to cognitive ability took place.

We first estimated the association of overall s_het_ burden with fluid intelligence (Figure 2F) and, because fluid intelligence is normalised and IQ is normally distributed, converted this effect size to a predicted change of IQ. To then estimate childlessness and fertility for low IQ values not actually observed in the general population, we fit actual observations to a sigmoidal model using the function ‘nls’ in R (Supplementary Figure 14; Supplementary Table 5). As our empirical distribution did not conform to a standard test distribution, we then simulated 100,000 individuals, with IQ values for each individual randomly selected from our original Swedish IQ distribution with the mean shifted by the expected reduction in IQ as explained by our s_het_ model. We then assigned each simulated individual an expected number of children and predicted probability of childlessness based on their simulated IQ value as given in Supplementary Table 5. Number of children across all 100,000 individuals was then averaged to generate an expected mean fertility for a given s_het_ score (Supplementary Figure 13). This value was then compared to the mean number of children for the unburdened population via the following formula:

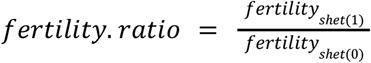

To then calculate the proportion of reduced reproductive success explained by IQ (and by extrapolation, other traits) we then divide this fertility ratio by the overall reduction in fitness given by s_het_ as described above. Please see Supplementary Note 3 for a more detailed example of how we performed this calculation.

#### Mental Health Disorders

As with general cognition, we used estimates from external studies to alleviate biases in UK Biobank phenotype ascertainment (Supplementary Figure 15)^37^. In this case, as we were unable to accurately estimate the increased risk of developing individual mental health disorders as a factor of individual s_het_ burden, we instead utilised odds ratios from Ganna et al.^9^. Only odds ratios for schizophrenia, autism spectrum disorder, and bipolar disorder were retained. As Ganna et al.^9^ estimated the risk based on total count of high pLI (≥0.9)^62^ genes with PTVs per individual instead of with s_het_, we assumed that an individual carrying one such variant had an s_het_ burden of 0.162, or the mean s_het_ value of all high pLI (≥0.9) genes. We then converted this into a proportion of individuals with a given mental health disorder, *t*, at s_het_ burden, *x*, by scaling the odds ratio with the following formula:

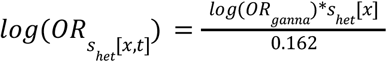

To establish a baseline expectation for the prevalence of each mental health disorder at s_het_ 0 we utilised population-level data from Power et al.^20^ and extrapolated for each trait at increasing s_het_ values (Supplementary Figure 21). To generate an expected mean number of children for simulated individuals with mental health disorders, we used fertility statistics generated by Power et al.^20^. As Power et al.^20^ did not provide childlessness data, we were unable to generate expected childlessness as we did for other traits. Overall predicted reduced fitness attributable to mental health and all values used for performing the above analyses are provided in Supplementary Table 6.

#### Having a Partner at Home, Having had Sex, Educational Attainment, and Infertility

To determine the contribution of having a partner at home, ever having had sex, educational attainment, and a medical diagnosis of infertility to the relationship between s_het_ burden and childlessness, we utilised a multiple regression model incorporating various combinations of these traits (Extended Data Figure 8A; Supplementary Figure 17). First, we calculated the variance in childlessness explained by a null model consisting of age, age^2^, birth cohort, WES batch, the first 40 ancestry PCs, and the first 100 IBD-derived ancestry PCs using Nagelkerke’s pseudo-r^2^as calculated using the “nagelkerke” function from the R package “rcompanion” (https://rcompanion.org/handbook/). Next, to determine the proportion of variance explained in childlessness by s_het_ alone, we calculated incremental pseudo-r^2^ between this null model and a model additionally incorporating a term for PTV s_het_ burden. We then repeated this analysis, except now including an additional covariate (e.g. having a partner at home) to determine the reduction in variance explained by PTV s_het_ when correcting for the additional covariate. This reduction in variance was then converted to a percent change via the following formula:

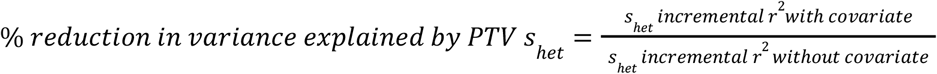

This basic analysis was then repeated for all possible combinations of having a partner at home, ever having had sex, having a university degree, and having a medical diagnosis of infertility (Extended Data Figure 8A; Supplementary Figure 17). Percent reduction in variance explained values plotted in Extended Data Figure 8A and Supplementary Figure 17 are displayed for s_het_ calculated using PTVs only.

Relatively complete mental health disorder data are available for all individuals via the complete health outcomes data; therefore, we also included a covariate for having a mental health disorder as a covariate in our multiple regression model (Extended Data Figure 8A). As income is provided by UK Biobank on a per-household basis (Figure 2E) and the number of individuals with fluid intelligence data recorded at recruitment is significantly smaller than for other covariates (Figure 2F), we did not include these as part of our multiple regression model.

## Supporting information

SuppTable1

SuppTable2

SuppTable3

SuppTable4

SuppTable5

SuppTable6

## Acknowledgements

We thank the reviewers for constructive comments and criticism. We thank Leo Parts, Joanna Kaplanis, Molly Przeworski and George Davey-Smith for useful discussions and advice on data analysis. We thank Manon Oud and Joris Veltman for helpful discussions regarding infertility. We thank Georgios Kalantzis and Pier Francesco Palamara for assistance with correcting for recent ancestry. We thank the INTERVAL study for sharing genotyping and exome data that allowed us to refine our CNV filtering methodology. This work has been funded by core Wellcome funding to the Wellcome Sanger Institute (grant WT098051) and as part of a Medical Research Council (MRC) Centre Grant to the MRC Centre for Neuropsychiatric Genetics and Genomics (grant MR/L010305/1). K.B. is supported by a fellowship from the Bank of Sweden Tercentenary Foundation (Riksbankens Jubileumsfond). This work has been conducted using the UK Biobank Resource under application numbers 14421 (to G.K.) and 44165 (to H.C.M.).

## Author Contributions

E.J.G, M.D.C.N, and K.E.S. assessed the contribution of rare genetic variation to the phenotypes and vital statistics presented in this manuscript. E.J.G and G.K. performed CNV calling. E.J.G. and M.E.K.N. annotated and assessed SNV and InDel variants from provided WES data. K.B., M.K., E.J.G, and M.E.H. curated and analysed Swedish IQ data. E.J.G., K.E.S., H.C.M., and M.E.H. designed experiments, oversaw the study and wrote the manuscript.

## Data Availability Statement

Raw data produced by this study are available as part of the UK Biobank data returns catalogue with application ID 44165: https://biobank.ndph.ox.ac.uk/ukb/docs.cgi?id=1.

## Code Availability Statement

Code used as part of this project to perform phenotype testing, CNV calling, variant quality control, and generate all main text figures, supplementary figures and supplementary tables is available on github: https://github.com/HurlesGroupSanger/UKBBFertility. All statistical analysis in this manuscript was performed using R v3.6.0.

## Extended Data Figures

**Extended Data Figure 1.**
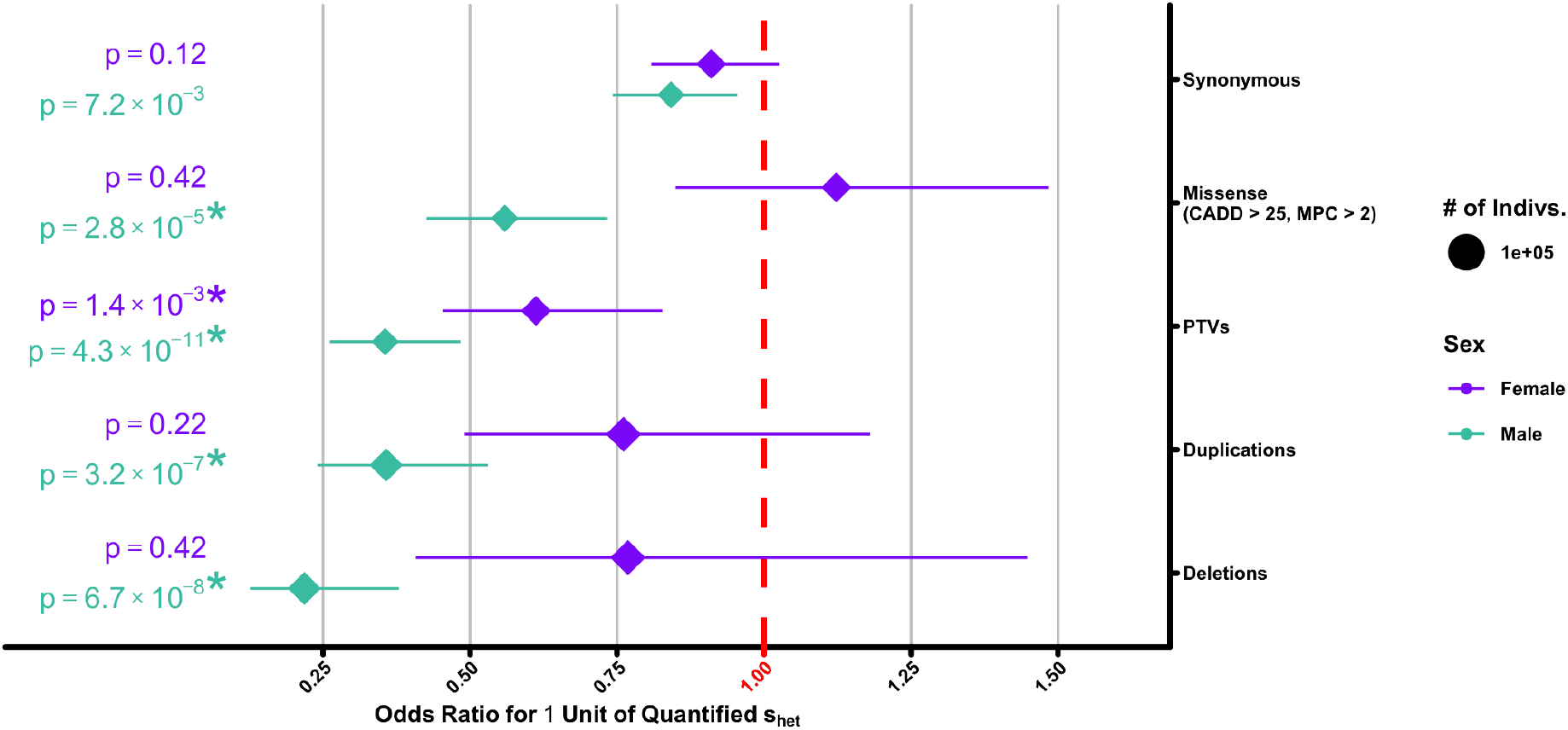
Odds ratio estimates for the association of s_het_ burden with having children for all variant classes. Identical plot to main text Figure 1A, but with additional data for synonymous, missense, and duplication s_het_ scores, separated into females (violet) and males (jade). Asterisks indicate significance after Bonferroni correction for 20 tests (p < 2.5×10^-3^; Methods).

**Extended Data Figure 2.**
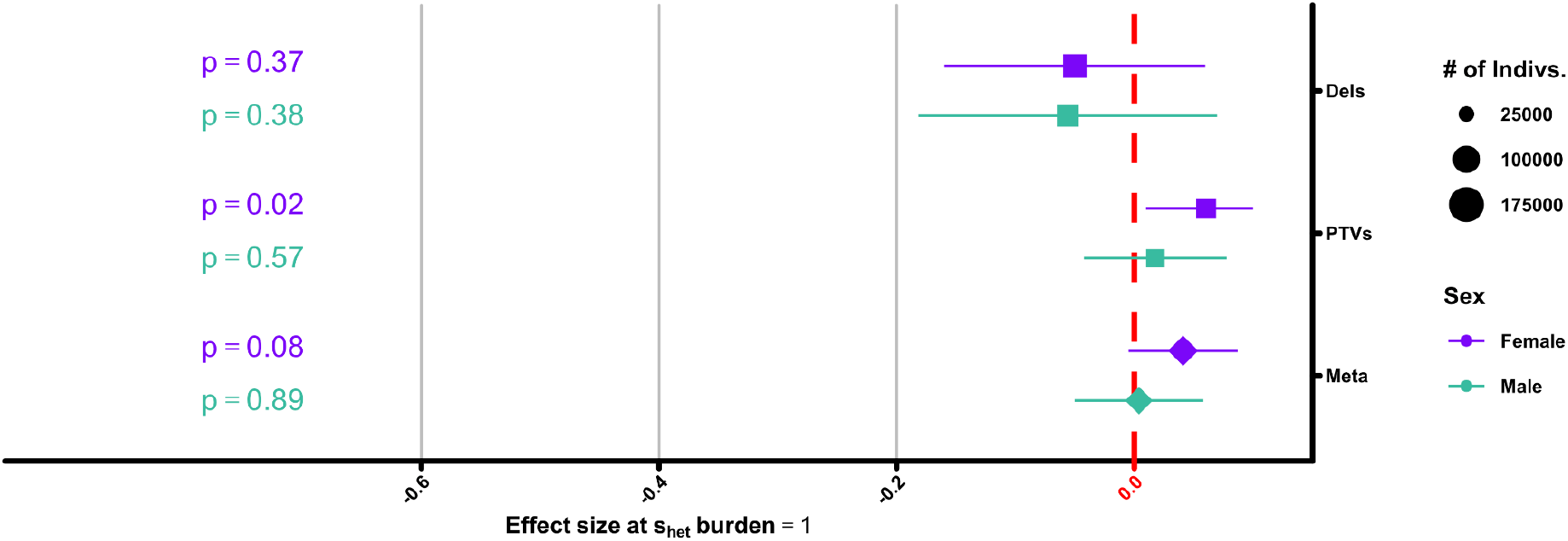
Effect size estimate for the association of s_het_ burden with number of children for individuals with children. Shown are the effect size estimates for the association of s_het_ burden with number of children, separated into females (purple) and males (jade), but with all childless individuals in the UK Biobank removed. Like Main Text Figure 1A, the regression used to generate the displayed result used the raw number of children, live births for females and children fathered for males, rather than a binary value for having children. Asterisks indicate significance after Bonferroni correction for 20 tests (p < 2.5×10^-3^; Methods).

**Extended Data Figure 3.**
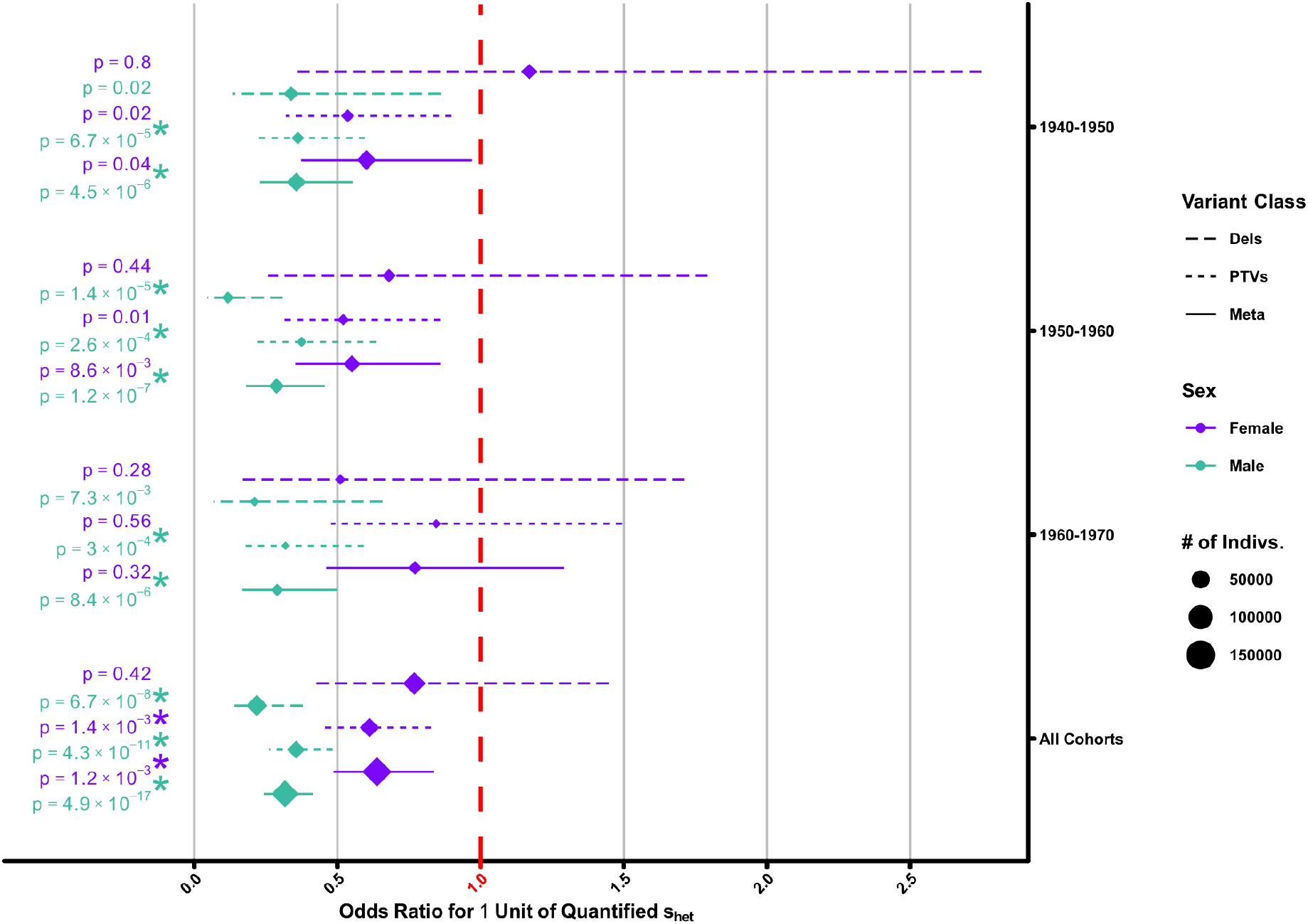
Odds ratio estimate for s_het_ burden stratified by age group. Shown are odds ratio estimates for the association of s_het_ burden with having children, stratified by participant age (y-axis) and separated into females (violet) and males (jade). Age range intervals are left-open. Dash of the line indicates whether the estimate comes from s_het_ burden calculated from deletions (long dash), PTVs (short dash), or from a fixed effects meta-analysis (no dash). Also shown for reference are the results for all individuals regardless of age (All Ages), which is identical to the result shown in main text Figure 1B. Asterisks indicate significance after Bonferroni correction for 20 tests (p < 2.5×10^-3^; Methods).

**Extended Data Figure 4.**
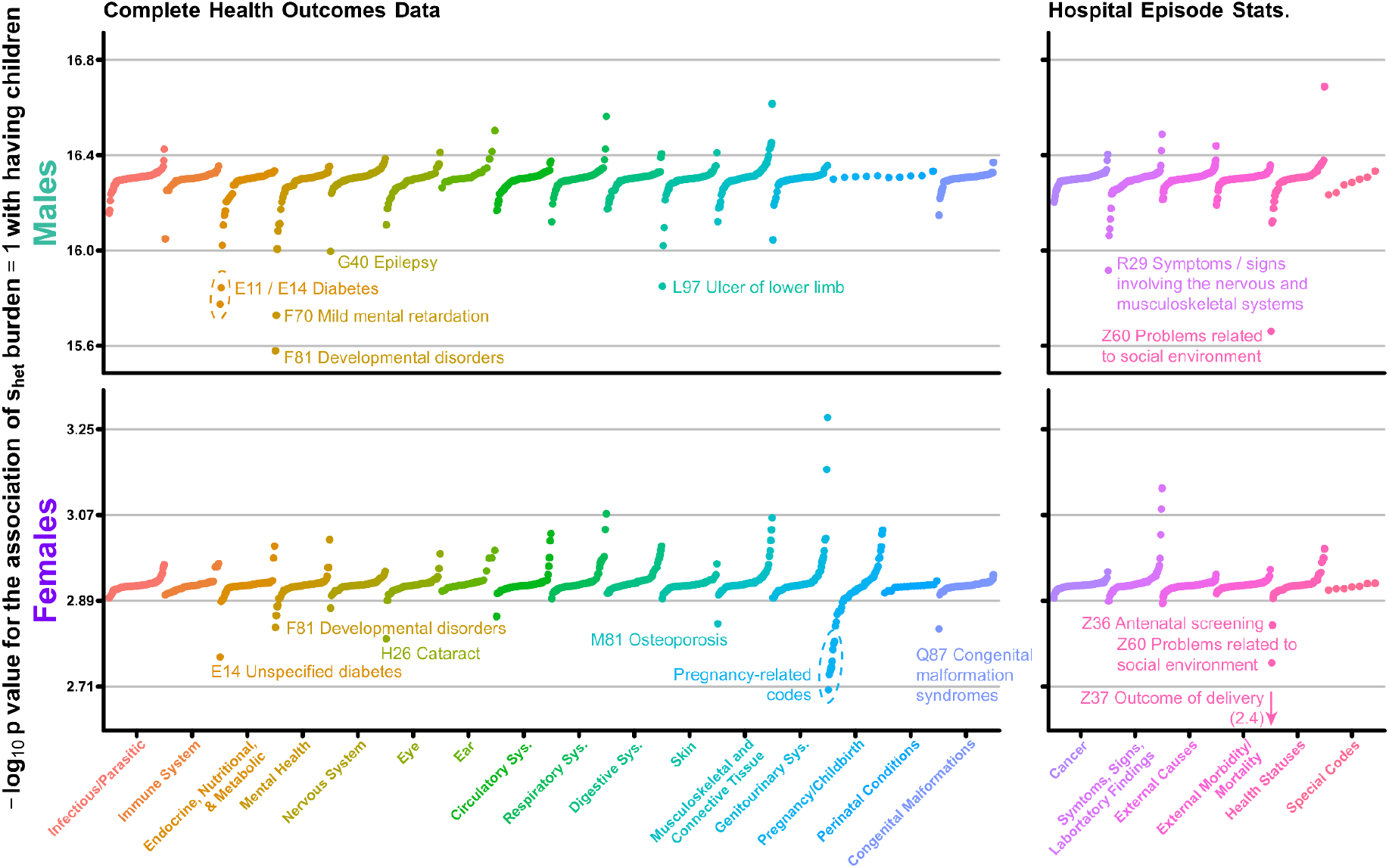
Mediation of the relationship between s_het_ burden and childlessness by various disorders. Plotted is the deletion and PTV meta-analysis -log_10_ p value for the association between s_het_ burden and having children, corrected by one of 1,294 ICD-10 codes from a combination of general practitioner, hospital episode records, and self-reported conditions (left) or hospital episode records alone (right) separately for males (top) and females (bottom). Remaining ICD-10 codes at different levels on the ICD-10 hierarchy not displayed here are plotted in Supplementary Figures 8 and 9. Results are ordered first by ICD-10 chapter (x-axis) and then by increasing -log_10_ p value (y-axis). The arrow for code Z37 indicates the point is below the scale of the y-axis with -log_10_ p value indicated in parentheses. Visual outliers are labelled and do not imply a significant change in the effect size of s_het_ on childlessness.

**Extended Data Figure 5.**
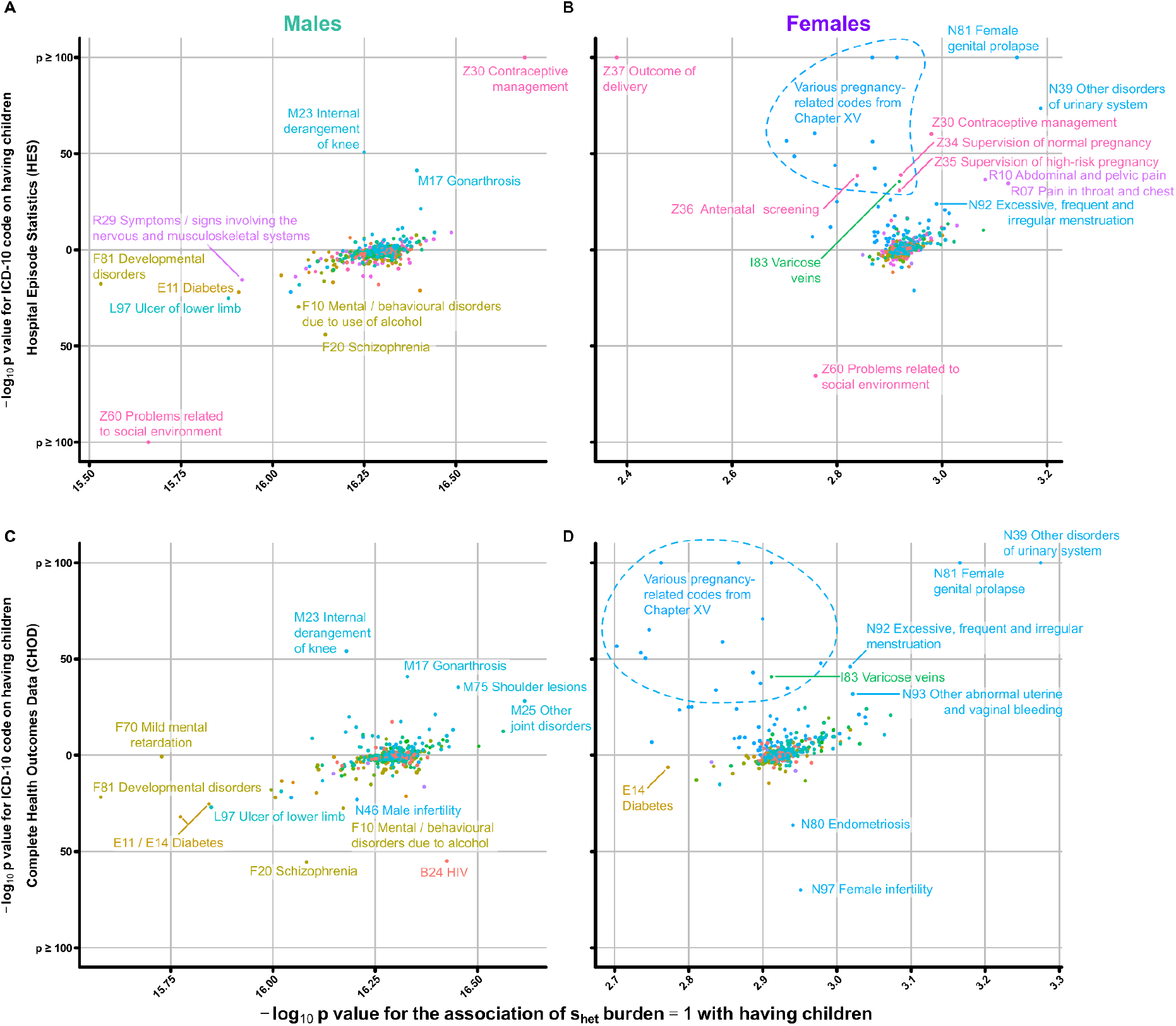
Mediation of the association between s_het_ burden and childlessness by various disorders. Depicted are the results of our primary association between childlessness and individual s_het_ burden corrected for presence/absence of approximately 2,000 different disorders, diseases, and health factors queried from (**A,B**) hospital episode statistics and (**C,D**) complete health outcomes data as represented by the ICD-10 medical coding system separately for (**A,C**) males and (**B,D**) females (see main text methods). Shown on the x-axis is the -log_10_ p value for the association of s_het_ with having children, corrected for a given diagnostic code. On the y-axis is the -log_10_ p value for having a given medical code on likelihood of having children; p values are placed above or below y = 0 based on the direction of effect, with disorders which are associated with having children above and those associated with not having children below. Codes were chosen for labeling to highlight outliers and not based on any statistical criteria. Codes with points at the top or bottom of plots have -log_10_ p values ≥ 100. Color of points and text is based on the ICD-10 chapter. Please note that text labels do not necessarily represent the full official name of a given ICD-10 code.

**Extended Data Figure 6.**
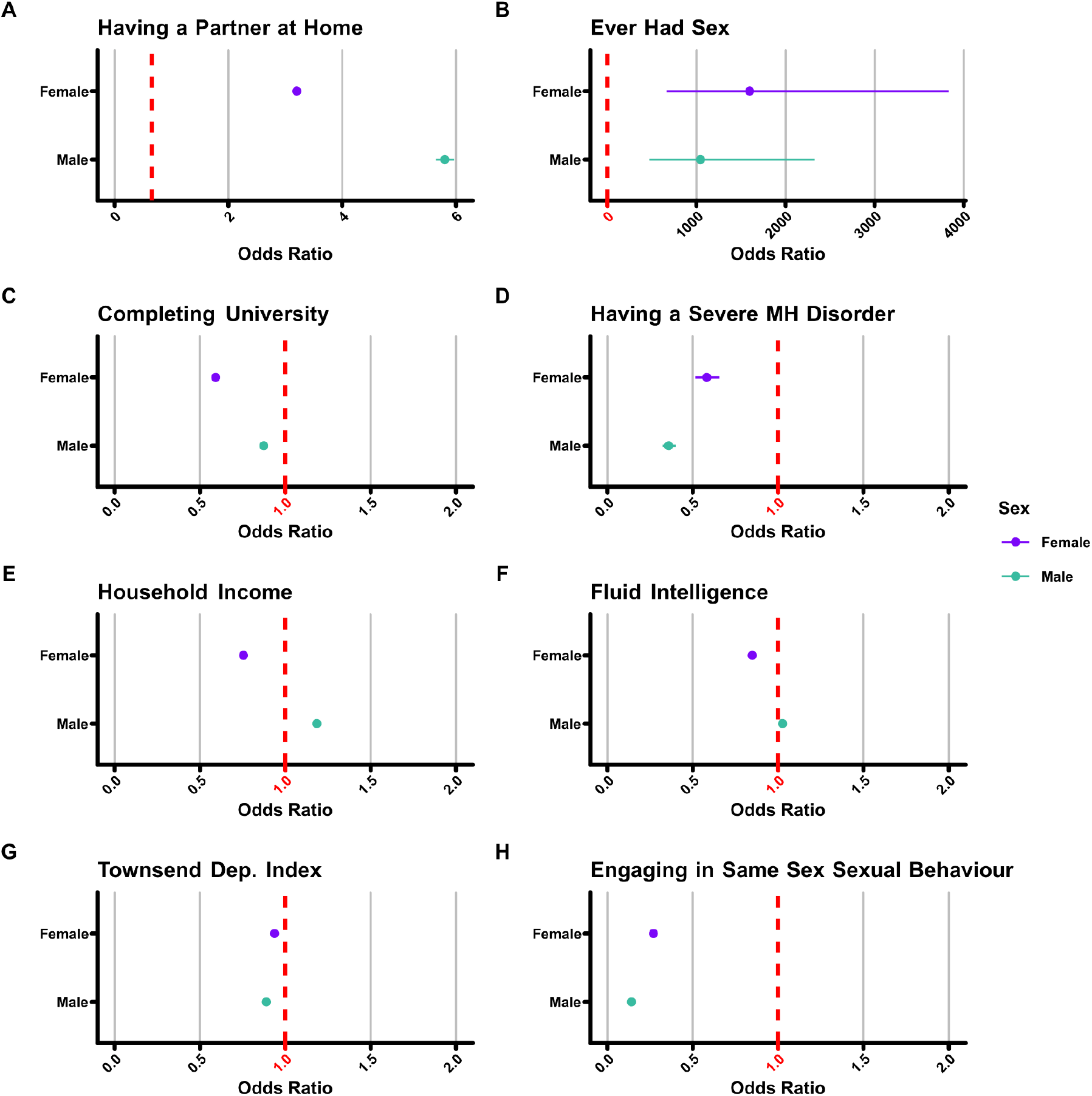
Association of eight relevant phenotypes/demographic measures with the likelihood of having children among UK Biobank participants. Shown are the results of a logistic regression estimating the odds ratio for the relationship of (**A**) having a partner at home, (**B**) ever having had sex (**C**) completing university, (**D**) having a severe mental health disorder, (**E**) household income, (**F**) fluid intelligence, (**G**) Townsend deprivation index, and (**H**) engaging in same sex sexual behaviour with likelihood of having children, separated into females (violet) and males (jade). 95% confidence intervals for all plots are included, but may be invisible at the resolution of the figure. Please note that the scales of the x-axis for plots (**A**) and (**B**) are different from plots (**C-H**) due to the relatively stronger association of these traits with having children.

**Extended Data Figure 7.**
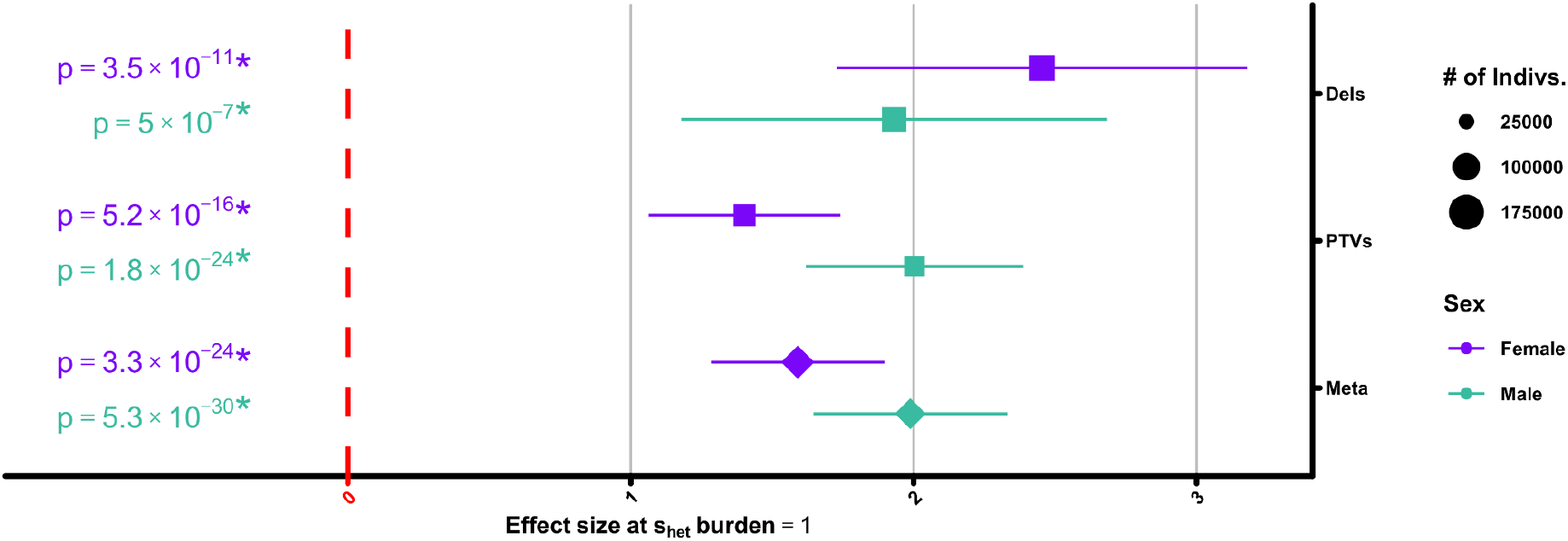
Effect size estimates for the association between s_het_ burden and Townsend Deprivation Index. Shown are the effect size estimates for the association between s_het_ burden and Townsend Deprivation Index, separated into females (purple) and males (jade). Units are unnormalized Townsend Deprivation Indices for each individual in the UK Biobank. Asterisks indicate significance after Bonferroni correction for 20 tests (p < 2.5×10^-3^; Methods).

**Extended Data Figure 8.**
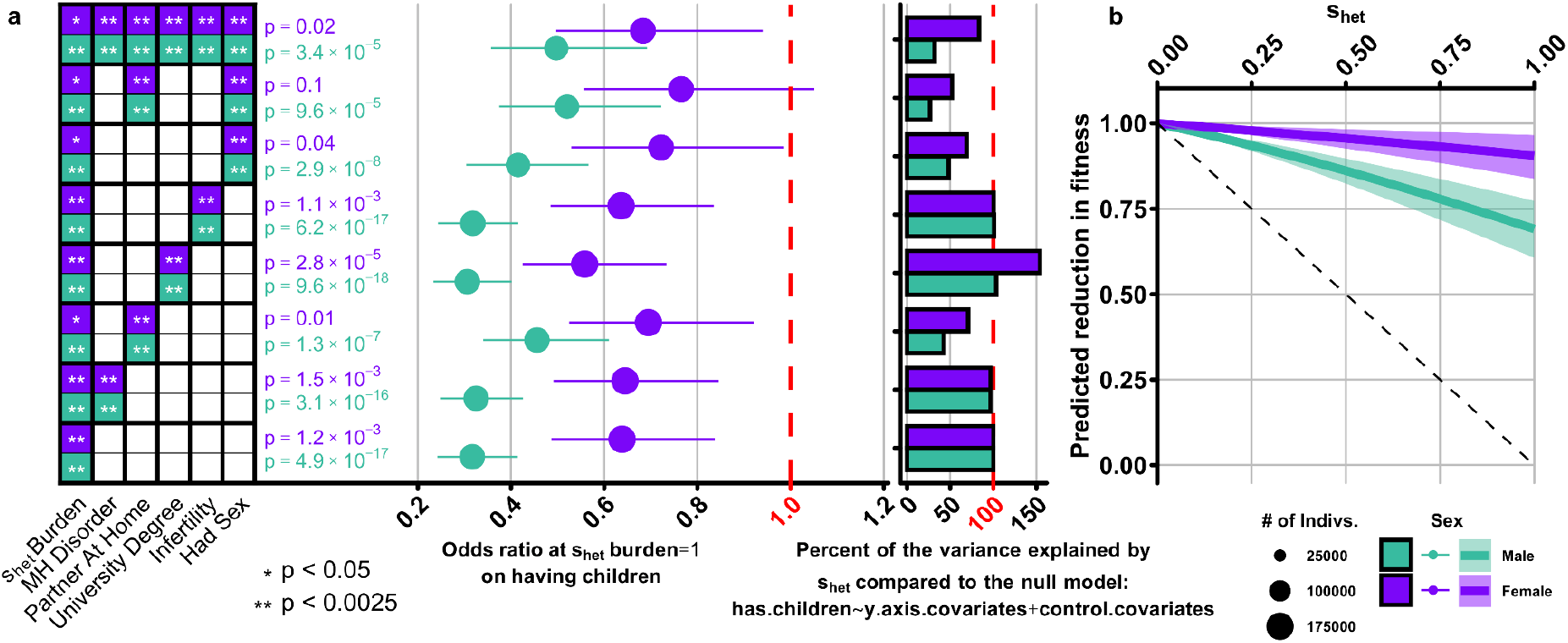
The role of individual phenotypes in the relationship between s_het_ burden, childlessness, and fitness. (A) Odds ratio estimates for the association of cumulative deleterious variation for a combined meta-analysis (deletions + PTVs) with childlessness (middle), corrected for a combination of whether or not a study participant has a mental health (MH) disorder, a partner at home, a university degree, infertility (as ascertained from Complete Health Outcomes Data; Methods), or ever had sex; traits included in each model are indicated as coloured boxes (males – jade, females – violet) on the y-axis. Stars within boxes indicate either nominal (*) or Bonferroni-corrected (**) significance level with childlessness for each covariate independently when correcting for PTV s_het_ burden. For all possible combinations of these traits, see Supplementary Figure 17. As indicated by coloured boxes, all models include s_het_ burden and were run separately for males and females. The marginal bar plot to the right gives the proportion of the variance in childlessness explained by s_het_ burden as calculated for PTVs only, scaled to the model which only includes s_het_ burden (i.e. the model on the bottom of the plot). (B) Predicted reduction in overall fitness as a factor of individual s_het_ burden. Displayed is the expected reduction in fitness as a factor of increasing s_het_ burden, independently for each sex. Error is shown as the lighter shaded area surrounding the trend line, and is based on the confidence intervals on the odds ratio as determined by our logistic regression model (Figure 1B; Methods). The dashed line represents the theoretical reduction in fitness as predicted by s_het_^4^.

**Extended Data Figure 9.**
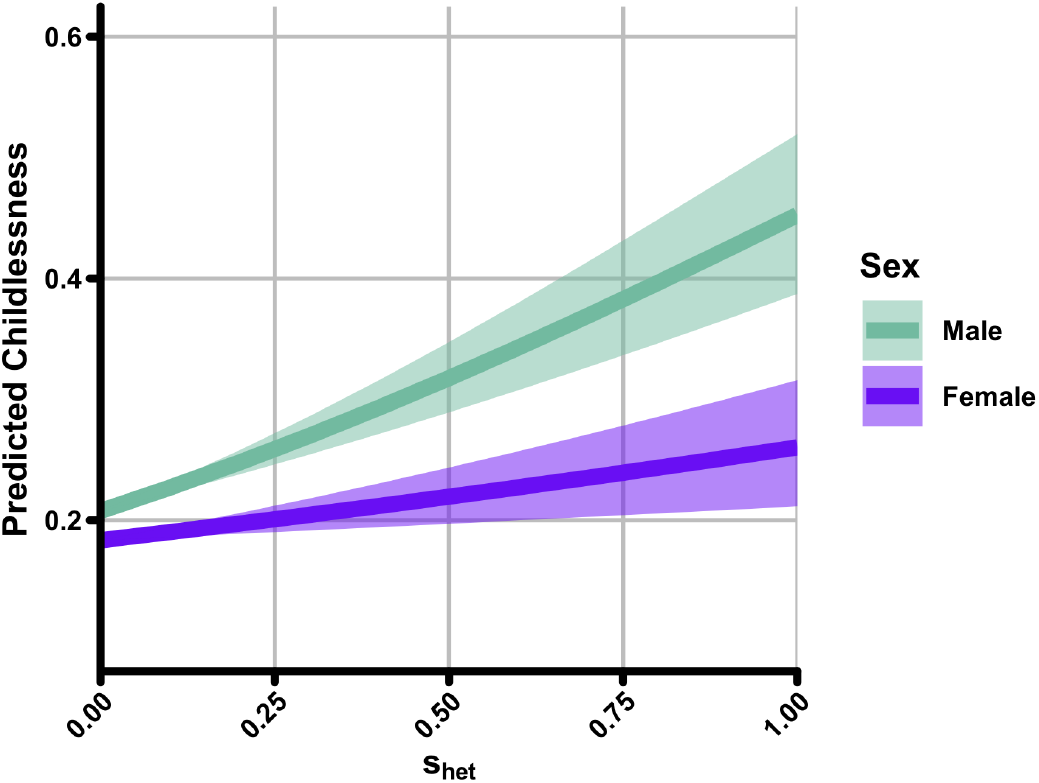
The association of s_het_ burden with childlessness. Identical to Extended Data Figure 8B, except in this instance, the y-axis represents predicted childlessness as a factor of individual s_het_ burden, rather than predicted reduction in fitness. Values at x = 0 represent actual mean childlessness among UK Biobank males (jade) and females (violet) with an s_het_ burden of 0.

