## Supplementary Materials for "Reduced reproductive success is associated with selective constraint on human genes"

### Table of Contents

**Supplementary Figures** **3**

1. Vital statistics for UK Biobank participants 3
2. Characteristics of CNVs in the UK Biobank 4
3. Characteristics of whole exome sequencing-ascertained variants in UK Biobank 5
4. $s_{het}$ distribution stratified by sex and number of children 6
5. Estimates for the impact of raw deleterious variant count on having children 7
6. Odds ratio estimate for the association of $s_{het}$ burden on having children when using various rare variant allele frequency cutoffs 8
7. Expression of genes in testis 9
8. Change in the p. value of the association between $s_{het}$ burden and male reproductive success after inclusion of ICD-10 codes 10
9. Change in the p. value of the association between $s_{het}$ burden and female reproductive success after inclusion of ICD-10 codes 11
10. The association of $s_{het}$ burden with childlessness when excluding genes based on expression status in GTEx 12
11. Correlation of per-gene tissue-specific expression with $s_{het}$ 13
12. Odds ratio estimates for the association of $s_{het}$ burden on likelihood of engaging in same sex sexual behaviour 14
13. Association of fluid intelligence with fitness 15
14. Population level IQ data from Swedish military records 16
15. Estimates for the association of $s_{het}$ burden with various recruitment biases in the UK Biobank 17
16. The association of $s_{het}$ burden with individual mental health disorders 18
17. Multiple regression models 19
18. Comparison of $s_{het}$ burden calculated with and without a demographic model 20
19. Investigation of the role of ancestry principal components in childlessness 21
20. Phenotypes not expected to have any relationship with $s_{het}$ burden 22
21. Association of mental health disorders with fitness 23

**Supplementary Tables** **24**

**Supplementary Notes** **25**

1. Frequently Asked Questions (FAQs) 25
2. Calculation of the contribution of $s_{het}$ to overall fitness 40
3. Calculation of the contribution of fluid intelligence to overall fitness 42

**Supplementary References** **44**

### Supplementary Figures

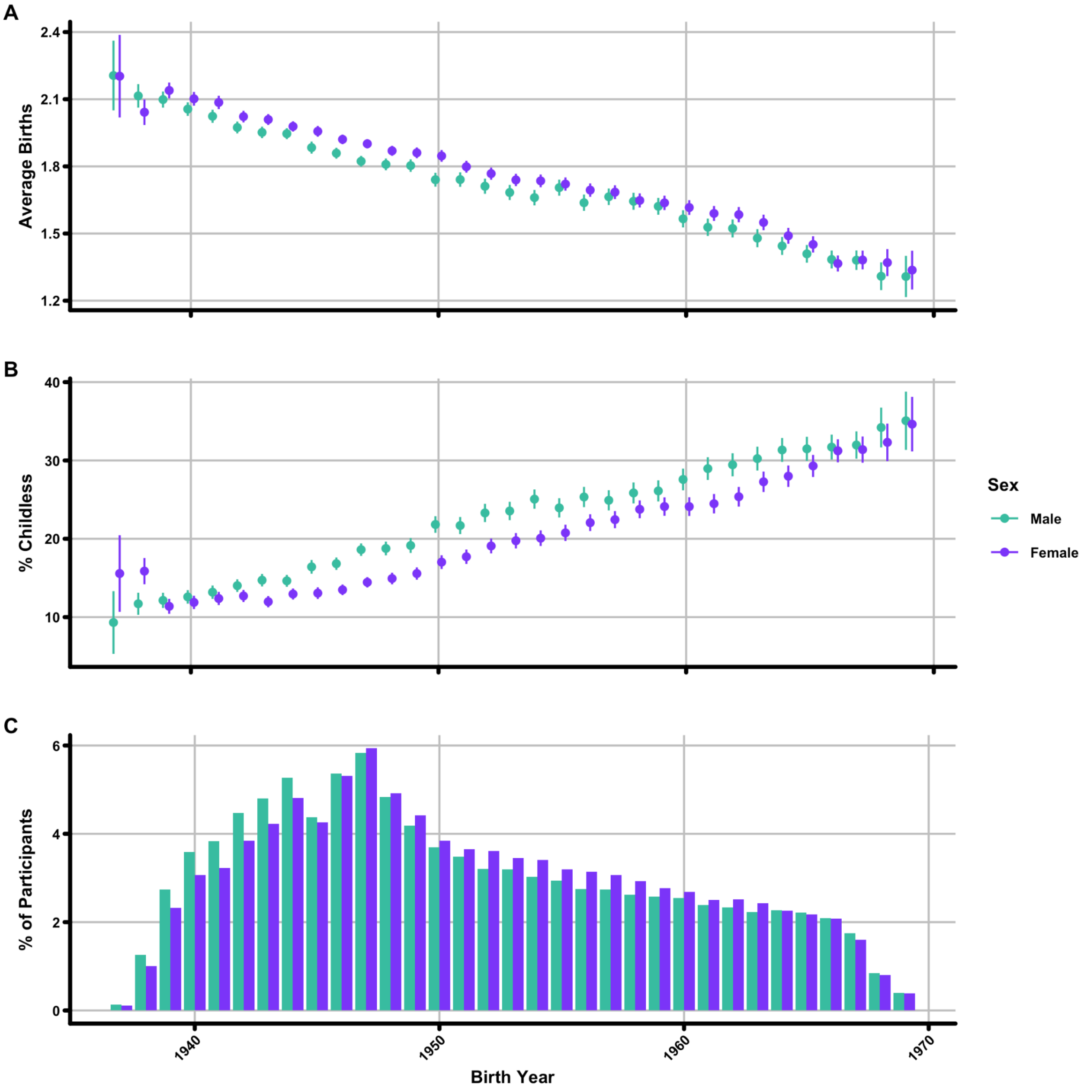


**Supplementary Figure 1. Vital statistics for UK Biobank participants**
(**A**) mean births and (**B**) proportion of individuals without children for all participants in this study. Bins are 1 year birth cohorts. Error bars are 95% confidence intervals on the population proportion. (**C**) Year of birth for all UK Biobank participants included in this study. All plots are separated into females (violet) and males (jade). Only years with greater than 100 individuals born are shown.

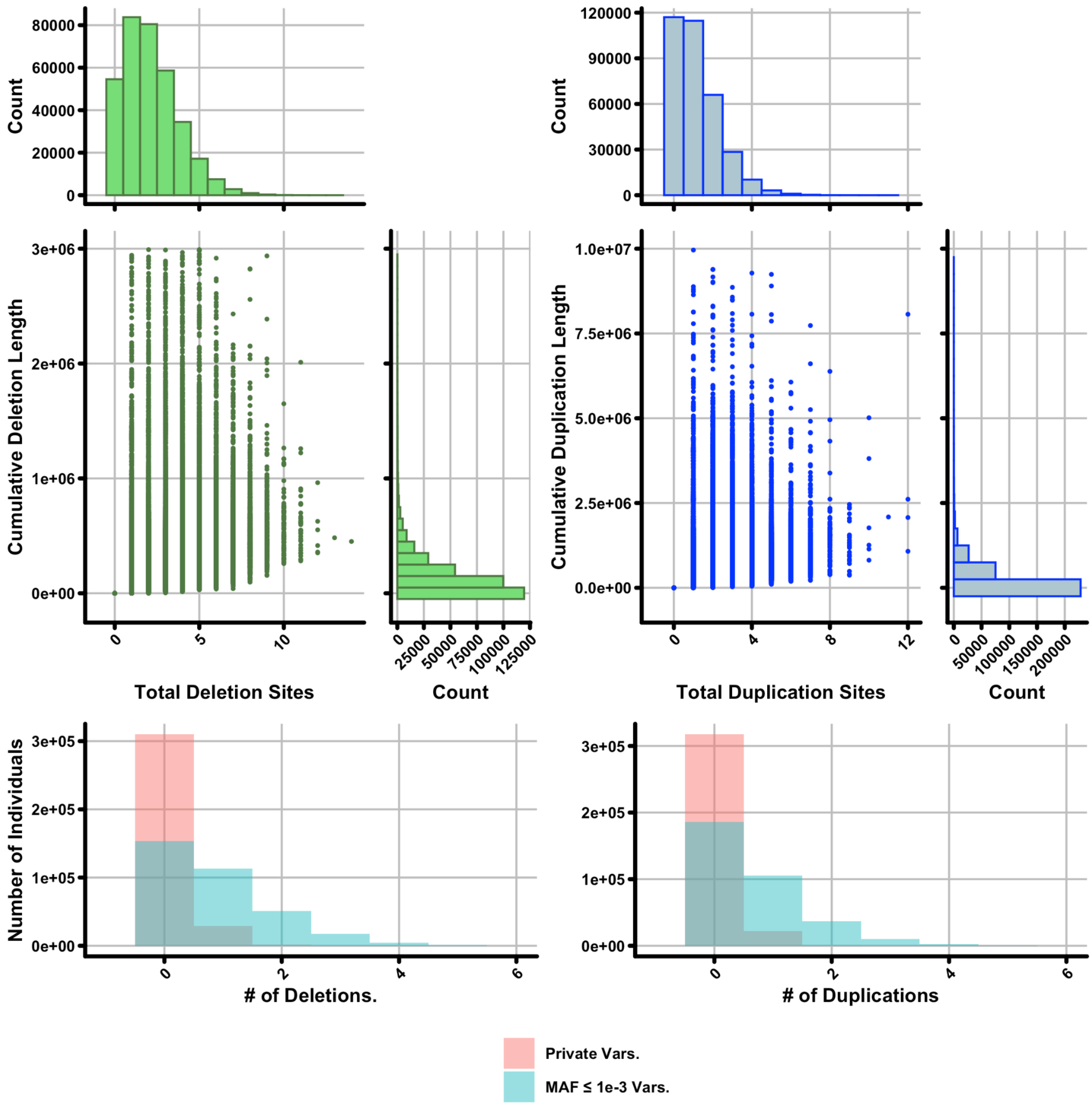


**Supplementary Figure 2. Characteristics of CNVs in the UK Biobank**
Shown are the total number and cumulative length for deletions (green) and duplications (blue) per individual among UK Biobank participants assessed in this study. X-marginal histograms (i.e. those above dot plots), represent the distribution of the number of CNVs per individual in UK Biobank. Y-marginal histograms (i.e. those to the right of dot plots) represent the distribution of cumulative CNV length per individual. Below both plots are per individual totals for private (red) and rare (minor allele frequency < $1x10^{-3}$; blue) variants.

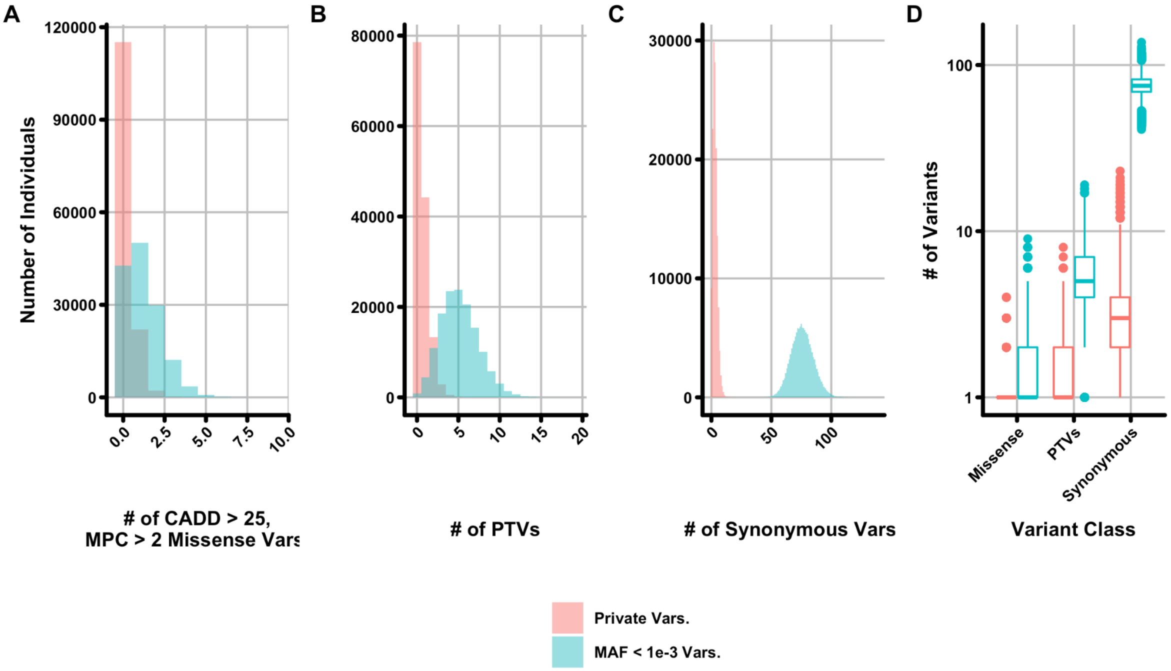


**Supplementary Figure 3. Characteristics of whole exome sequencing-ascertained variants in UK Biobank**

(**A-C**) Total number of (**A**) damaging (CADD > 25 and MPC > 2) missense (**B**) protein-truncating (PTV) and (**C**) synonymous variants per individual among UK Biobank participants with available whole exome sequencing, after applying filtering (see Methods). Shown are per individual totals for private (red) and rare (minor allele frequency < $1x10^{-3}$; blue) variants. (**D**) Comparison between total variants for the three variant classes shown in panels (**A-C**).

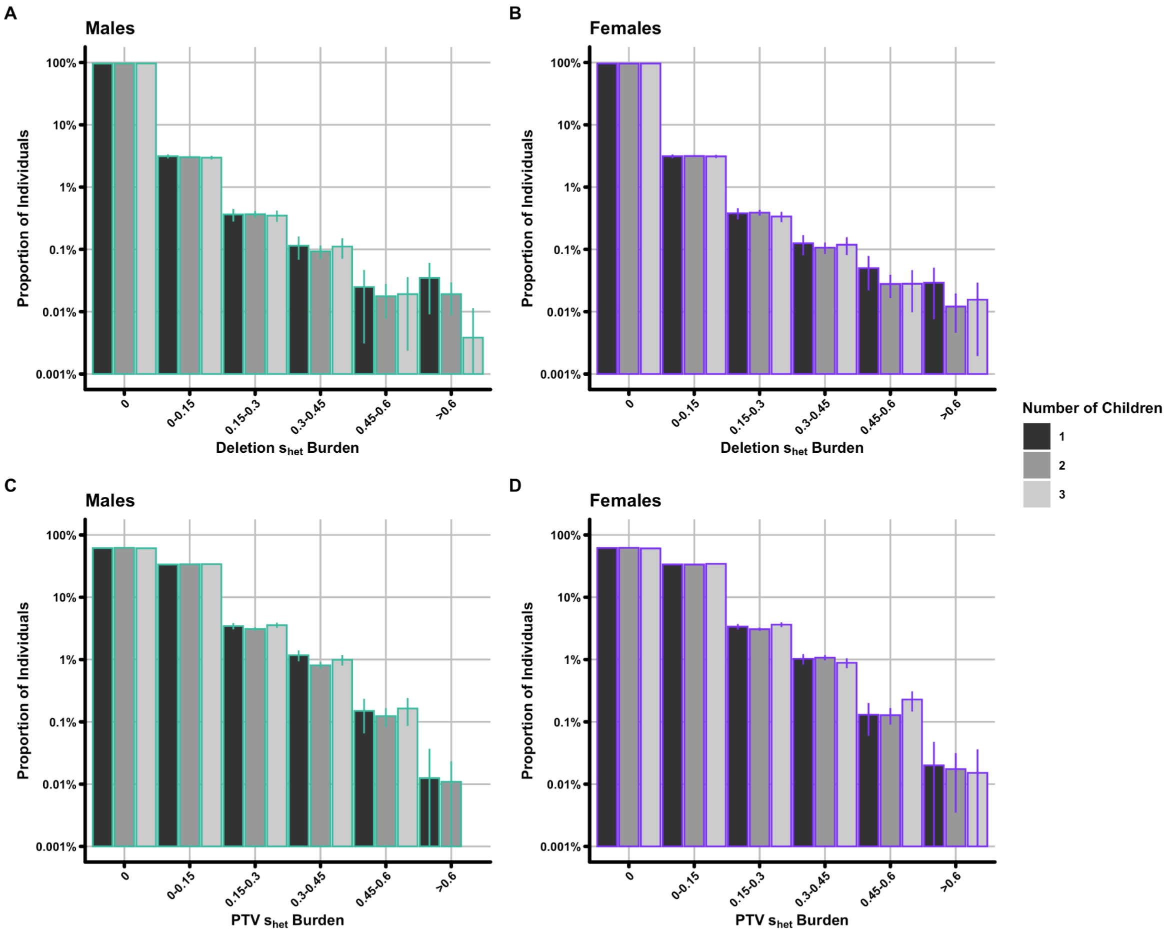


**Supplementary Figure 4. $s_{het}$ distribution stratified by sex and number of children**
Shown are plots displaying the (**A**,**B**) Deletion and (**C**,**D**) PTV $s_{het}$ distributions separately for (**A**,**C**) Males and (**B**,**D**) Females. These plots are identical to main text figure 1C,D, except stratified by the number of children, for individuals with between one to three children (greyscale). Proportions are calculated within the number of children groups, error bars which appear to stop at 0.001 are simply truncated for display purposes. There are no males with three children and PTV $s_{het}$ burden > 0.6; error bars for the other two categories in this bin overlap zero.

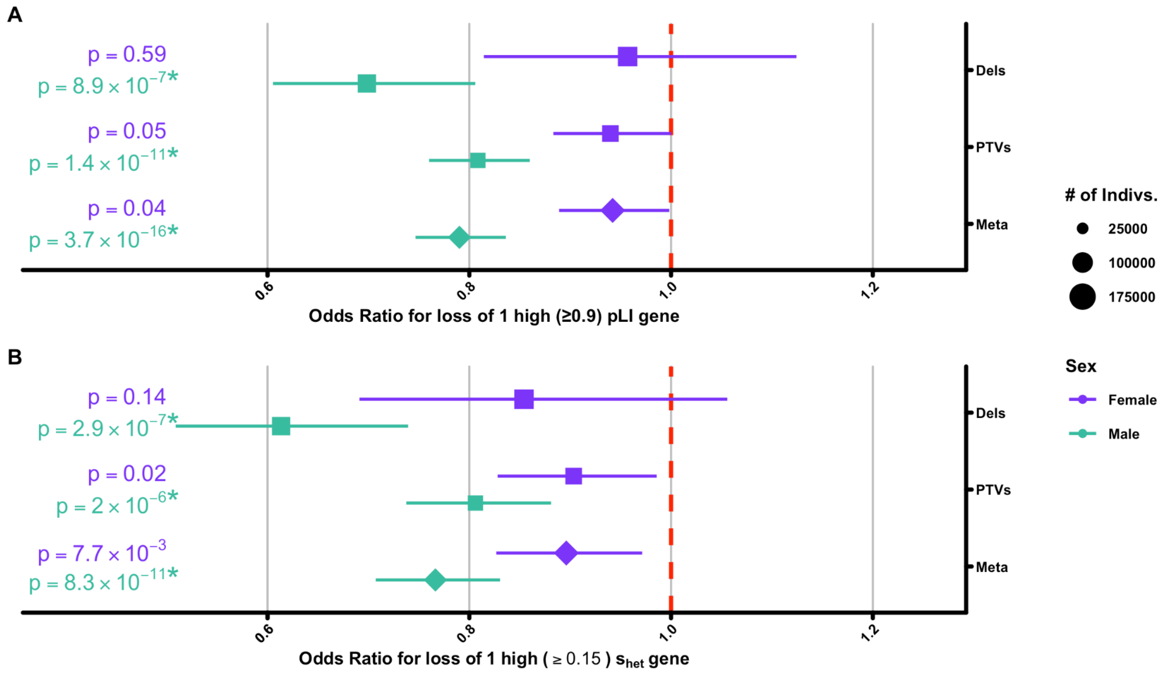


**Supplementary Figure 5. Odds ratio estimates for the association of raw deleterious variant count with having children**

Odds ratio estimates for the loss of (**A**) 1 high pLI (≥0.9)[1] or (**B**) 1 high $s_{het}$ gene[2] on having children, separated into females (violet) and males (jade). Instead of using calculated $s_{het}$ burden as in the main text, we have quantified the total number of genes lost per-individual (see Methods). Asterisks indicate significance after Bonferroni correction for 20 tests ($p < 2.5\text{x}10^{-3}$; Methods).

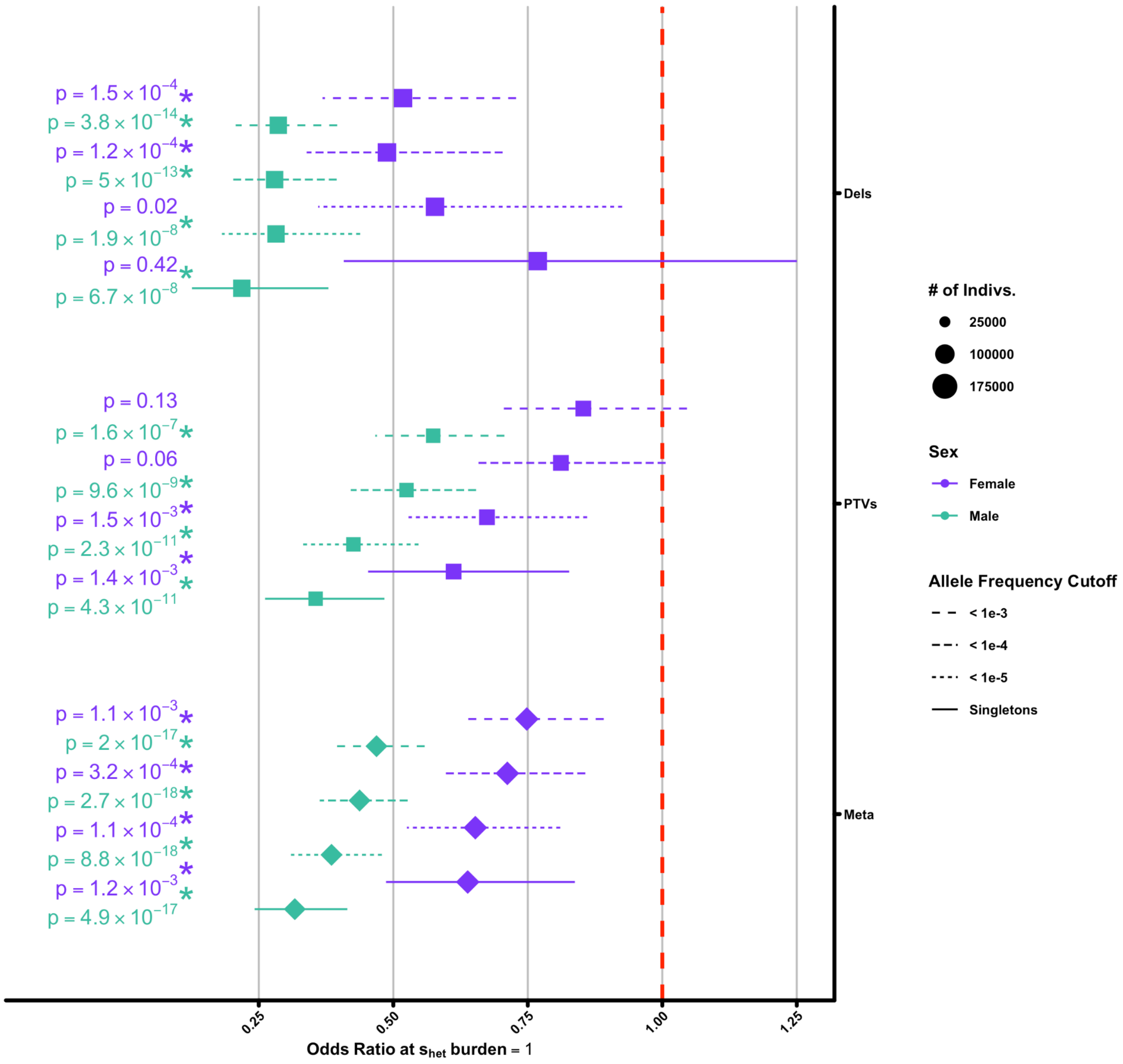


**Supplementary Figure 6. Odds ratio estimate for the association of $s_{het}$ burden with having children when using various rare variant allele frequency cutoffs**

Identical to main text Figure 1B, but showing additional results when $s_{het}$ burden is calculated from deleterious variants with a minor allele frequency of < 1e-3, < 1e-4, or < 1e-5 (indicated by line dash), separated into females (violet) and males (jade). Results marked “Singletons” are identical to those in main text Figure 1B. Asterisks indicate significance after Bonferroni correction for 20 tests ($p < 2.5\times10^{-3}$; Methods).

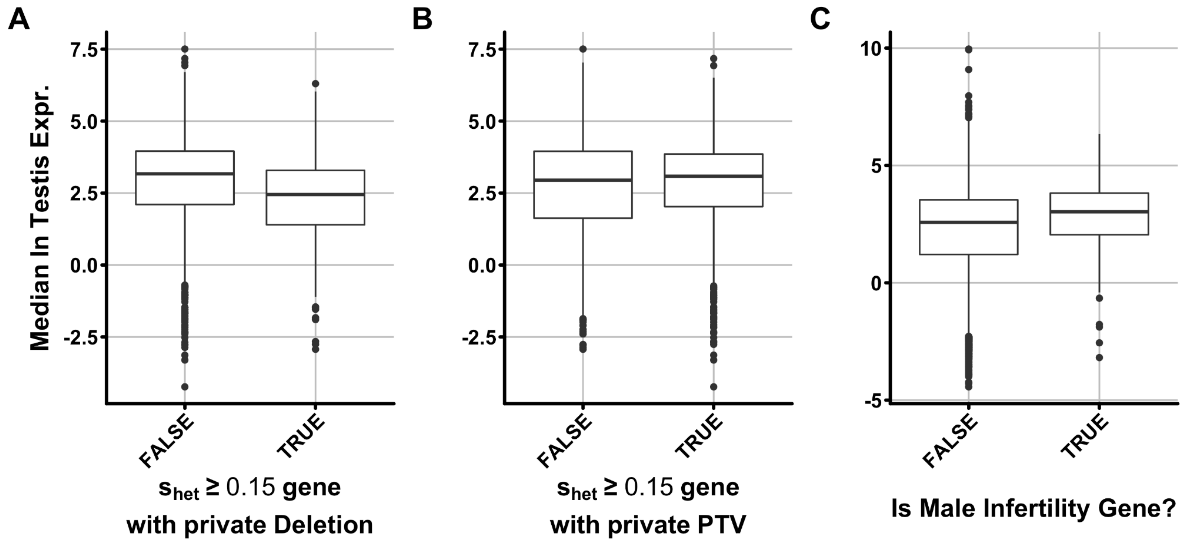


**Supplementary Figure 7. Expression of genes in testis**
(**A-B**) All genes with an $s_{het}$ score ≥ 0.15 subset by whether or not they have any private (**A**) deletions or (**B**) PTVs among individuals in the UK Biobank. (**C**) Genes subset by whether or not they have a relationship with male infertility. The Y-axis for all plots is the median ln(expression testis in TPM) from GTEx.

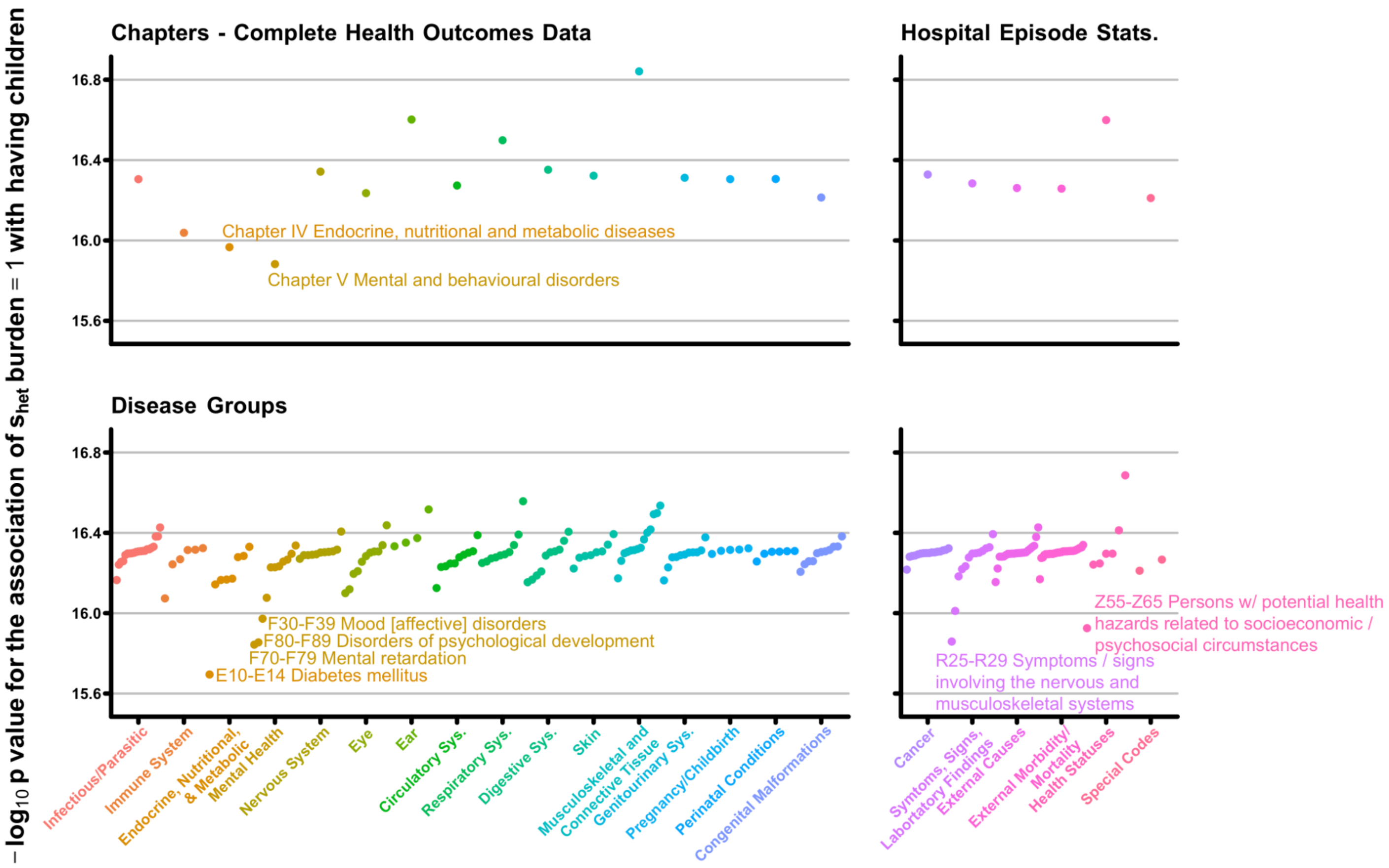


**Supplementary Figure 8. Change in the p. value of the association between $s_{het}$ burden and male reproductive success after inclusion of ICD-10 codes**

Similar to main text Figure 2, plotted are the meta-analysis (Deletion + PTV) odds ratios for the association of individual $s_{het}$ burden (y-axis) on the probability of males having children when corrected for ICD-10 codes across the first two levels of the ICD-10 hierarchy collated from complete health outcomes data (left) or hospital episode statistics (right). Points are colored for the relevant chapter (x-axis) and visual outlier codes are labelled with that code's meaning from ICD-10. Note that text labels do not necessarily represent the full name of a given ICD-10 code. Please see Supplementary Table 2 for a catalogue of all values included in this plot.

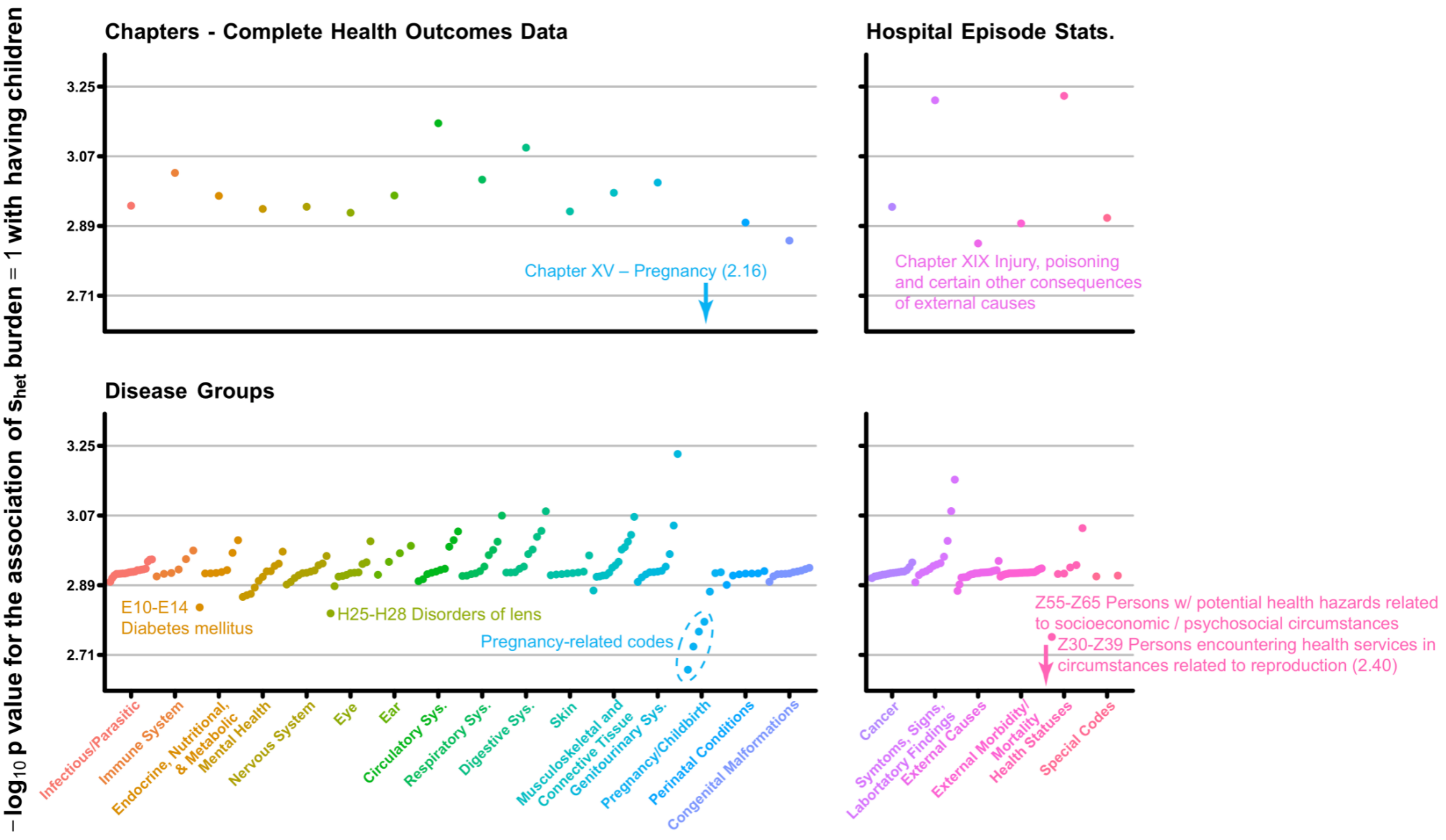


**Supplementary Figure 9. Change in the p. value of the association between $s_{het}$ burden and female reproductive success after inclusion of ICD-10 codes**

This plot is identical to Supplementary Figure 12, except shows associations for females. Plotted are the meta-analysis (Deletion + PTV) odds ratios for the association of individual $s_{het}$ burden (y-axis) on the probability of females having children when corrected for ICD-10 codes across the first two levels of the ICD-10 hierarchy collated from complete health outcomes data (left) or hospital episode statistics (right). Points are colored for the relevant chapter (x-axis) and codes which deviate substantially are labelled with the code meaning from ICD-10. Arrow indicates the point for the indicated code is below the scale of the y-axis and -$\log_{10}$ p value are indicated in parentheses. Note that text labels do not necessarily represent the full name of a given ICD-10 code. Please see Supplementary Table 2 for a catalogue of all values included in this plot.

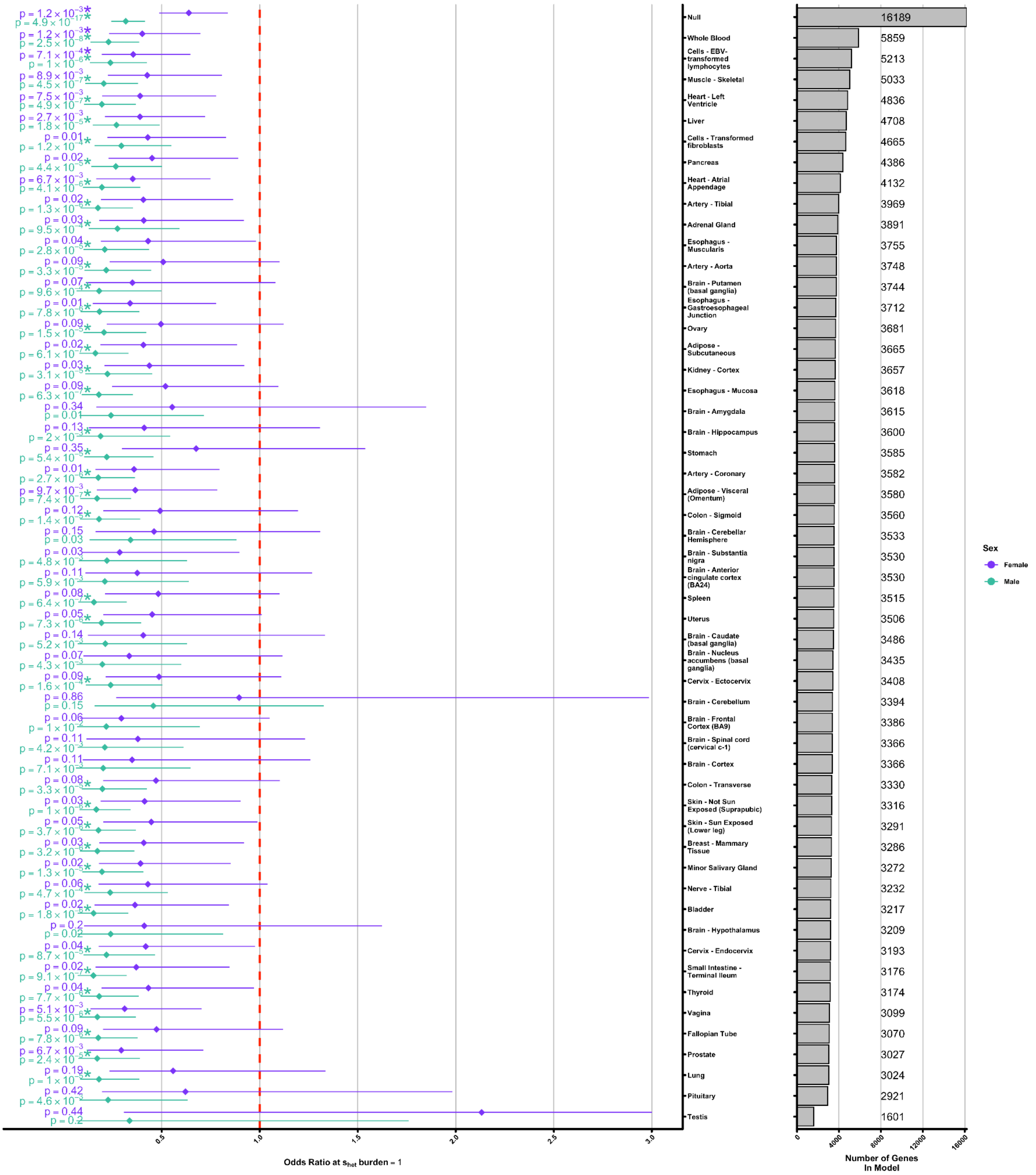


**Supplementary Figure 10. The association of $s_{het}$ burden with childlessness when excluding genes based on expression status in GTEx**

Shown are meta-analysis (Deletions + PTVs) odds ratios separately for females (violet) and males (jade) when excluding all expressed genes (TPM > 0.5) in the tissue listed on the y-axis. Meta-analysis odds ratios are calculated and presented identically to main text Figure 1B, except the total number of individuals is not depicted as the size of the point estimate – the total number of individuals included in all models shown is identical. Per-tissue expression values for all genes were extracted from files provided by GTEx (Methods)[3]. Number of unexpressed genes included in each model when calculating per-individual $s_{het}$ burden are shown as a marginal barplot at right. “Null” is the same as the meta-analysis result depicted in main text Figure 1B for the sake of comparison.

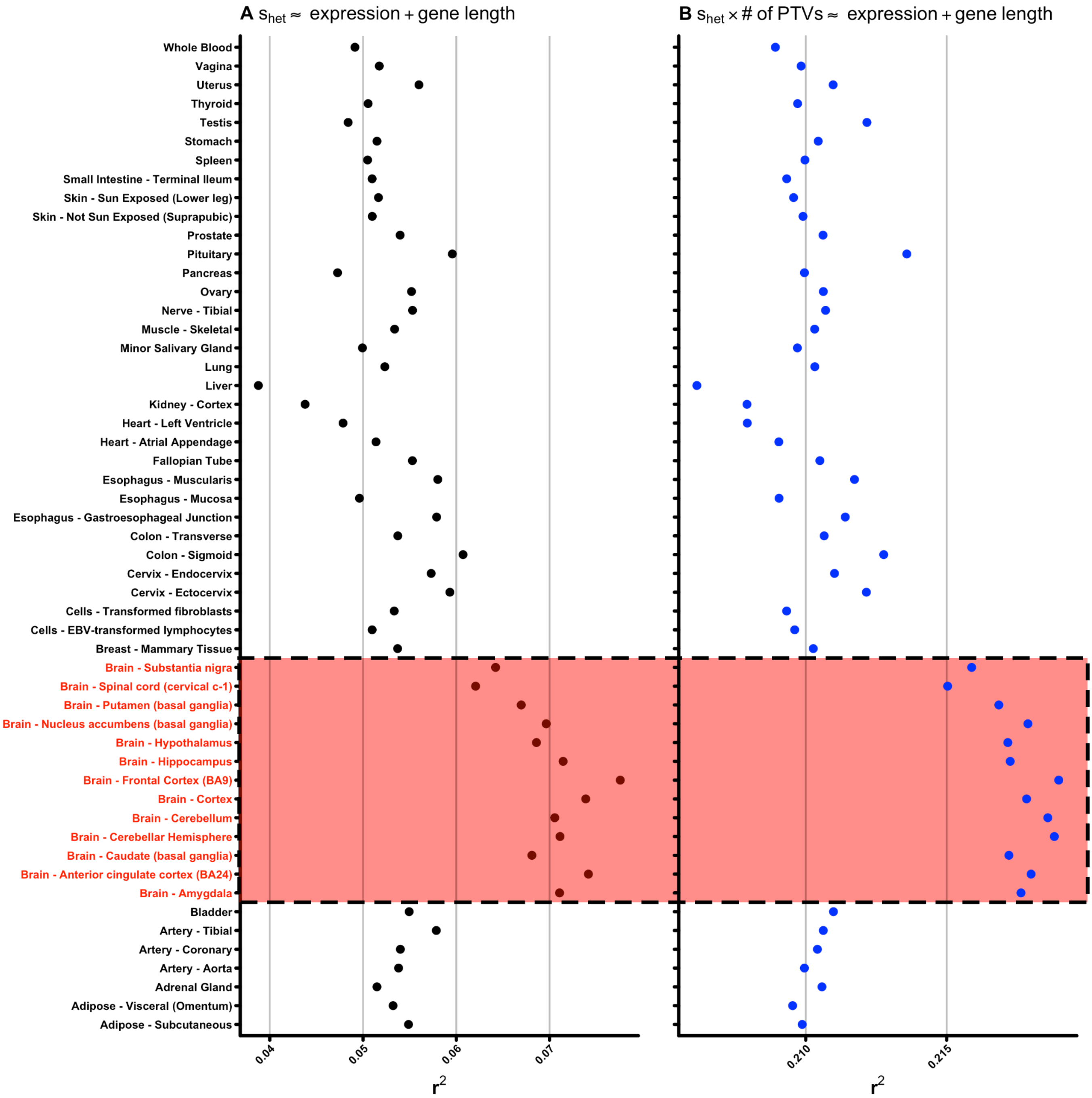


**Supplementary Figure 11. Correlation of per-gene tissue-specific expression with $s_{het}$**
For all 53 tissues sampled as part of the Genotype Tissue Expression (GTEx) study[3], we performed two linear regressions: (**A**) the mean tissue-specific ln(expression) with per-gene $s_{het}$ score corrected for gene coding sequence length and (**B**) the mean tissue-specific ln(expression) with per-gene $s_{het}$ score weighted by the number of private PTVs found per-gene among UK Biobank participants corrected for gene coding sequence length. Displayed above is the variance explained ($r^2$) for each model for each tissue. Brain tissues are highlighted in red.

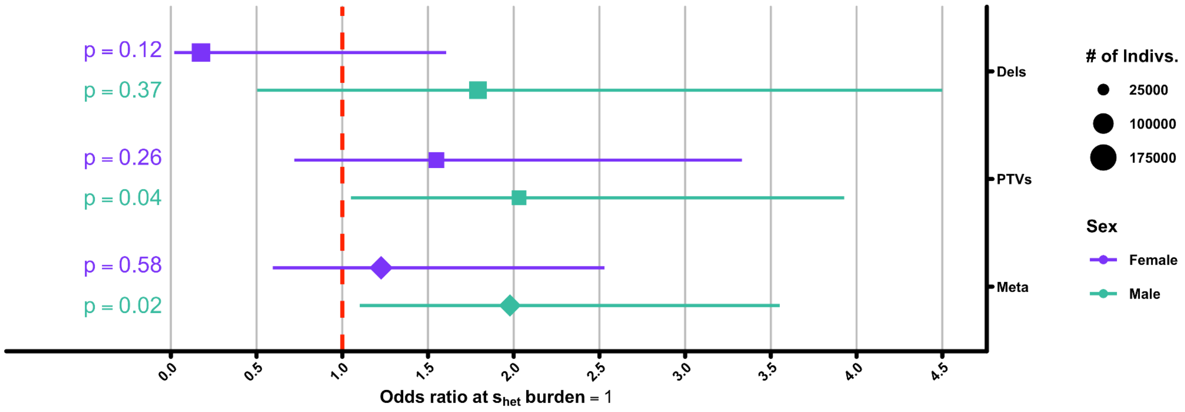


**Supplementary Figure 12. Odds ratio estimates for the association of $s_{het}$ burden with likelihood of engaging in same sex sexual behaviour**

Odds ratio estimates using a logistic regression on the answer to the question 'Have you ever engaged in same-sex sexual behaviour' [1=Yes] as asked during UK Biobank recruitment, separated into females (violet) and males (jade). Asterisks indicate significance after Bonferroni correction for 20 tests ($p < 2.5x10^{-3}$; Methods).

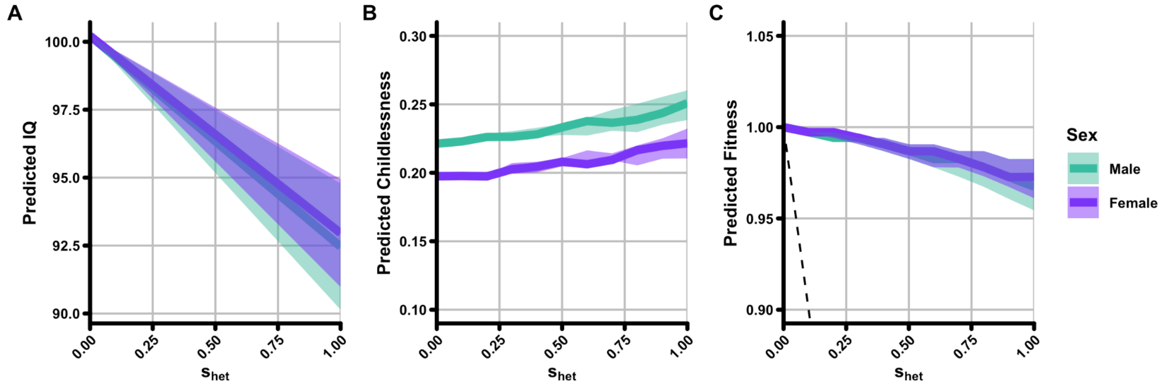


**Supplementary Figure 13. Association of fluid intelligence with fitness**
(**A**) Shown is the predicted mean population IQ score (y-axis) as a factor of individual $s_{het}$ burden based on the logistic model of fluid.intelligence ~ $s_{het}$. (**B**) Predicted childlessness (y-axis) as a function of $s_{het}$ burden if only considering IQ as an explanatory factor. (**C**) Predicted reduction in fitness (y-axis) as a factor of $s_{het}$ burden if only considering IQ as an explanatory factor. For all panels, males (jade) and females (violet) are plotted separately.

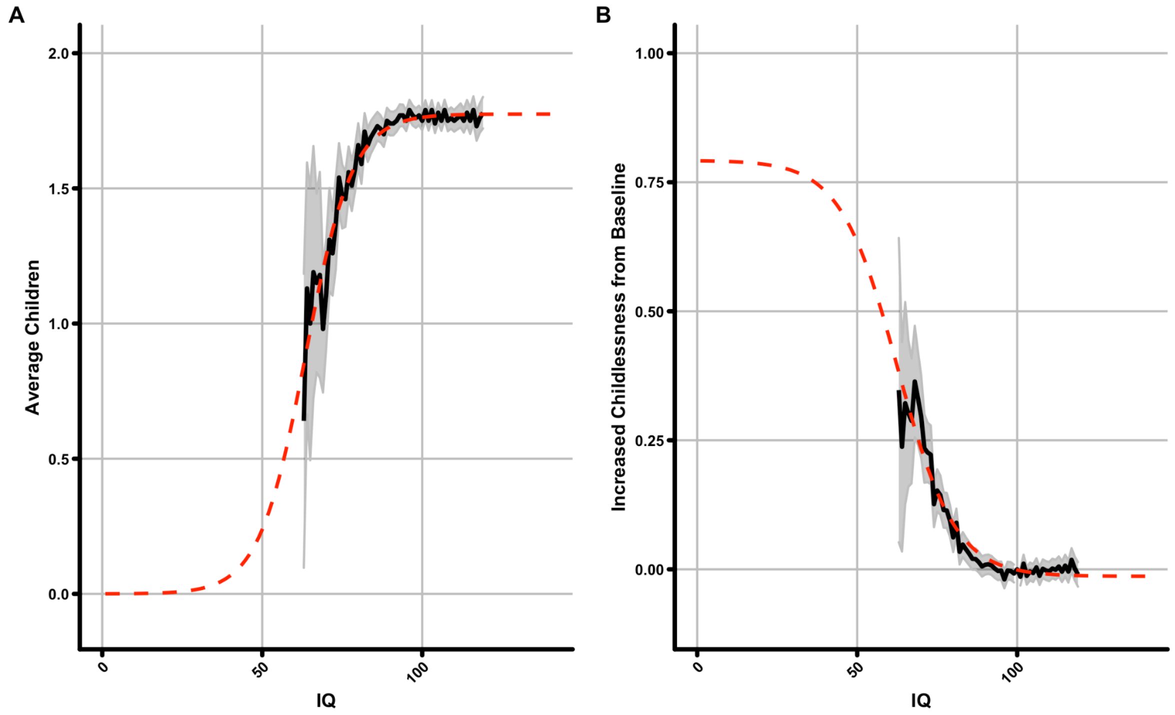


**Supplementary Figure 14. Population level IQ data from Swedish military records**
Shown are actual (black line) and fitted sigmoid curves (red dashed lines) for (**A**) mean number of children and (**B**) increased childlessness from baseline among all Swedish males born between 1965-1967 and tested for IQ as part of military conscription. Grey shading represents the 95% confidence interval from the standard error of the distribution for each IQ bin. See Supplementary Table 5 for raw values used to generate this plot.

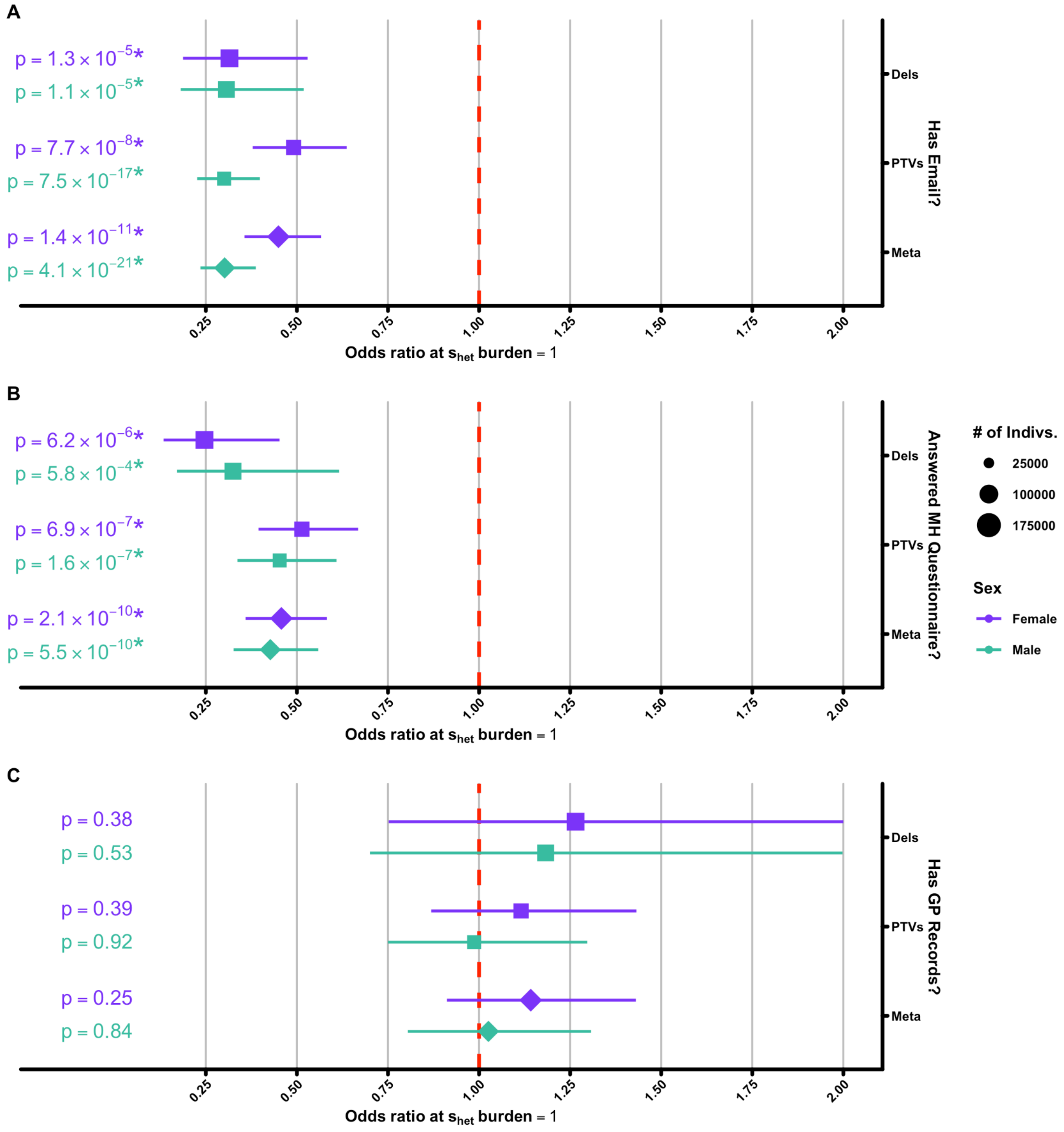


**Supplementary Figure 15. Estimates for the association of $s_{het}$ burden with various recruitment biases in the UK Biobank**

(**A**) Odds ratio estimate for the relationship of individual $s_{het}$ burden with having a functioning email address. (**B**) Odds ratio estimate for the relationship of individual $s_{het}$ burden with whether or not a participant answered the UK Biobank mental health questionnaire[4]. (**C**) Odds ratio estimate for the relationship of individual $s_{het}$ burden on whether or not a participant has general practitioner records. All plots are separated into females (violet) and males (jade). Asterisks indicate significance after Bonferroni correction for 20 tests ($p < 2.5x10^{-3}$; Methods).

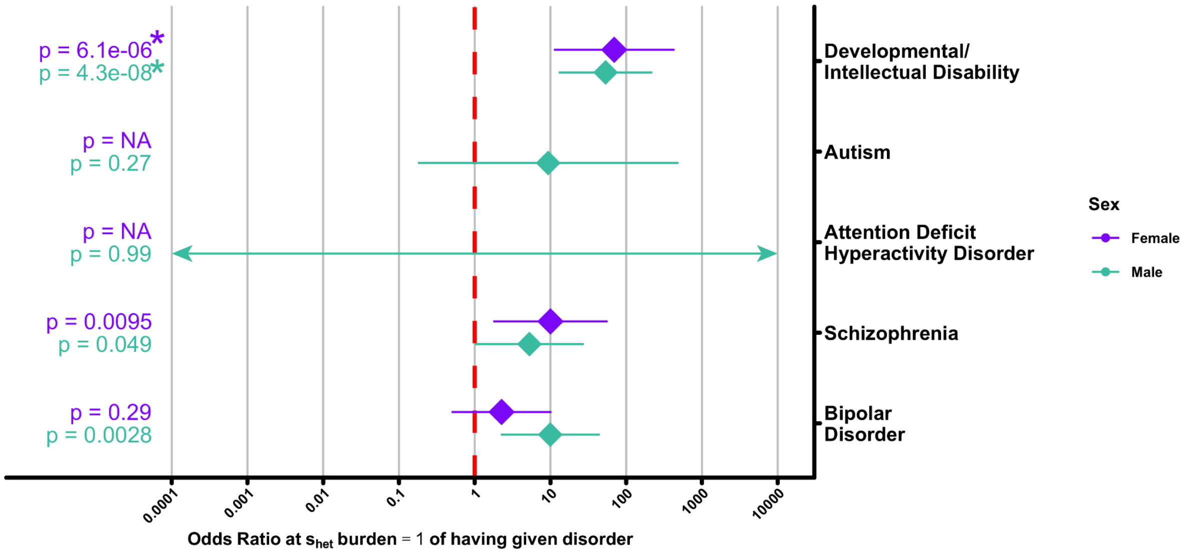


**Supplementary Figure 16. The association of $s_{het}$ burden with individual mental health disorders**

Shown are the odds ratio estimates for $s_{het}$ burden on having a mental health disorders (one of developmental/intellectual disability, autism, attention deficit hyperactivity disorder, schizophrenia, or bipolar disorder; y-axis) from any mental health data source provided by UK Biobank (complete health outcomes, hospital episode statistics, or mental health questionnaire[4]) separated into females (violet) and males (jade). This plot is scaled to show the best view of the majority of disorders – error bars and point estimates for male attention deficit hyperactivity disorder (ADHD) extend beyond the limits of the x-axis (indicated by arrows), likely due to a low number of individuals recruited to UK Biobank with this condition. Additionally, due to the small number of female individuals with both genetic data and a diagnosis of ADHD (n = 42) or ASD (n = 70), our logistic model for these conditions failed to converge (Supplementary Table 4). As such, p value, odds ratio, and standard errors for these data points are not displayed. Asterisks indicate significance after Bonferroni correction for 20 tests ($p < 2.5 \times 10^{-3}$; Methods).

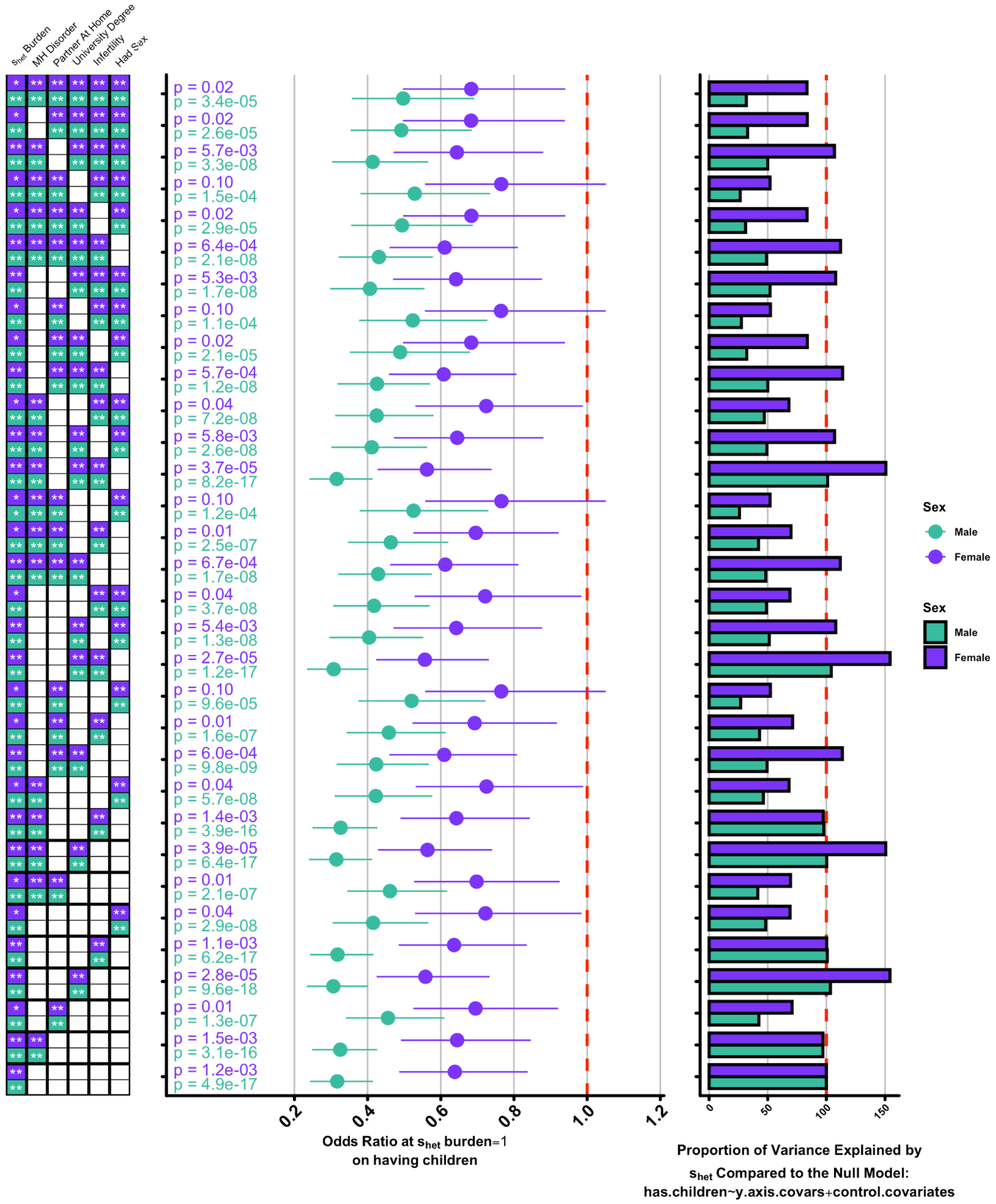


**Supplementary Figure 17. Multiple regression models**

Plotted are the odds ratios for $s_{het}$ burden on childlessness from meta-analyzed (Deletion + PTV) logistic regressions (middle), corrected for a combination of whether or not a study participant has a mental health disorder, a partner at home, a university degree, infertility (as based on CHOD data; Methods), or ever had sex (left); traits included in each model are indicated as coloured boxes (males – jade, females – violet) on the y-axis. Stars within boxes indicate nominal (*; $p < 0.05$) or Bonferroni-corrected (**; $p < 0.0025$) significance level with childlessness for each covariate independently when correcting for PTV $s_{het}$ burden. As indicated to the left, all models include $s_{het}$ burden. The additional bar plot to the right gives the proportion of the variance in childlessness explained by $s_{het}$ burden (for deletions alone) in each model, scaled to the model without any additional covariates (i.e. the model on the bottom of the main plot; see main text Methods).

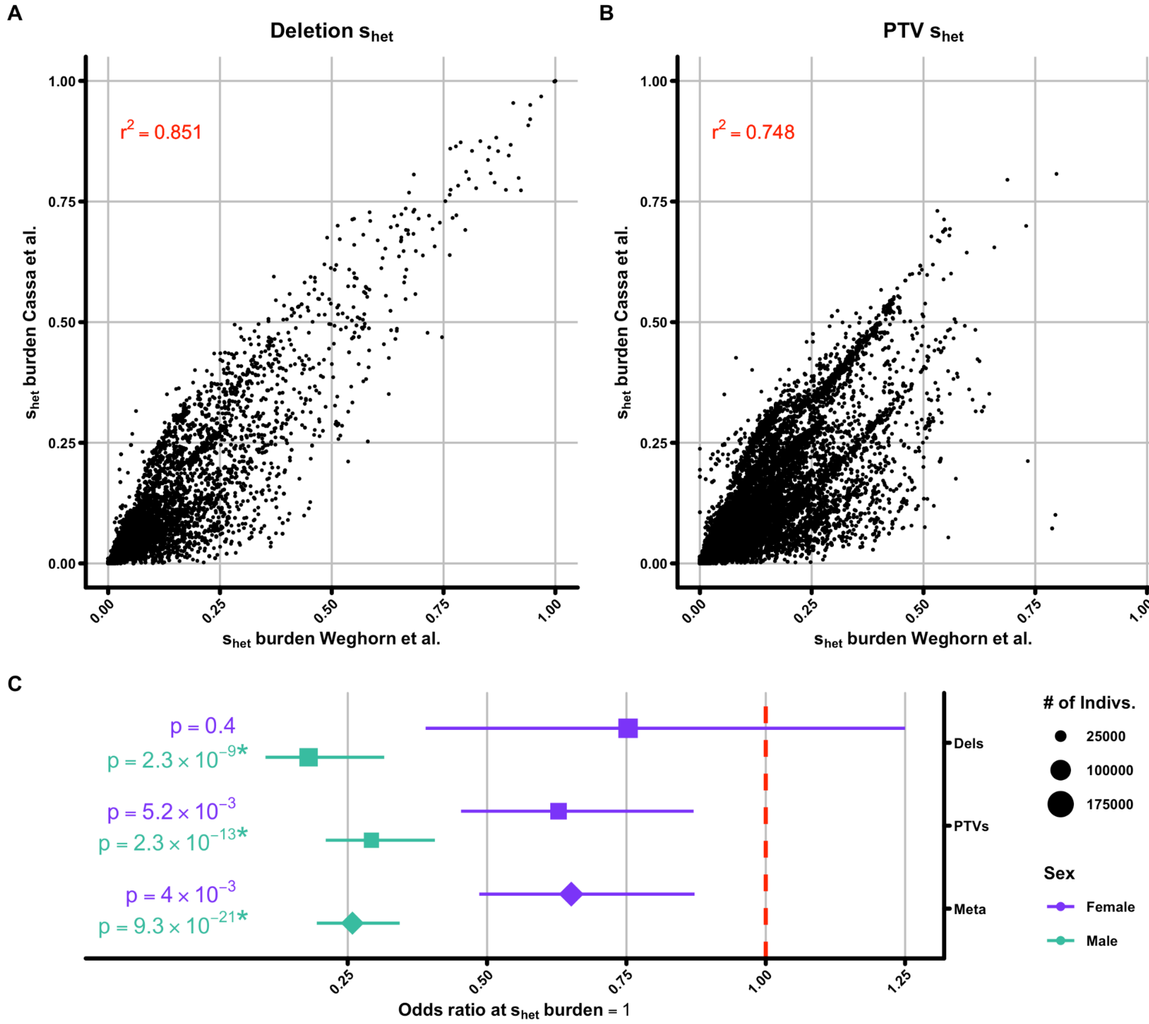


**Supplementary Figure 18. Comparison of $s_{het}$ burden calculated with and without a demographic model**

(**A,B**) Comparison when using per-gene $s_{het}$ scores calculated with[2] and without[5] a demographic model for (**A**) deletions and (**B**) PTVs. Each point represents the relationship between the different $s_{het}$ burden scores for one individual with the correlation between scores shown as red text in each plot. (**C**) The primary result as shown in main text figure 1A except with an $s_{het}$ burden score derived from Cassa et al.[5] rather than Weghorn et al.[2] Asterisks indicate significance after Bonferroni correction for 20 tests ($p < 2.5\times10^{-3}$; Methods).

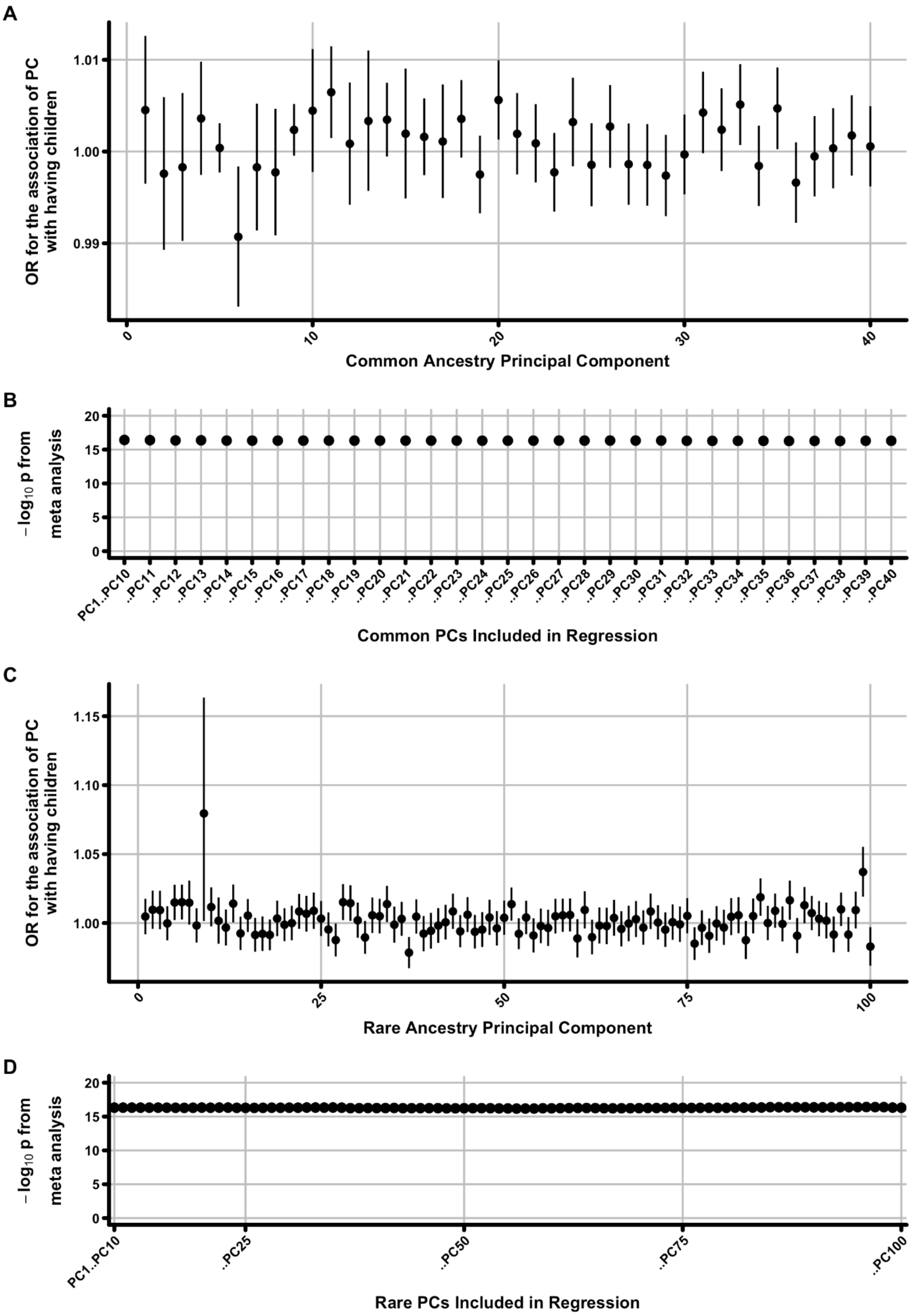


**Supplementary Figure 19. Investigation of the role of ancestry principal components in childlessness**

(**A**) Shown are odds ratios (y-axis) for each of the first 40 ancestry principal components (PCs; x-axis) extracted from a model of has.children ~ age + $age^2$ + birth.cohort + wes.batch + PC1..PC40 for males. Error bars are 95% CIs. (**B**) The meta-analysis $-\log_{10}$ p value for the association of $s_{het}$ burden with male childlessness when controlling for between 10 and 40 ancestry PCs. (**C**, **D**) Identical to plots (**A**) and (**B**), except for the first 100 principal components calculated from IBD sharing over the last 10 generations (Methods)[6].

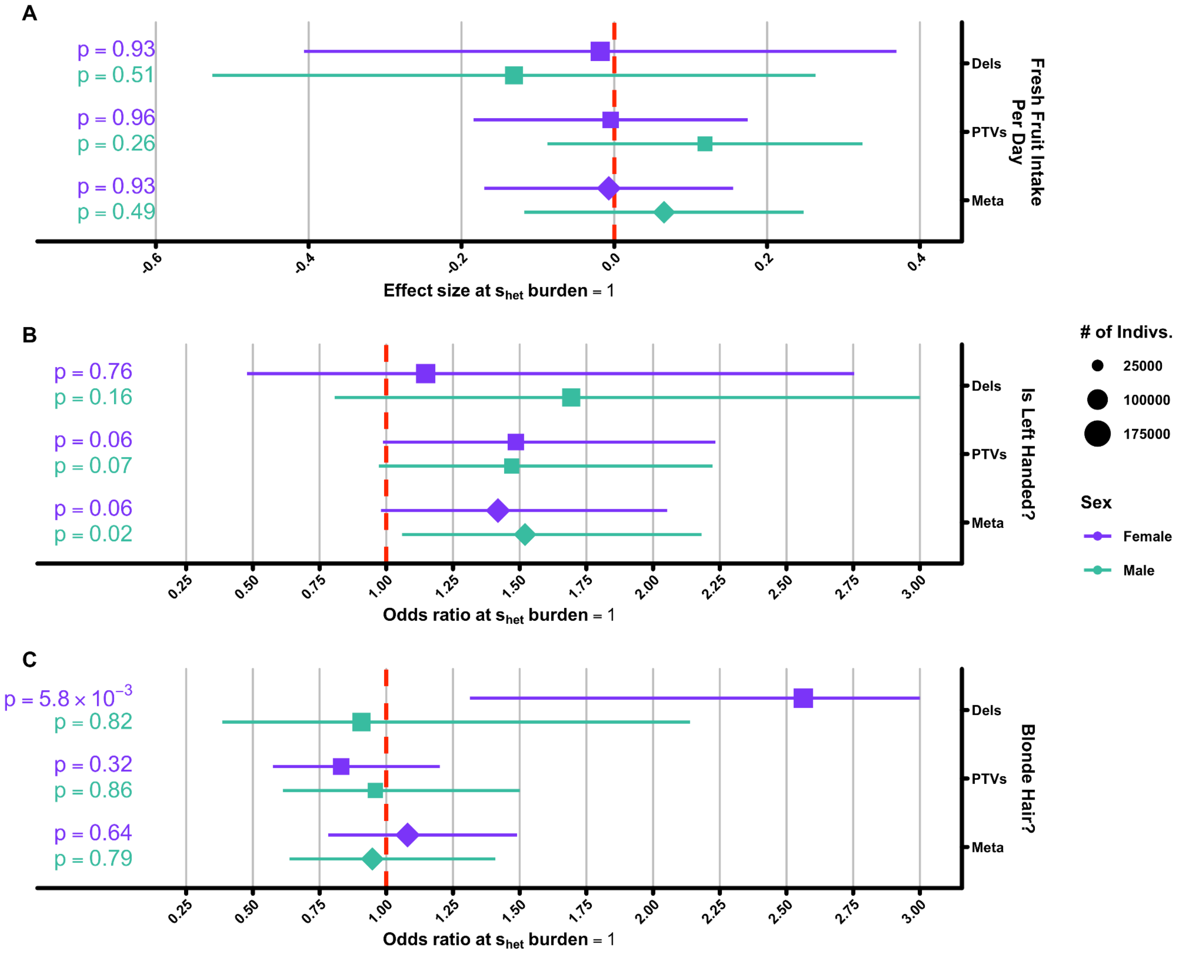


**Supplementary Figure 20. Phenotypes not expected to have any association with $s_{het}$ burden**

Shown are a subset of phenotypes not expected to have a significant relationship with $s_{het}$ burden: (**A**) Total fresh fruit intake per day, (**B**) being left handed, and (**C**) having blonde hair. See Supplementary Table 1 for more details on how these phenotypes were processed. Asterisks indicate significance after Bonferroni correction for 20 tests ($p < 2.5\times10^{-3}$; Methods).

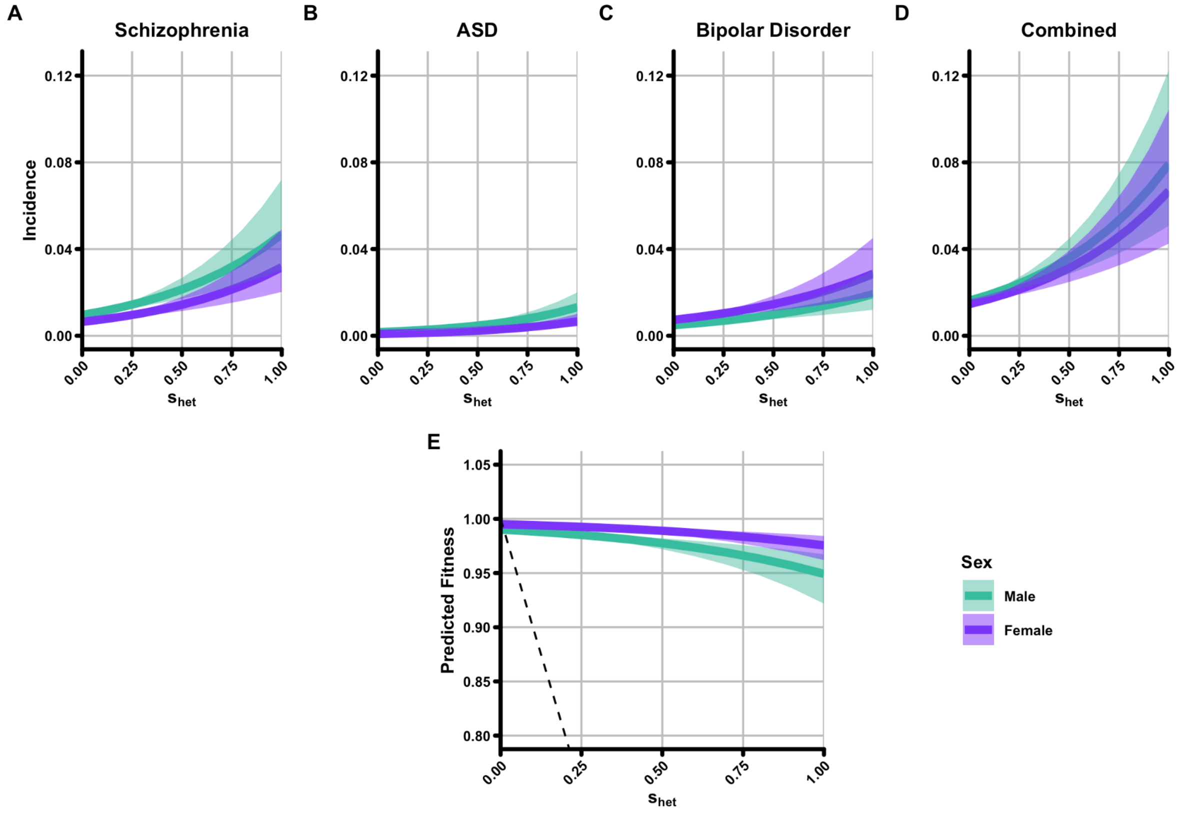


**Supplementary Figure 21. Association of mental health disorders with fitness**
(**A-C**) Predicted incidence (y-axis) of various mental health disorders separately for males (jade) and females (violet) as a factor of individual $s_{het}$ burden (x-axis). Mental health disorders shown are (**A**) schizophrenia, (**B**) autism spectrum disorder (ASD), and (**C**) bipolar disorder. Panel (**D**) represents the summed predicted incidence of all three disorders from panels (**A-C**). (**E**) Predicted contribution of the combined predicted incidence of all mental health disorders to fitness. Note the y-axis in panel (**E**) has been scaled to show a more interpretable representation of the data.

### Supplementary Tables

All supplementary tables have been provided as separate Supplementary Files in Microsoft Excel format and are not included here.

### Supplementary Notes

**Supplementary Note 1. Frequently Asked Questions (FAQs)**
This FAQ document is written to communicate what was found less technically than in the paper, as well as what can and cannot be concluded from the research findings more broadly.

**The Supplementary Note was prepared by:**
*Matthew Hurles, Hilary Martin, Eugene Gardner, Kaitlin Samocha, Kieron Barclay, and Martin Kolk*

***This Supplementary Note has the following sections:***


**Section 1:** Concepts and Terminology

a. DNA, genome, genes, and genetic variants
b. Damaging genetic variants
c. Natural selection
d. Reproductive success
e. Sexual selection
f. Constrained genes
g. Genetic burden and $s_{het}$ burden
h. Association, correlation and how they differ from causality

**Section 2:** Background to the study

a. Who conducted this study? What was the group's overarching goal?
b. What do we already know about the consequences of damaging genetic variation in constrained genes?
c. What do we already know about childlessness?

**Section 3:** Study design and results

a. What did you do in this study and what were the primary results?
b. Why did you use UK Biobank for this research?
c. How could damaging genetic variants affect people's chance of being childless?
d. What do you mean when you say that the effect of damaging genetic variants on childlessness is "mediated by cognitive and behavioural factors"?
e. What is the relative importance of these cognitive and behavioural factors for explaining the association between $s_{het}$ burden and childlessness?
f. Do your findings have implications for health? Could they be used to advance medical research?
g. Why do you think Darwin's theory of sexual selection is relevant to your results?
h. What are the limitations of your study?

**Section 4:** Social and ethical implications of the study

a. Have you found the 'gene for' childlessness?
b. Can you now predict which individuals will be childless based on their burden of damaging genetic variation?
c. If the association of these damaging genetic variants with childlessness is so small at a population level, why are you excited about these results?
d. How are these two results compatible?

e. Many people feel like having children is a choice. Do your results suggest that this isn't true?
f. Is there a risk that this research could lead to discrimination?

#### Concepts and Terminology

***DNA, genome, genes, and genetic variants***

All living organisms carry a template that has instructions on how to make this organism. This template is stored using a complex molecule called deoxyribonucleic acid, or **DNA** for short, which is composed of four smaller molecules: Adenine (A), Thymine (T), Cytosine (C), and Guanine (G). These nucleotides are combined into a specific sequence which is known as the **genome**. In humans, the genome consists of 3.2 billion As, Ts, Cs, and Gs and contains around 19,000 individual **genes**. The As, Ts, Cs, and Gs of genes are arranged in a specific order that instructs the generation of proteins that have diverse functions. These functions can range anywhere from giving us the ability to digest sugar to allowing us to see colour.

While 99.9% of the As, Ts, Cs, and Gs in our genome are identical between any two humans, there are small differences. These are what are known as **genetic variants** and can lead to some of the differences between two humans that we can readily observe such as hair or eye colour.

***Damaging genetic variants***

In some cases, genetic variants can lead to negative outcomes such as diabetes or severe intellectual disability. These **damaging genetic variants** are changes in a person's DNA that we predict will interfere with the function of an encoded protein. In this study we consider two types of damaging genetic variation: (i) deletions, i.e. where a large piece of DNA is removed from a specific location in a genome, and (ii) protein-truncating variants (PTVs), i.e. where there is a small change in the DNA sequence that is predicted to truncate the encoded protein. For simplicity in this FAQ, we will not use the terms "deletion" or "protein truncating variant" and just refer to both as damaging genetic variants or collectively as damaging genetic variation.

***Natural selection***

**Natural selection** is the process by which some individuals in a population that are better adapted to their environment have better chances of surviving and producing (more) offspring. Thereby, they also propagate genetic variation that helps them to better adapt to their environment. In turn, this leads to the depletion of genetic variation from the population that causes poorer adaptation to the environment. Over many generations, natural selection can result in changes in traits within a population. It is the key mechanism of evolution.

##### *Reproductive success*

The term **reproductive success** is used in biology to describe the number of biological offspring an individual has in their lifetime. Reproductive success of an individual depends on them a) surviving to reproductive age, b) being able to biologically reproduce (for example, being able to produce viable sperm or eggs), and c) finding a partner with whom to have children. Reproductive success is directly related to natural selection. Individuals who are better adapted to their environment tend to have more children and thus pass their genetic variants on to the next generation.

##### *Sexual selection*

**Sexual selection** is a type of natural selection in which members of one biological sex choose partners of the other sex with particular characteristics to have offspring with ("mate choice"), or compete with members of the same sex for access to partners of the opposite sex ("mate competition"). Hence, sexual selection directly impacts reproductive success within a species, rather than affecting the survival of the individual. The two mechanisms of sexual selection described above (mate choice and mate competition) represent selection pressures that disproportionately affect one sex more than the other. Thus, genetic variants or traits that do not impact survival but that impact reproductive success differently in males versus females may be doing so via sexual selection.

##### *Constrained genes*

New genetic variants arise through the process of mutation. Each child will have a small number of changes in their genome that are different from their mother and father. The majority of these new genetic variants will have no effect on the child, but in a small number of cases, these variants will be "damaging" (to the gene function) and potentially lead to disease.

Previous studies that have sequenced the DNA of thousands of people have found that some of the 19,000 genes in the human genome contain many fewer damaging genetic variants than we would expect, based on what we know about the process of mutation[1]. We call these **"constrained genes"**. It is thought that natural selection acting on damaging genetic variation in these genes is stronger than on other genes and that this leads to a relative depletion of damaging genetic variants in constrained genes. For example, if both of the following are true:

1. Damaging genetic variants in Gene A cause (or increases the risk of) Disease X
2. Disease X decreases the likelihood that a given person will have children

then individuals with Disease X will be less likely to pass on damaging genetic variation in Gene A via sexual reproduction. Scientists will thus observe this relationship as a depletion of damaging genetic variation in Gene A. In other words, damaging genetic variants in these genes confer (or increase the risk of) traits which reduce reproductive success in humans.

For many constrained genes, we do not know why damaging genetic variants reduce reproductive success. Potential mechanisms are that these variants are selected against because they cause a severe disorder which is fatal during childhood, because they cause infertility, or because they alter other characteristics (e.g. behaviours) that make individuals less likely to have children.

##### *Genetic burden and $s_{het}$ burden*

To be able to test whether damaging genetic variation is associated with reproductive success, we needed a metric that quantifies the number of damaging genetic variants in each person's genome. In genetics, this quantification is often referred to as an individual's **genetic burden**. In this paper, we calculated a metric for each person that summarises how much damaging genetic variation they carry across their entire genome, and termed it their **$s_{het}$ burden**. The details of this calculation are more complicated than simply counting the number of damaging genetic variants, but it is similar to the number of damaging genetic variants within constrained genes that an individual carries. In this FAQ and the paper, when we say that an "individual/person has a high $s_{het}$ burden", it roughly means that the person's genome contains at least one damaging genetic variant that disrupts a constrained gene.

##### *Association, correlation and how they differ from causality*

In statistics, **association** is used almost interchangeably with **correlation**. Two events or traits are said to be 'associated' if a change in one tends to be accompanied by a change in the other.

Importantly, association does not necessarily imply causation. If two variables (let's call them A and B) are associated (or correlated), this could imply one of three things:

- A causes or contributes to B – B is the result of, or partly due to, the occurrence of A
- B causes or contributes to A – A is the result of, or partly due to, the occurrence of B
- a third variable, C, causes or contributes to both A and B. In this case, C is sometimes termed a 'confounder'.

In the field of genetics, researchers often report that a particular genetic variant is *associated* with a certain trait. Because an individual's DNA sequence is set at their conception and does not change throughout their life it is not possible for the trait to *cause or contribute to* the genetic variant. Thus, if the genetic study has been conducted in such a way that one can control for or remove the effect of confounders, it is likely that the genetic variant causes or contributes to the trait; however, it can be difficult to definitively prove causation because it is i) difficult to control for all possible confounders or ii) prove that all confounders have been controlled for.

#### Background to the study

***Who conducted this study? What was the group's overarching goal?***

This study was conducted collaboratively between an international, multi-institution group of geneticists and demographic researchers.

The goal of the study was to better understand why some genes are constrained. We assume that this constraint against damaging genetic variants arises because they are associated with traits which reduce reproductive success and so natural selection removes them from the population. For a minority of constrained genes (about a third), we think that this happens because these damaging genetic variants cause severe, often life-limiting disorders. Patients with these disorders are less likely to survive to adulthood, and are less likely to have children and to pass on these genetic variants to the next generation. Neurodevelopmental disorders represent the largest subset of such severe disorders; however, the majority (about two-thirds) of constrained genes have not yet been linked to such disorders. Thus, we don't know why natural selection is removing damaging genetic variants in these genes from the population, and the goal of this study was to investigate this observation. We hoped that, by uncovering this, we might be able to use this information in the future to help discover more genes in which damaging genetic variation is linked to severe, often life-limiting disorders. Many patients with such disorders do not currently receive a genetic diagnosis after genetic testing. There is strong evidence that some of these undiagnosed patients have damaging genetic variants in genes that are causing (or increasing risk of) their disorder, but have yet to be robustly linked to disease. It has been estimated that there may be more than a thousand genes in which we have not yet linked damaging genetic variation to such disorders[7]. These novel disorders will be enriched among constrained genes that have not yet been linked to disease. If we were to identify that a subset of constrained genes were likely linked to reduced reproductive success through causing (or increasing risk of) infertility, rather than a severe, life-limiting disorder, then this could allow us to focus our attention on a smaller subset of constrained genes that would be more likely to be linked to a severe, life-limiting disorder.

We did not initially set out to study the genetic and non-genetic factors that influence childlessness or reproductive success.

***What do we already know about the consequences of damaging genetic variation in constrained genes?***

Damaging genetic variation in many constrained genes has previously been shown to cause, or greatly increase the risk of, specific rare genetic diseases[8]. Many of these rare genetic diseases are severe and life-long, and would be expected to lead to earlier death, especially before the advent of modern medicine.

In addition, damaging genetic variation in constrained genes has also been previously associated with increased risk of developing several relatively more common psychiatric disorders such as autism, schizophrenia, and ADHD and has also been associated with reduced educational attainment (measured as the number of years an individual spends in education) and shorter height[8]. On the other hand, damaging genetic variation was not found to be associated with blood pressure, BMI or cholesterol levels, or with risk of various common cardiometabolic (e.g. type 2 diabetes) or auto-immune diseases (e.g. inflammatory bowel disease)[8].

***What do we already know about childlessness?***

The proportion of people who do not have biological children of their own varies over time, and from country to country. In the UK, approximately 20% of adults over age 50 are childless. Childlessness can be voluntary or involuntary. In a recently published survey of childless UK adults, 28% of men and 31% of women said they did not want children, 7% of men and 15% of women said it was because either they or their partner were infertile, and 23% of men and 19% of women said it was because they had never met the right person[9].

Additionally, many health and socio-demographic factors have been found to be associated with increased likelihood of being childless in contemporary studies. These factors have often been found to be different between males and females. For example, several psychiatric disorders such as schizophrenia are associated with increased likelihood of being childless, but more so in men than in women[10]. Among socio-demographic factors, low socioeconomic status has been more strongly associated with childlessness in males than females, higher educational attainment has been more strongly associated with childlessness in females than males[11], and low performance on an intelligence test has been found to be associated with increased childlessness in males[12].

Previous genetic studies of reproductive success, which include twin studies and those studying genetic variation directly, have shown that both male and female reproductive success is partly heritable (meaning reproductive success has both genetic and environmental components). In the Swedish TwinGene study, a collection of both identical and non-identical twins born in Sweden from 1911-1958, ~13% and ~14% of women and men, respectively, were childless[13]. Importantly, though, genetic variation was found to only explain about half the variation in childlessness in this cohort[13]. It is likely that the relative contribution of genetic and environmental factors to childlessness, and their specific nature, vary across cultures and time.

#### Study design and results

***What did you do in this study and what were the primary results?***

Using data provided by the UK Biobank study, we set out to investigate the consequences of damaging genetic variants in constrained genes to understand why such genes are depleted for damaging genetic variation in humans. We performed this work to assess our primary hypothesis that damaging genetic variation in constrained genes leads to reduced reproductive success, which in turn leads natural selection to remove damaging genetic variation from the population.

To investigate whether damaging genetic variation was associated with lower reproductive success, as we hypothesized, we first tested whether $s_{het}$ burden (for a description of $s_{het}$ burden, see Genetic burden and $s_{het}$ burden) was associated with the number of children that UK Biobank participants had. We found that it was associated with males having an average of 0.25 fewer children (but not females). To understand whether this finding was due to a decrease in the number of children individuals have or due to individuals not having any children at all (i.e. childlessness) we then tested whether $s_{het}$ burden was associated with childlessness. We found that $s_{het}$ burden was associated with a higher chance of being childless in both males and females, but again much more so in males, but we did not find an association with the number of children (in those who had at least one child).

We then investigated why higher $s_{het}$ burden might be associated with childlessness, testing three hypotheses:

1. Whether $s_{het}$ burden was associated with an inability to produce viable eggs or sperm (i.e. infertility) in males or females. We found that it was not.

2. Whether $s_{het}$ burden might be associated with childlessness via an impact on various medical conditions. In this analysis, we controlled for the presence of all diseases, disorders, and special medical codings listed in the WHO International Classification of Diseases v10 (ICD-10) when testing the association between $s_{het}$ burden and childlessness. We found that none of them made a large difference to the association, which indicated to us that it was unlikely that $s_{het}$ burden was affecting childlessness by influencing risk of health conditions that themselves influence reproductive success.

3. Whether $s_{het}$ burden was associated with various other traits that have been previously associated with childlessness[10,11,14]. We showed that higher $s_{het}$ burden was associated with increased risk of having any mental health disorder, weaker performance on an intelligence test, lower chance of having a university degree, lower income and lower socioeconomic status. We also found that it was associated with a lower chance of living with a partner and of ever having had sex. We found that these cognitive and behavioural factors explained 68% of the association between $s_{het}$ burden and childlessness in males, and about 16% in females.

By comparing the associations between $s_{het}$ burden and these various factors between males and females (see below for more details), we concluded that the association between $s_{het}$ burden and childlessness in males was likely to be due to its effect on cognitive and behavioural traits that made males less likely to find a partner to have children with. Thus, our results are consistent with the hypothesis that these damaging genetic variants in constrained genes are likely under negative selection partly due to sexual selection, and specifically, due to mate choice.

***Why did you use UK Biobank for this research?***

UK Biobank is a study funded by several public agencies and charities in the United Kingdom to provide a data resource of about 500,000 people that enables studies of human health[15,16]. We focused on studying the participants in the UK Biobank for three main reasons:

1. The large size of the UK Biobank and availability of genetic data for a large subset of the cohort gave us the ability to detect even very subtle differences between groups of individuals. Here, this means looking for differences in childlessness between individuals with different levels of damaging genetic variation. In statistics and genetics, we call the ability to detect such differences "power", and the large number of individuals within the UK Biobank maximised our power to detect meaningful differences between individuals.

2. Most UK Biobank participants are over the age of 40, and past the age people normally have children. Thus, we could be reasonably certain that the number of children reported by the participants represented the final number of children they would have in their lifetime. Additionally, we could be confident that any association between $s_{het}$ burden and childlessness in this cohort was not because $s_{het}$ burden was affecting an individual's chances of surviving long enough to have children.

3. The availability of a large amount of data for UK Biobank participants, including both health-related data (such as hospital visits) and demographic information, enabled a broad range of analyses exploring the potential contribution of different medical and non-medical factors to any genetic associations that we identified.

***How could damaging genetic variants affect people's chance of being childless?***

To try to understand how damaging genetic variation may be linked to childlessness, we examined the association of $s_{het}$ burden with traits and outcomes that have previously been linked to childlessness. Results for these traits fell into three categories:

1. Traits with which $s_{het}$ burden showed no association:
    - Diagnosis of infertility as reported by health care records
    - Self-reported same-sex sexual behaviour

2. Traits where $s_{het}$ burden had an equal association in both sexes. We found that people with a higher $s_{het}$ burden tended to:
    - be more likely to have mental health disorders
    - perform less well on intelligence tests
    - be less likely to have a university degree
    - have lower household income
    - be more likely to live in a deprived/poor area
    - be less likely to ever have had sex

3. Traits where $s_{het}$ burden had a different association in the two sexes:
    - Men with a higher $s_{het}$ burden were less likely than women to report that they live with a partner at home

Together, these findings led us to suspect that the link between damaging genetic variation and childlessness is mediated by cognitive and behavioural factors which may affect the likelihood of finding a partner and opportunities to have children.

***What do you mean when you say that the effect of damaging genetic variants on childlessness is "mediated by cognitive and behavioural factors"?***

In thinking about which traits might be affected by damaging genetic variation that could, in turn, have an impact on whether a person is childless or not, we considered two main alternatives: (i) that damaging genetic variation affects the ability to form viable sperm and eggs, or (ii) that damaging genetic variation affects the likelihood of forming reproductive partnerships. We found no evidence for the former but consistent associations for the latter as demonstrated by the summarised results discussed in the previous question of this FAQ.

We use "cognitive and behavioural factors" as a general term to capture diverse aspects of brain function that may be associated with the likelihood that individuals form reproductive partnerships. How people behave, and how others respond to those behaviours are important factors in how reproductive partnerships are formed.

We do not know the specific cognitive and behavioural factors that might be mediating the association between damaging genetic variation and childlessness. It is likely that there is not a single behavioural or cognitive trait that is playing a role, but rather a combined effect of several different traits, possibly including but not limited to the ones we investigated here. There are important behavioural traits that are known to be associated with reproductive success (e.g. personality traits), which we have not been able to evaluate in this study. Therefore, it would not be appropriate to speculate on the relative importance of the different cognitive and behavioural factors that could potentially be involved.

***What is the relative importance of these cognitive and behavioural factors for explaining the association between $s_{het}$ burden and childlessness?***

We found that not having a partner at home was the most important factor in the association between damaging genetic variation and childlessness, particularly for males. We also found that most of the association between damaging genetic variation and childlessness in men could be accounted for by the combination of a lower likelihood of having a partner at home and increased likelihood of never having had sex.

***Do your findings have implications for health? Could they be used to advance medical research?***

There are at least two ways our findings have implications for health and medical research:

1. Our findings suggest that there are important overlaps between damaging genetic variation that causes (or increases risk of) severe neurodevelopmental disorders in patients and damaging genetic variation that has a more subtle effect on cognitive and behavioural traits in the general population. These connections suggest that integrating genetic data on population cohorts, such as UK Biobank, with data on families with neurodevelopmental disorders will improve our ability to identify genes in which damaging genetic variation is causing (or increasing risk of) severe neurodevelopmental disorders. We hope and expect that this will ultimately lead to providing genetic diagnoses for patients that are currently undiagnosed.

2. Involuntary childlessness and an inability to find a partner can have profound psychological effects. We hope that our study will motivate more studies exploring the factors influencing childlessness, especially male childlessness, which has been relatively under-studied compared to female childlessness, and more studies of the relationships between childlessness and mental health.

***Why do you think Darwin's theory of sexual selection is relevant to your results?***

We concluded that the burden of damaging genetic variation is associated with a higher chance of an individual being childless, likely because it primarily makes it harder to find a partner to have children with. We did not find evidence for the burden of damaging genetic variation being associated with the ability of an individual to produce viable sperm or eggs. We came to this conclusion because we found that damaging genetic variation was strongly associated with various cognitive and behavioural traits, and with reporting never having had sex, but not with infertility.

We found that damaging genetic variation is similarly associated in both sexes with the cognitive and behavioural traits that we assessed in this study, but that its correlation with the chance of living with a partner differs between the sexes. This is consistent with the possibility that females might be, on average, more discriminating than males in their choices of partners. In other words, females may be choosier than males: they may be less likely to choose a male partner who has, for example, mental health problems, low income, or less education than a male would be to choose a female partner with the same characteristics. In his work on sexual selection, Darwin proposed this phenomenon of one sex being more discriminating in their choice of partners as 'mate choice'.

There are some traits that are known to be highly valued by both men and women in a prospective partner (e.g. emotional stability) that we could not assess in this study. Thus, we can't exclude the possibility that, for some of these highly valued traits, there is a stronger effect of damaging genetic variation in men than in women, and that this contributes to the differential association of damaging genetic variation with childlessness between the sexes.

***What are the limitations of your study?***

There are several limitations of our study, including the following:

- While our findings are consistent with a role for sexual selection in shaping the gene pool of modern humans, our results are still correlational. While we have controlled for many possible confounders, alternative potential explanations cannot be excluded. Please see our definition of "association, correlation and causation" above for more information.

- UK Biobank participants are, on average, healthier and wealthier than the general UK population[17]. As a consequence, our estimates of the prevalence of damaging genetic variation within UK Biobank may be different from their prevalence within the general UK population. Additionally, we limited our study to UK Biobank participants of European genetic ancestry and thus our results may not be generalizable to other populations.

- We could not study the association of damaging genetic variation with all the cognitive and behavioural traits that have previously been linked to childlessness and reproductive success (e.g. personality traits), because this information is not available for UK Biobank participants.

- Besides genetic data (e.g. Whole Exome Sequencing) provided by the UK Biobank, much of the phenotypic information we use in this study is either self-reported by study participants or provided by Electronic Health Records (EHRs). These two data types can be unreliable due to human or technological limitations. In self-reported data, individuals may misremember events in the distant past (e.g. childhood illnesses) or may omit information they feel is sensitive or personal (e.g. sexual behaviour). EHRs were only widely implemented in the last 30 years in the UK. Considering that UK Biobank participants were 40-70 years old in 2010, some older health records may not have been digitised or were lost altogether. While we have attempted to address these issues by assessing important phenotypes using orthogonal data (e.g. having a partner at home) or using phenotypes that are less likely to be biased (e.g. number of children), it remains possible that our results could be biased by limitations in these data.

- We had no data about the partnerships of UK Biobank participants at the age when people typically have children (on average between 20 - 40 years old), only about whether they said they were living with a partner at the time that they were recruited into the UK Biobank (between 39-73 years old). We assume that people who live with a partner at this older age are also more likely to have lived with a partner at a typical child-bearing age. It is possible that living with a partner at a typical child-bearing age plays a larger role in the association between damaging genetic variation and childlessness than estimated in our study.

- We do not have any direct information about why childless individuals in our study do not have biological children. Thus we cannot distinguish between individuals who are involuntarily childless, and those who choose to be childfree. In the future, surveying UK Biobank participants on this topic could be used to investigate whether the reasons given are different between individuals with a high versus low burden of damaging genetic variation.

#### Social and ethical implications of the study

***Have you found the 'gene for' childlessness?***

No. The phrase 'a gene for' implies a strongly causal, potentially deterministic effect of genetic variation in a single gene on an outcome, here childlessness. This is not what we have found. Rather, we have shown relatively subtle differences in the probability of childlessness between people with a higher versus lower burden of damaging genetic variation. This burden is spread among thousands of genes, and is not focused on any one gene.

Furthermore, genes do not directly shape cognition and reproductive behaviours. Rather, they contribute to how our brains develop and process information from the world, along with important contributions from environmental factors. As a result, both genes and environment influence our behaviour, our cognitive abilities, our personalities, and can influence how others interact with us.

***Can you now predict which individuals will be childless based on their burden of damaging genetic variation?***

No. Similar to what we discuss above in regards to finding "a gene for childlessness", when we and other scientists say that genetic variants (and other factors, such as demographics) "predict" certain outcomes, our use of the word differs in several important ways from how "predict" is used in day-to-day speech. First, we do not mean that the presence of a genetic variant is *guaranteed* to lead to that outcome. Rather, we mean that on average, people with the genetic variant *have a higher chance* of the outcome compared to people without it. A genetic variant is said to be "predictive" of an outcome even if the presence of the genetic variant only very weakly increases the chance of that outcome.

Without any genetic information, the chance that an average participant (both male and female) in UK Biobank will be childless is ~20%. Even for male UK Biobank participants with the very highest burden of damaging genetic variation, this chance only increases to ~50%. The association of damaging genetic variation with childlessness is considerably weaker in females. Thus, this burden of damaging genetic variation does not actually "predict" an individual's likelihood of being childless in the general sense of the word, but it is associated with it.

The genetic factors we have studied here only make a very minor contribution to childlessness on a population scale. We found that $s_{het}$ burden only accounts for a very small fraction (<1%) of childlessness in the general population. Many other factors (especially sociodemographic factors) play a considerably greater role in whether people have children. These include whether they want to have children, and when they were born. For example in the UK, 10% of women born in 1945 remained childless whereas 20% of women born in 1965 remained childless[18,19].

***If the association of these damaging genetic variants with childlessness is so small at a population level, why are you excited about these results?***

We set out to understand why constrained genes are constrained, and not to study the determinants of childlessness. What was exciting to us was that an appreciable proportion (~20%) of why genes are constrained could be explained by the reduced reproductive success associated with increased burden of damaging genetic variants in constrained genes. Therefore, not all of the selection against damaging genetic variation in constrained genes is due to their effects on causing diseases. We were intrigued to discover that this reduced reproductive success was considerably stronger in men, which seems consistent with Darwin's theory of sexual selection by mate choice. If we had not found differences in the association of damaging genetic variation with reproductive success between the sexes, that would not have been compatible with an appreciable role for sexual selection in shaping recent human evolution.

***How are these two results compatible?***

The impact of natural selection over successive generations is a bit like compound interest. In each generation, a damaging genetic variant that, on average, decreases reproductive success is less likely to be passed on to the next generation than variants that have no impact on reproductive success. In each generation, the pool of such damaging genetic variants is reduced further and further. Thus, while damaging genetic variants arise at similar rates in all genes, those that decrease reproductive success are eliminated from the population sooner than other genetic variants. Therefore, when we measure damaging genetic variants across all genes in a population, we find that a subset of genes (constrained genes) have many fewer damaging genetic variants than others. In other words: the size of the reduction in reproductive success in any one generation can be very small, but the effect on which genetic variants survive over many generations can be large.

***Many people feel like having children is a choice. Do your results suggest that this isn't true?***

While it is true that many people in the UK who do not have children chose not to, quite a lot of childless individuals (30% of men, 34% of women) report that they are involuntarily childless - either because they have not have met the right person, or because they or their partner are affected by infertility[9]. This indicates that individual choice is not the only factor. As our genetic findings only explain a very small proportion of the overall variability in the chance of being childless in the population, our findings do not change this situation in any substantive way. While there are some well-documented relationships between specific genes and childlessness[20,21], genetic variation is not the primary cause of involuntary childlessness in the general population; other non-genetic factors are more influential.

***Is there a risk that this research could lead to discrimination?***

As with many studies that make use of genetic data, our findings could unfortunately be misused, misinterpreted, or abused for discrimination. However, we hope that our findings will not be misapplied in that way. We emphasize that the aim of this study is not to identify genetic variants associated with childlessness, but rather to understand the selective pressures that remove damaging genetic variation from the general population.

Here we list three potential misinterpretations of our primary results and why they are incorrect:

- ***Misinterpretation***: An individual's burden of damaging genetic variation determines whether they are going to have children or not, and therefore, individuals with the highest burden of damaging genetic variation are inevitably destined to be childless.
  ***Why this is wrong***: Our results apply at the level of a whole population and over many generations, and very little can be inferred about the likelihood of any specific individual having children.
  ***For more information***: Can you now predict which individuals will be childless?

- ***Misinterpretation***: An individual's burden of damaging genetic variation is more important than free choice and sociodemographic factors in influencing whether an individual has children or not.
  ***Why this is wrong***: While the number of children a person has is partially shaped by genetic variation, genetics only play a minor role compared to non-genetic factors[13]. The type of variation that we studied here, rare damaging genetic variation, contributes very little (<1%) to the overall variation in childlessness in the human population. Free choice and sociodemographic factors contribute more to childlessness.
  ***For more information***: What do we already know about childlessness? and Why are you so excited about these results?

- ***Misinterpretation****:* That there must be a causal role between damaging genetic variation and childlessness.
  ***Why this is wrong***: Although we have observed a correlation between damaging genetic variation and childlessness, this does not necessarily mean that genetic variation causes childlessness. While we have made significant efforts to support our findings and hypotheses, we cannot rule out that other mitigating or confounding factors may be playing a role in the association between damaging genetic variation and childlessness.
  ***For more information***: Association, correlation, and how they differ from causality.

Several steps have been taken to reduce the risk that our findings are not misapplied or misinterpreted. The UK Biobank study applied for and received Research Tissue Bank (RTB) approval from its ethics committee that covers the majority of proposed uses of the Resource, so researchers do not typically need to obtain separate ethics approval (https://www.ukbiobank.ac.uk/learn-more-about-uk-biobank/about-us/ethics). Our project proposal was reviewed and approved by the UK Biobank team (application 44165). We have also communicated our findings to the UK Biobank team. We have invited critical comments from external researchers not involved in this work and beyond those who took part in the peer-review process at *Nature*, including presenting the results of our study at national and international conferences and as a preprint on BioRχiv. Finally, we have prepared this Frequently Asked Questions (FAQ) document. As part of this process, we consulted a diverse range of readers including social scientists and science communication professionals on a wide variety of topics and solicited feedback on the answers. Preparation of such FAQ documents that place the research findings in a wider context in a more accessible way is recognised to be good practice in communicating genetics findings on human behaviour and social outcomes, as pioneered by the Social Science Genetic Association Consortium (https://www.thehastingscenter.org/genomics-research-index/).

#### Supplementary Note 2. Calculation of the contribution of $s_{het}$ to overall fitness

Here we document and provide as a working example our methodology for how we derived the value of 20% for the contribution of $s_{het}$ to fitness as presented in the abstract and main text. This value is based on using the results of the regression analyses to estimate the fertility ratio of individuals with $s_{het} = 0$ and $s_{het} = 1$, as a consequence of the association of $s_{het}$ burden on increased childlessness. This is done separately for males and females, and then averaged (see formula 9 below). Our logistic model for the association of $s_{het}$ burden with childlessness is:

$$has.children \sim s_{het[i.v]} + age + age^2 + birth.cohort + wes.cohort + PC1..PC40 + rare.PC1..PC100 \quad (1.1)$$

The OR derived from this model is generalizable to the formula of:

$$OR = \frac{childless_{shet(0)}/has.child_{shet(0)}}{childless_{shet(1)}/has.child_{shet(1)}} \quad (1.2)$$

where $childless_{shet(0)}$ and $childless_{shet(1)}$ are the proportion of individuals with an $s_{het}$ burden = 0 and 1, respectively, who do not have children and $has.child_{shet(0)}$ and $has.child_{shet(1)}$ are the proportion of individuals with an $s_{het}$ burden = 0 and 1, respectively, who do have children. For males, we know that in the UK Biobank-recruited population $childless_{shet(0)}$ = 20.8% (and thus $has.child_{shet(0)}$ = 79.2%). Additionally, since we are using a proportion we can use the formula:

$$has.child_{shet(1)} = 1 - childless_{shet(1)} \quad (1.3)$$

To further simplify equation (1.2):

$$OR = \frac{childless_{shet(0)}/has.child_{shet(0)}}{childless_{shet(1)}/(1-childless)_{shet(1)}} \quad (1.4)$$

which, with the OR from Supplementary Table 4 and known fertility values for the UK Biobank population as inputs is:

$$0.317 = \frac{0.208/0.792}{childless_{shet(1)}/(1-childless)_{shet(1)}} \quad (1.5)$$

We can solve equation (1.5) for $childless_{shet(1)}$ to obtain the expected proportion of males at $s_{het} = 1$ without children using the formula:

$$childless_{shet(1)} = \frac{0.262}{0.317 + 0.262} \quad (1.6)$$

This calculation gives a value of 0.453. In other terms, we expect that 45.3% of males at $s_{het}$ = 1 will be childless. We next use this value to calculate the expected mean number of children among high $s_{het}$ carriers:

$$mean.expected.children_{shet(1)} = (1 - childless_{shet(1)}) * 2.232 \quad (1.7)$$

where 2.232 represents the mean number of children born to individuals who have children. Equation (1.7) assumes that, in individuals with any children, $s_{het}$ does not have an association with the number of children individuals have, as shown in Extended Data Figure 2. Solving this equation gives a mean expectation of 1.222 children among a sufficiently large population of $s_{het} = 1$ individuals. We can then derive a fertility ratio from this value using the following formula:

$$fertility.ratio = \frac{mean.expected.children_{shet(1)}}{mean.expected.children_{shet(0)}} \quad (1.8)$$

Since we know that the mean number of children born to $s_{het} = 0$ males in the UK Biobank is 1.769 (i.e. $mean.expected.children_{shet(0)}$), equation (1.8) provides a fertility ratio of 0.691. Since $s_{het}$ is calculated from a sex-combined cohort (the Exome Aggregation Consoritum)[2], we also calculate the same value for females, which gives a female fertility ratio of 1.634/1.803, or 0.906. Assuming a 1:1 sex ratio, we then average these two values to derive a mean sex-averaged fitness of 0.798. As this value represents fertility in relation to the unburdened population rather than the reduction in fitness, we then subtract this value from 1:

$$reduction.fitness = 1 - \left(\frac{fertility.ratio_{male} + fertility.ratio_{female}}{2}\right) \quad (1.9)$$

To give an estimate of 20% for the sex-averaged contribution of $s_{het}$ burden on fitness. Ideally, we would have based the values used above for population expectations (e.g. $childless_{shet(o)}$) on estimates from the entire population rather than from the UK Biobank, but such data is not available for the entire UK birth cohort overlapping the same birth years as the UK Biobank. As such, our calculation may underestimate the error on our point estimates. Nonetheless, at a population the size of the UK Biobank we do not expect this issue to be significant.

#### Supplementary Note 3. Calculation of the contribution of fluid intelligence to overall fitness

Here we provide a detailed method for the derivation of our estimate for the individual contribution of cognition to overall fitness as predicted by $s_{het}$ burden, which is generalisable to our similar calculation for mental health disorders. To do this, we use the following equation:

$$contribution_t = \frac{1 - fertility.ratio_t}{1 - fertility.ratio_{s_{het}}} \quad (2.1)$$

The denominator is derived from the calculation performed in Supplementary Note 1 (specifically equation 1.8) but how we generate the numerator for each trait, $t$, is slightly different. Here we provide a worked example of how we determine the numerator of equation (2.1) for fluid intelligence.

When estimating the contribution of fluid intelligence to the reduction in fitness as predicted by $s_{het}$ burden, we first take our estimate of the reduction in fluid intelligence as predicted by $s_{het}$ from the linear model:

$$fluid.intelligence \sim s_{het[i,v]} + age + age^2 + birth.cohort + wes.cohort + PC1..PC40 + rare.PC1..PC100 \quad (2.2)$$

As shown in main text Figure 3F, this model predicts that a male with $s_{het}$ = 1 has a reduction in fluid intelligence of 0.65 standard deviations. As we describe in the main text Methods, we next utilize population-level data from Sweden with paired IQ-fertility data on males (Supplementary Table 5)[12]. As the relationship between fertility and IQ from those data is an empirical distribution, we used simulations to model a "population" of $s_{het}$ = 1 males with the IQ distribution of this "population" shifted by the estimate derived from equation (2.2). To translate a reduction in fluid intelligence to a reduction in IQ, we used the formula:

$$\Delta_{IQ} = \beta_{fluid.intel} * \sigma_{IQ} \quad (2.3)$$

This formula, when solved, gives a reduction of 7.54 IQ points for a male with $s_{het}$ = 1.

We then simulated the IQ score distribution of $1\times10^6$ males with $s_{het}$ = 1 based on a normal distribution with μ = 92.46 (i.e. 100 - 7.54) and σ = 12. These "individuals" were then assigned a number of children based on the empirical Swedish distribution (lookup table provided in Supplementary Table 5). We then compared this overall fertility to a simulation of males with $s_{het}$ = 0 (i.e. the unburdened population with a distribution μ = 100 and σ = 12) to generate a fertility ratio:

$$fertility.ratio_{fluid.intelligence} = \frac{fertility_{shet(1)}}{fertility_{shet(0)}} \quad (2.4)$$

where fertility for both the numerator and denominator are the average number of children in $1x10^6$ simulated individuals from the affected and unaffected distributions, respectively. For males, the numerator and denominator in equation (2.4) are 1.71 and 1.77, respectively. Solving this formula thus gives a value of 0.967. When subtracted from 1 as in equation (2.1), this value represents the reduction in fitness attributable to a decrease in IQ caused by $s_{het}$ = 1. We then substitute this value as the numerator in equation (2.1) and divide this value by the overall reduction in male fitness caused by $s_{het}$ = 1 as calculated in equation (1.8):

$$contribution_{fluid.intelligence} = \frac{1-0.967}{1-0.691} = \frac{0.033}{0.307} = 10.5\% \quad (2.5)$$

In other words, for males, we expect IQ to contribute ~10% of the observed association of $s_{het}$ with fertility (note that rounding errors in numbers displayed in equation (1.8) result in slightly different values). This calculation was also performed at various $s_{het}$ values and is shown in Supplementary Figure 19.
